# Metal-Binding Ligands Rather Than Redox Active Metabolites Are Essential to Microbially-Induced Corrosion of Cobalt

**DOI:** 10.1101/2025.06.04.657868

**Authors:** Annika DeJager, Anna Kirkland, Aaron Hinz, David R. McMullin, Daniel S. Grégoire

**Author notes:** Address correspondence to Daniel S. Grégoire.

## Abstract

Microbially-induced corrosion (MIC) has been well-studied in the context of damage to iron-bearing infrastructure, where microbes can solubilize solid elemental iron using redox-active metabolites such as phenazines. Whether such pathways can solubilize economically-critical cobalt remains poorly understood. We hypothesized that secondary metabolites produced by the model bacterium *Pseudomonas chlororaphis* subspecies *aureofaciens* associated with MIC of iron could corrode cobalt by oxidation (*i.e.,* phenazines) or by acting as a ligand (*i.e.,* cyanide). One-week incubations using live cells supplied with Co(0) wires led to 20-30% cobalt mass losses and ∼2200 µM Co(2+) recovered in solution, which was at least 3-fold higher than filtered cultures and sterile medium controls. Removing the capacity for cells to produce phenazines and cyanide showed similar corrosion compared to wild type cells and phenazine standards did not corrode cobalt, ruling these metabolites out as contributors to MIC. Further experiments testing whether metabolite mixtures devoid of phenazines could corrode Co(0) in the presence and absence of oxygen confirmed that atmospheric oxygen initiates Co(0) oxidation, and that unidentified cell-derived metabolites drive MIC forward by keeping Co(2+) from precipitating as oxides that form a passivation layer on the wire surface. This revised mechanistic explanation underscores the importance of considering how abiotic redox cycling and cellular metabolites that solubilize metals interact in the MIC of non-ferrous metals. Identifying the biosynthetic pathways involved in solubilizing cobalt will be key to reframing MIC as more than an environmental concern and optimizing sustainable cobalt recovery strategies for solid waste that rely on naturally-occurring microbial metabolites.

**Importance:** Characterizing microbially-induced corrosion mechanisms for cobalt is important for predicting the fate of critical metals in the environment and optimizing sustainable strategies for cobalt reclamation from electronic waste. This study uses the model bacterium *Pseudomonas chlororaphis* subspecies *aureofaciens* to test if redox-active phenazines and the metal-binding metabolite cyanide can corrode elemental cobalt. We repeatedly observed that live cells corroded cobalt, but that corrosion did not rely on phenazines or cyanide. Instead, our results show that oxygen oxidized elemental cobalt and unidentified metabolites keep cobalt ions in solution to limit cobalt oxide precipitation and drive the corrosion reaction forward. Our study reinforces the need to consider connections between abiotic and biotic reactions in the corrosion of metals other than iron. Understanding these mechanisms is critical to reframing microbial corrosion as a sustainable approach for metal recovery and our study makes the case that it is possible for the critical metal cobalt.

## Introduction

Microbially-induced corrosion (MIC) due to biofilm formation is estimated to damage metal-bearing infrastructure at a cost on the order of trillions of USD per year (1–5). MIC occurs when bacteria or archaea oxidize solid elemental metals into soluble ions (1). Diverse metabolic guilds that thrive along the redox gradients within biofilms contribute to MIC (1). These guilds include anaerobic heterotrophic bacteria, nitrate-reducing bacteria, iron-reducing bacteria, sulphate-reducing bacteria, acetogenic bacteria, and methanogenic archaea (1, 6).

MIC mechanisms described in model organisms from these guilds rely on direct electron transfer (DET) or mediated electron transfer (MET) between cells and metal surfaces (6–8). Multiple species can corrode metals through both types of pathways depending on growth conditions and reaction kinetics (1, 9–12). DET has been observed in model iron-reducing bacteria and methanogenic archaea that couple Fe(0) oxidation to energy production through direct contact between metals and multi-heme outer surface *c*-cytochromes or electroconductive pili (8–14). MET can be facilitated by hydrogenase-mediated H_2_ oxidation or organic acid deprotonation, which produce protons that can accept electrons from metal surfaces, promote corrosion, and regenerate H_2_ (1, 15–21). MET can also occur when redox-active metabolites (*e.g.,* flavins and phenazines) oxidize metals (1, 22–30), given their capacity to act as exogenous electron acceptors that typically shuttle electrons in and out of cells to maintain redox balance (31, 32).

The current mechanistic understanding of MIC primarily stems from studies on iron and stainless steel (1, 2, 6–8). Although the focus on iron-based materials is understandable given concerns surrounding damage to infrastructure, MIC mechanisms that affect the chemistry of other metals remain understudied. MIC has the potential to mobilize elemental metals into nearby environments through oxidation, given evidence that MET within biofilms accelerates the corrosion of non-ferrous metals (33–44). Previous work has demonstrated that MIC of Al(0) alloys occurs through H_2_ and organic acid-mediated MET pathways (33–35). H_2_S, oxalic acid, and NH_3_ have all been linked to the MIC of Cu(0) (36–38). Riboflavin and phenazines have also been associated with the corrosion of Ni(0) and Ti(0), respectively (39–44).

Despite MET pathways being widespread in metabolically versatile guilds, whether such pathways support the MIC of other economically, environmentally, and physiologically relevant metals remains poorly understood. Characterizing these pathways will advance our understanding of how MIC controls the mobility and toxicity of elemental metals that are prevalent in habitats contaminated with electronic and municipal solid waste (45, 46). Assessing if MIC can solubilize metals critical to the global economy could open new possibilities for sustainable metal recovery strategies, reframing MIC as more than an undesirable process that damages infrastructure.

This study tests if phenazines produced by the model bacterium *Pseudomonas chlororaphis* subspecies *aureofaciens* can corrode elemental cobalt. We focus on cobalt because it is an essential component of consumer electronics that is a priority for reclaiming from electronic waste because it is mined at great environmental expense (45–47). We selected *P. chlororaphis* subsp. *aureofaciens* because it is a genetically tractable non-pathogenic strain that produces carboxylated and hydroxylated phenazines (48–50). These metabolites are similar to pyocyanin (PYO) and phenazine-1-carboxamide (PCN) produced by the model organism *Pseudomonas aeruginosa*, which can corrode Fe, Ti, and Cr by MET (26–30). Lastly, we chose to work with this strain and cobalt because Co(0) oxidation to Co(2+) (E^0^_red_ = −0.282 V) by phenazine-1-carboxylic acid (PCA) (E^0^_red_ = −0.116 V), the main phenazine produced by *P. chlororaphis* subsp*. aureofaciens*, is predicted to be thermodynamically favourable (ΔG^0^ = -nFE^0^_cell_ = −32.04 kJ/mol) (49–52).

To test this mechanistic hypothesis, we compare the corrosion of Co(0) wires in the presence of wild-type cells to corrosion by mutant cells devoid of the *phz* operon required for phenazine production. We assess whether PCA can directly corrode Co(0) or if corrosion depends on complex mixtures of other metabolites produced by cells. We consider whether the capacity for cells to produce cyanide, which can form soluble complexes with cobalt (53), can also solubilize Co(0). Multiple independent analytical techniques including ICP-MS, LC-HRMS, and SEM-EDX confirmed that live cells corrode cobalt through pathways that do not rely on phenazines or cyanide production. Instead, this study shows that unidentified metal bindings ligands produced by cells keep cobalt in solution following oxidation by atmospheric oxygen. We consider how this alternative explanation reshapes our view of MIC mechanisms including existing paradigms of MET pathways involved in iron corrosion.

## Results & Discussion

### Live cells are required for cobalt corrosion, but not phenazines

Cobalt corrosion was confirmed by measuring aqueous Co(2+) concentrations released from solid Co(0) wires over 168 hours using ICP-MS. Aqueous Co(2+) concentrations were used to corroborate mass losses from the Co(0) wires in mass balances. To ensure representative comparisons between treatments, mass balance recoveries from each individual flask were used to correct Co(2+) concentrations in all subsequent analyses. Mean cobalt recoveries for all experiments ranged from 93.17 ± 4.66 % and 125.45 ± 3.57 % with an average across all experiment of 99.13 ± 2.80 %, indicating that all cobalt in the system was accounted for (**Table S1**). Deviations from 100 % recovery can be attributed to analytical variance on the ICP-MS, having ruled out the effects of evapoconcentration through water mass balances in line with expectations (91.62 ± 3.11 % to 99.94 ± 0.22 %, avg. 96.67 ± 2.16 %) (**Table S1**).

Live *P*. *chlororaphis* subsp. *aureofaciens* cells corroded more cobalt than sterile medium treatments after 168-hour incubations. Mean Co(2+) concentrations for the wild-type and Δ*phz* strains increased steadily over 72 hours before reaching a plateau at 2188.17 ± 730.91 µM and 2139.84 ± 308.55 µM, respectively, by 168 hours (**Figure 1**). By comparison, abiotic treatments in sterile medium had mean Co(2+) concentrations that plateaued after 24 hours, with 240.47 ± 76.45 µM recovered after 168 hours (**Figure 1**). This observation suggests that abiotic reactions in the medium accounted for ∼6 to 11 % of the corrosion observed with live cells.

**Figure 1:**
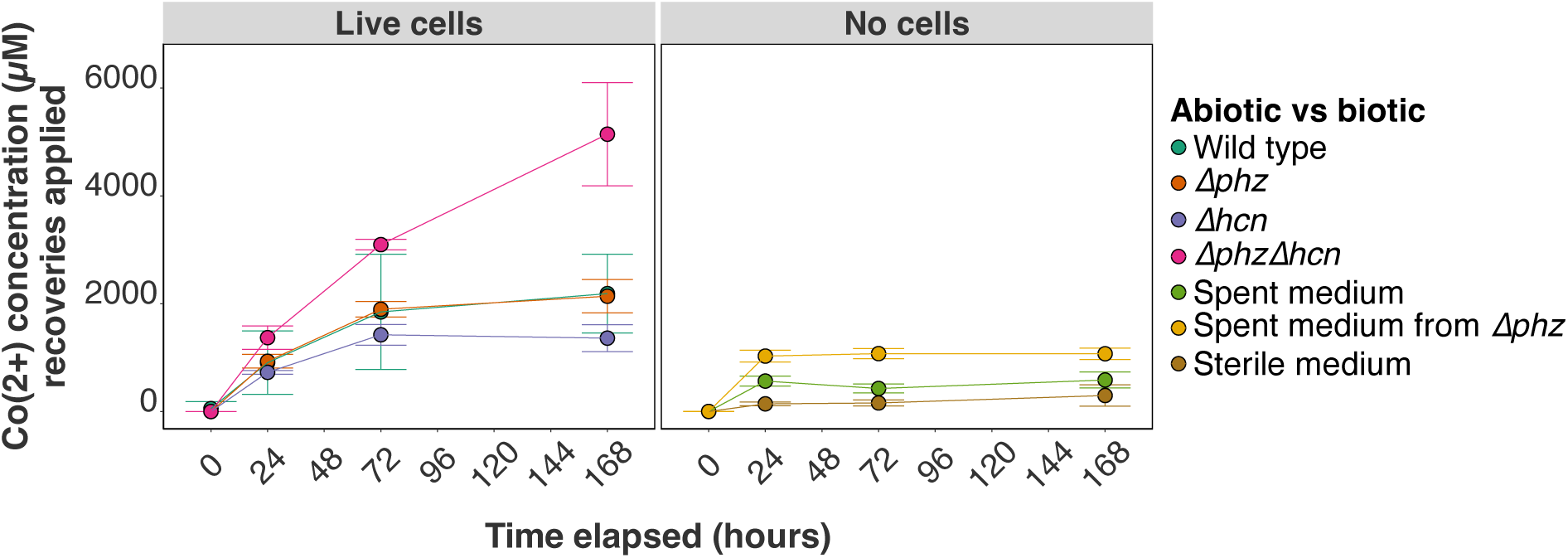
Mean Co(2+) concentrations ± SD obtained by ICP-MS from biotic and abiotic treatments during 168-hour incubations, corrected for mass balance recoveries. Replicates used for mean calculations were Wild type: n = 6, Δ*phz*: n = 4, Δ*hcn*: n = 4, Δ*phz*Δ*hcn*: n = 4, Spent medium: n = 4, Spent medium from Δ*phz*: n = 3, and Sterile medium: n = 9.

Differences observed in live cell vs abiotic treatments are not attributable to variance in background concentrations of Co(2+) in different batches of media, which were negligible relative to Co(2+) concentrations recovered from corrosion assays with wires (*i.e.,* 1.69 to 3.39 µM, **Figure S1**). The trends observed for aqueous Co(2+) measured in biotic treatments are supported by wire mass losses. Wires incubated with wild-type and Δ*phz* live cells had mean relative losses of 26.22 ± 12.02 % and 34.11 ± 2.65 %, which were 3 to 10-fold greater than the losses recorded for abiotic controls (**Table S1**). Statistical comparisons of Co(2+) concentrations at 168 hours confirmed that live cells solubilized significantly (p < 0.05) more Co(0) than abiotic treatments (**Figure S2**).

We can confidently rule out that phenazines contribute to cobalt corrosion since no significant difference was observed in Co(2+) recovered from the Δ*phz* treatment compared to the wild type (**Figure 1 & Figure S2**). Furthermore, we confirmed that Δ*phz* mutants lacked the genetic basis for phenazine production via long-read sequencing (**Methods**) and phenotypic tests measuring phenazine production. Total phenazine concentrations for wild-type cells in corrosion experiments ranged from 5.33 µg/mL to 9.48 µg/mL, which was largely comprised of PCA and minor amounts of 2-hydroxyphenazine (2-OHPHZ) and 2-hydroxyphenazine-1-carboxylic acid (2-OHPCA) (**Figure 2**). In contrast, we could not detect any phenazines produced in the Δ*phz* mutant by LC-HRMS (**Figure 2**). These observations align with previous reports measuring phenazines in this strain and close relatives, showing that the analytical techniques were appropriate for confirming the absence of phenazines in Δ*phz* mutants (49, 50, 54, 55). As an additional independent test of our mechanistic hypothesis, we carried out abiotic experiments with a PCA standard at 5-fold higher concentrations than what we recovered from wild-type cultures. These experiments confirmed that even when PCA was incubated in the presence of oxygen, the Co(2+) recovered at 168 hours was indistinguishable from sterile medium with no amendments (**Figure S3**). These analyses provided multiple lines of evidence pointing to alternative mechanistic hypotheses for cobalt corrosion that needed to be tested.

**Figure 2:**
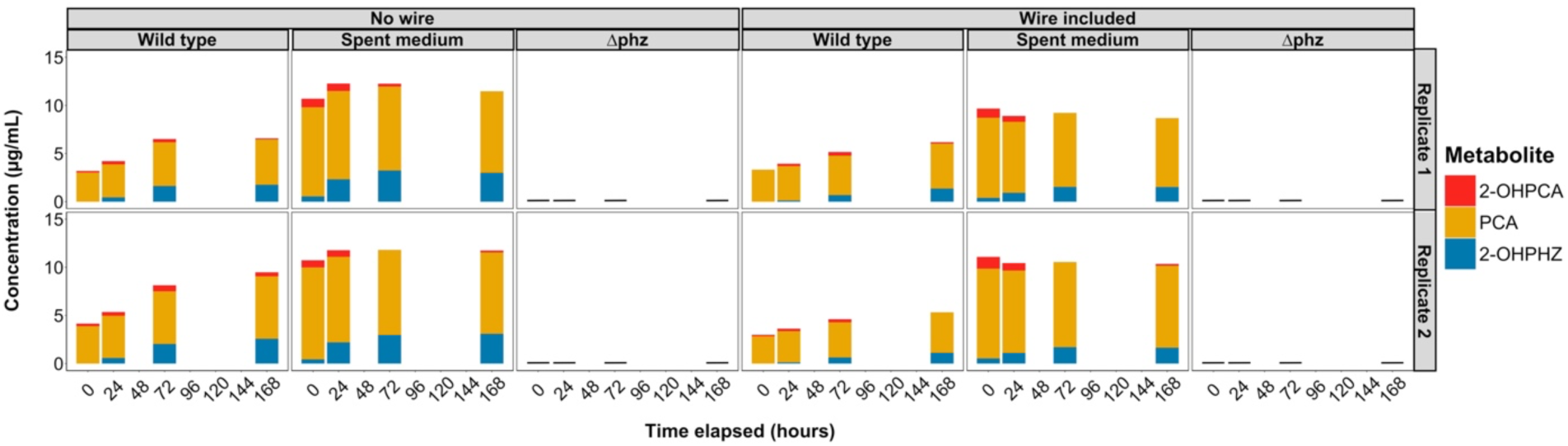
Total phenazine concentration recovered from representative duplicate wild type, Δ*phz*, and spent medium incubations in the presence and absence of Co(0) wires over 168 hours. Stacked bars show contributions of phenazine-1-carboxylic acid (PCA), 2-hydroxyphenazine (2-OHPHZ), and 2-hydroxyphenazine-1-carboxylic acid (2-OHPCA) to the total phenazine amount at each time-point. Black bars are shown for the Δ*phz* experiments to denote that phenazines were not detected at these specific time points.

### Cell-derived ligands other than cyanide are critical to cobalt corrosion

Having ruled out phenazines as oxidants and solubilizing agents for Co(0), we looked to the spent medium treatments for additional insights into the physiological controls for the MIC of cobalt. When incubating Co(0) wires with spent medium from filtered wild-type cultures containing a mixture of metabolites, we observed intermediate corrosion in between what was seen for live cells and sterile medium treatments (**Figure 1 & Figure S2**). Spent medium from wild-type cells showed plateaus in Co(2+) concentrations at 24 hours with a final concentration of 586.44 ± 147.91 µM recovered by 168 hours, which aligned with an average wire mass loss of 10.02 ± 2.83 % (**Figure 1 & Table S1**). To further confirm phenazines were not solubilizing Co(0), additional experiments were run using spent medium from Δ*phz* mutants. Interestingly, spent medium from Δ*phz* cells achieved higher Co(2+) concentrations compared to spent medium from wild-type cells at 168 hours of 1073.20 ± 105.79 µM Co(2+) and 14.86 ± 1.97 % wire mass loss (**Figure 1 & Table S1**). Although this difference was marginally not statistically significant (*i.e.,* p-value of 0.08), this observation suggests that removing the capacity for phenazine production had a slight stimulatory effect on cobalt corrosion that was not apparent in the Δ*phz* live cell experiments (**Figure S2**). The intermediate corrosion observed in spent medium compared to sterile and live cell treatments suggests that cells can potentially recycle metabolites responsible for cobalt solubilization, leading to increased Co(2+) released from wires over time (**Figure 1**).

To better understand what potential metabolites could be solubilizing cobalt, we carried out biosynthetic gene cluster analyses (discussed in detail in **Supplementary Information)**. Genome annotations generated with antiSMASH showed that *P. chlororaphis* subsp. *aureofaciens* carries the *hcn* genes required for cyanide production (**Figure S4 & Table S2**). Biogenic cyanide production has been associated with gold, platinum, and cobalt solubilization with the CN^-^ ion acting as a ligand to keep metal ions in solution (53, 56, 57). Typically, glycine is supplied as a carbon and nitrogen source to stimulate cyanide production in Pseudomonads (53). Although we did not provide glycine to cells, independent annotations with DRAM confirmed the presence of accessory genes for hydrogen cyanide synthesis [*e.g.,* sarcosine (*N*-methyl glycine) oxidase subunit] that could supply glycine via amino acid metabolism (58) (**Table S2**). Based on the detection of these genes and cyanide’s precedence in forming stable metal complexes with Co(2+) ions in solution, we developed a Δ*hcn* deletion mutant to test an alternative mechanistic hypothesis for pathways that could contribute to cobalt solubilization.

Live cell experiments with Δ*hcn* mutants led to similar trends in Co(2+) release over 72 hours compared to wild-type cells, achieving a maximum of 1362.93 ± 249.18 µM Co(2+) by 168 hours (**Figure 1**). Preclusion of cyanide production did not result in statistically significant differences in Co(2+) recovered at 168 hours when compared to wild-type and Δ*phz* live cell treatments (**Figure S2**). However, the slightly lower Co(2+) concentrations associated with Δ*hcn* mutants did not statistically differ when compared to spent medium from Δ*phz* cells (**Figure S2**). The trend observed in Δ*hcn* mutants and spent medium from Δ*phz* cells suggests that the availability of common metabolic substrates involved in phenazine and cyanide synthesis can affect cobalt corrosion, which warrants further investigation.

To further test for connections or compensatory effects between phenazine and cyanide biosynthesis, cobalt corrosion assays were carried out with a double deletion mutant devoid of both Δ*phz* and Δ*hcn* operons. Interestingly, the Δ*phz*Δ*hcn* mutant achieved the highest levels of cobalt corrosion among all live cell experiments with mean wire mass losses of 51.44 ± 5.55 % (**Table S1**). Mean Co(2+) concentrations never plateaued like the other live cells treatments, achieving a maximum of 5144.30 ± 956.90 µM by 168 hours, which was significantly higher than all other experimental treatments (**Figure 1 & Figure S2**).

These independent tests reinforce that phenazines are not solubilizing Co(0), nor is the well-known metal-binding ligand cyanide. The increase in Co(0) corrosion observed for Δ*phz*Δ*hcn* mutant suggests that the synthesis of an unidentified metabolite capable of solubilizing cobalt likely relies on the same metabolic substrates (*e.g.,* carbon and nitrogen) involved in the production of phenazines and cyanide. This explanation is supported by previous transcriptomic studies with cyanide-deficient *Pseudomonas chlororaphis* R47 that showed how the removal of certain biosynthetic pathways promoted the redirection of carbon and nitrogen resources towards the synthesis of ligands or oxidants that could solubilize cobalt (59).

Our biosynthetic gene cluster analyses showed several putative metal-binding metabolites other than cyanide that could affect cobalt solubility (**Figure S4**). The core genes we identified in cluster 6 show moderate homology (∼46-53 %) to genes involved in the production of schizokinen and vibrioferrin (**Table S3**). Schizokinen is a known siderophore produced by cyanobacteria (60) and bacteria (61), while vibrioferrin plays a similar role in iron acquisition by *Vibrio alginolyticus* (62). When annotated with DRAM (**Section 2** in **Supplementary Information**), these genes were linked to staphyloferrin production, a siderophore produced by members of the *Staphylococcus* genus (63) (**Table S3**). Genes in cluster 10 were predicted to synthesize different types of pyoverdines including Pf-5 pyoverdine and pyoverdine SXM-1 (**Table S3**). Pyoverdine synthesis genes in cluster 10 occurred alongside genes involved in siderophore transport, suggesting a potential pathway that could affect cobalt uptake (**Table S2**). Similarly, genes in cluster 11 (encoding for a non-ribosomal peptide synthase) were also predicted to support Pf-5 pyoverdine production in addition to azotobactin-D, a siderophore originally isolated from *Azotobacter vinelandii* strain D (64). Notably, pyoverdines are well-characterized metallophores involved in metal uptake in *P. aeruginosa* and are known to form complexes with cobalt (65, 66).

These observations show that biogenic cyanide is not involved in cobalt solubilization. These observations also highlight that multiple other metabolites that rely on carbon and nitrogen for synthesis potentially contribute to corrosion when cells lack the capacity for phenazine and cyanide production. Our biosynthetic gene cluster analyses revealed that multiple genes, many of which are not well-characterized in *P. chlororaphis* subspecies *aureofaciens*, are likely involved in the synthesis of these metabolites. Given this complexity and potentially unknown phenotypic effects, targeted deletions like those carried out for the well-characterized *phz* and *hcn* operons are not practical for identifying which metabolite(s) solubilize cobalt. Instead, we plan to carry out transcriptomic assays in future work to compare gene expression profiles in wild-type cells to Δ*phz*Δ*hcn* mutants during corrosion assays. These future experiments will provide mechanistic insights into whether one or several of the genes involved in metal-ligand synthesis are upregulated during cobalt corrosion. Importantly, such approaches will allow us to account for the complex interplay that likely exists between biosynthetic pathways in this strain.

### Cobalt bioavailability controls corrosion

Having excluded phenazine and cyanide production as contributors to cobalt corrosion, we considered how cell viability and Co(2+) uptake could provide additional insights into the physiological mechanisms controlling MIC of cobalt. We can rule out that differences in cobalt solubilization observed in live cell treatments were due to differences in cell viability between the mutants, which displayed identical growth in the absence of Co(0) wires (**Figure 3**). Likewise, all mutants experienced similar decreases in viable cell density from ∼10^8^ to ∼10^6^ CFU/mL during incubation with Co(0) wires (**Figure 3**). The decreases in cell density shows that cells grown in the presence of Co(0) wires experience cobalt toxicity due to Co(2+) exposure.

**Figure 3:**
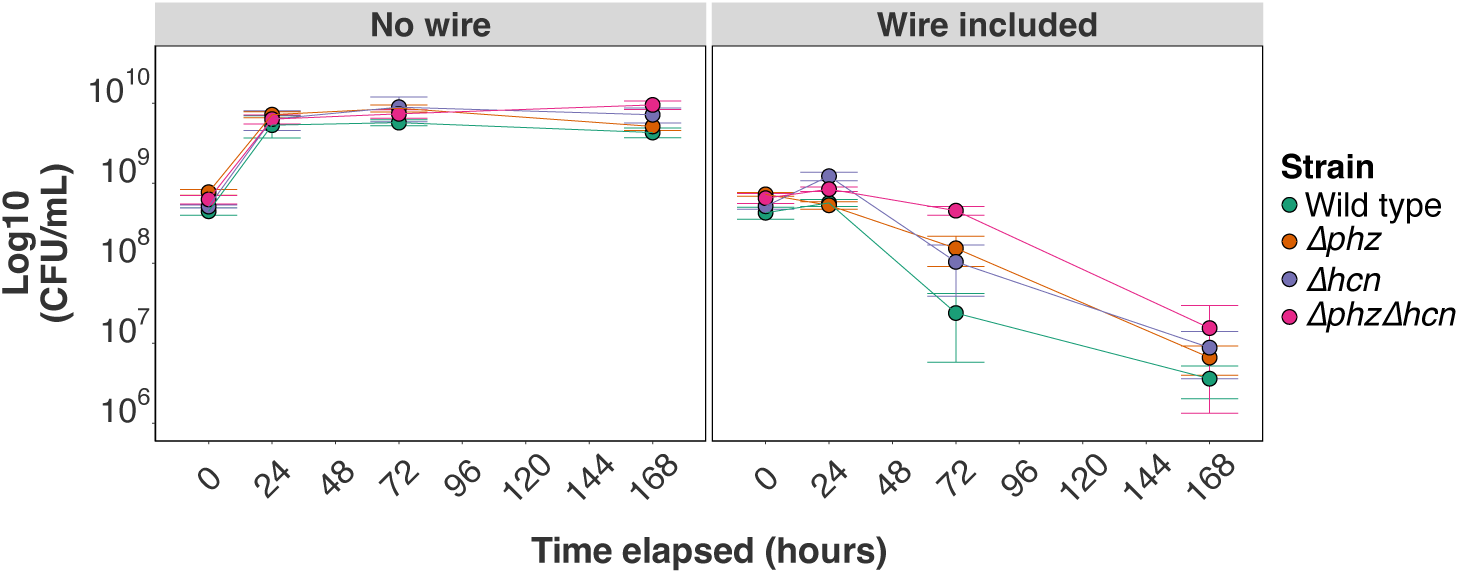
Mean live cell counts of wild type and Δ*phz* cultures in log10(CFU/mL) ± SD over 168-hour corrosion assays. Wild type: n = 4, Δ*phz*: n = 4, Δ*hcn*: n = 4, Δ*phz*Δ*hcn*: n = 4.

Additional tests were conducted to confirm that toxicity stemmed from cells adsorbing and taking up cobalt. ICP-MS analyses comparing samples acidified to lyse wild-type cells before filtering to samples filtered to remove biomass prior to acidification showed that 6.64-21.36 % (or 77.04 to 585.63 µM) of Co(2+) was cell-associated (**Figure S5**). This range of concentrations aligns with dose-response experiments demonstrating a 50 % decrease in growth rate at exposure to 23.9 µM Co(2+) (**Figure S6**) and previous reports where 500 µM Co(2+) inhibited the growth of *P. aureofaciens* BS1393 in minimal medium (67). Notably, the capacity for phenazine production, which can be used to mitigate oxidative stress (31), did not lead to improved cobalt tolerance in wild-type cells vs *Δphz* mutants. Based on genome annotations, we cannot rule out the contributions of other pathways such as the disruption of metal homeostasis, metal binding to thiol groups, and the consumption of ATP to support metal efflux as contributing to cobalt toxicity (68–70) (**Supporting Data 1**). Understanding how different modes of toxicity affect the kinetics of cobalt corrosion will be important to consider in future studies to optimize the use of MIC in biological cobalt recovery.

When we carried out the same comparison of sample preservation techniques for sterile medium incubated with a Co(0) wire, we noted greater relative Co(2+) losses when samples were filtered first when compared to corresponding samples from live cells (*i.e.,* 58.9-76.5%, **Figure S7**). We suspected this difference was attributable to the abiotic formation of cobalt oxides during storage under normal atmosphere in the absence metabolites that could keep Co(2+) in solution. This observation prompted us to consider how the precipitation of cobalt oxides might hinder MIC through the formation of a passivation layer, as seen in previous MIC studies (1,4,26–29,35,38,43). Moreover, it made us consider whether atmospheric oxygen was the main oxidant for Co(0) and if cell-derived ligands favoured Co(0) solubilization by precluding cobalt oxide formation.

To formally test this hypothesis, we examined the extent of oxide formation in our original experimental treatments using scanning electron microscopy coupled to energy dispersive x-ray spectroscopy (SEM-EDX). We also carried out corrosion assays under oxic and anoxic conditions with sterile medium and spent medium obtained from filtered Δ*phz* cells to assess whether cell-derived metabolites other than phenazines could oxidize cobalt without atmospheric oxygen as an oxidant.

SEM-EDX analyses of wires in sterile medium alone and sterile medium supplemented with PCA showed uniform dissolution of cobalt and oxide formation (depicted by consistently shaded regions on the wire surface) likely driven by abiotic oxidation but were similar in appearance to the unmanipulated wire (**Figure 4A-C**). Spent medium from filtered wild-type and Δ*phz* cultures produced slight changes in wire surface topography and uniform loss of cobalt, as would be expected if more Co(0) was being solubilized by metabolites that were not present in sterile controls (**Figure 4D, E**). In contrast, wild-type and Δ*phz* cells cored out large lengthwise strips on the wires (**Figure 4F, G**), corresponding to respective decreases in wire diameters by 12.00 % and 16.30 % relative to the sterile wire (**Table S4**). The extent of corrosion observed for the Δ*hcn* treatment was less pronounced than other biotic treatments (**Figure 4H**) and resembled the wire incubated in spent medium from Δ*phz* cells, in line with our previous aqueous Co(2+) analyses (**Figure S2**). The wire incubated with Δ*phzΔhcn* cells produced the most visibly apparent corrosion, with patchy recessions and a curved indentation leading to a relative decrease of 28.31-33.51 % in diameter compared to the sterile medium wire (**Figure 4I & Table S4**). Although corrosion patterns varied considerably between live cell replicates based on imaging (all images have been uploaded to the repository: https://zenodo.org/records/17408329, see **Data Availability Statement**), these observations align with heterogenous pitting appearance seen in previous MIC studies where *P. aeruginosa* biofilms corroded Fe and Ti stamps (26–30, 43, 44).

**Figure 4:**
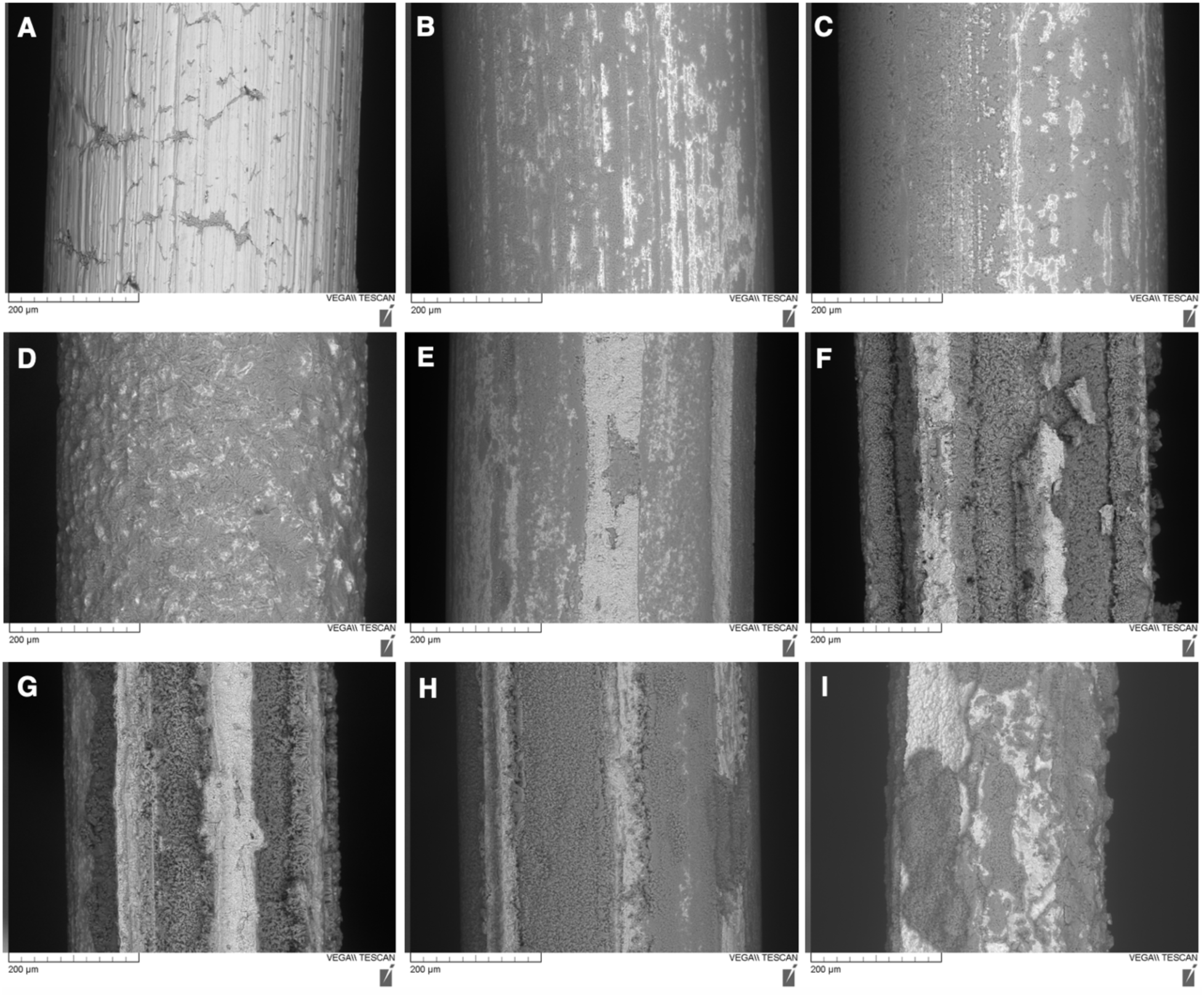
SEM images of Co(0) wires across all treatments after 168 hours. Wires were magnified to 400X and images obtained using the backscattered electron (BSE) detector are displayed. Regions of lighter colour represent greater Co(0) content compared to shaded regions indicative of oxidized areas on the wire. A) Unmanipulated wire. B) Sterile medium. C) PCA. D) Spent medium. E) Spent medium from Δ*phz*. F) Wild type. G) Δ*phz*. H) Δ*hcn*. I) Δ*phzΔhcn*.

EDX scans comparing the lighter to darker regions of the wires, indicative of higher and lower Co(0) content based on respective amount of electrons that were backscattered, supported that Co(2+) released during abiotic and biotic corrosion formed cobalt oxides at the wire surface. Co:O ratios were calculated from percent weights of cobalt and oxygen of darker areas on the representative wires (**Figures S8-S18**) and were compared to those from lighter areas. Unlike lighter regions corresponding to ratios of >1:1, Co:O ratios in darker areas were estimated at ∼1:1 or ∼2:3 when rounding up to whole integers (**Table S5**). Although this EDX data is semi-quantitative, it suggests that cobalt oxides could form including cobalt(II) oxide (CoO), cobalt(II) hydroxide (Co(OH)_2_), and cobalt(II/III) oxide (Co_3_O_4_) with cobalt in its respective 2+ and 3+ oxidation states (71). Pink precipitate was observed in corrosion flasks, providing qualitative support for cobalt hydroxide in solution with Co(2+) solubilized from the corroding wire (72).

The comparison of corrosion under oxic and anoxic conditions supports the explanation that cobalt oxide precipitation forms a passivation layer that inhibits corrosion after 24 hours in abiotic treatments (**Figure 1 & Figures S9, S10, S13, S14**). When removing oxygen from the treatment with spent medium from filtered Δ*phz* cells, cobalt oxidation was strongly inhibited. The mean Co(2+) concentration of 82.45 ± 30.01 µM at 168 hours for anoxic spent medium was significantly lower than its oxic counterpart, from which 1073.20 ± 105.79 µM Co(2+) was recovered (**Figure S19**). Wires in the anoxic spent medium treatment lost on average 0.69 ± 1.33% of their initial mass (**Table S1**), demonstrating no notable decrease in diameter and minimal formation of cobalt oxides from EDX data (**Figure S20 & Table S5**). Notably, the mean Co(2+) concentrations at 168 hours for anoxic sterile medium (83.59 ± 21.01 µM) were also significantly lower than its oxic counterpart (240.47 ± 76.45 µM) (**Figure S19**). Both lines of evidence support that oxygen is the oxidant for Co(0), and that limiting the precipitation of cobalt oxides is essential for the continued solubilization of Co(0). This mechanistic explanation is further supported by earlier iterations of corrosion assays, where DMSO was employed as a solvent to solubilize phenazines. The addition of DMSO significantly decreased Co(2+) concentrations recovered at 168 hours likely due to the inadvertent formation of large crystalline oxides on the wire surface (**Figure S21 & Figure S22**). Taken with our previous observations, these results suggest that cells and metabolites likely bind to Co(2+) to keep cobalt in solution and preclude the formation of a passivation layer that limits oxygen’s capacity to oxidize and solubilize Co(0).

### Conclusion

Our study provides multiple lines of evidence that *P. chlororaphis* subsp. *aureofaciens* can corrode elemental cobalt through pathways that do not rely on phenazines or cyanide, which are summarized in **Figure 5**. Cobalt corrosion relies on Co(0) being oxidized by oxygen (**Figure 5A**) and cell-derived metabolites facilitate corrosion by preventing Co(2+) from precipitating as oxides (**Figure 5B**). Live cells can potentially promote corrosion by directly taking up Co(2+), possibly through passive or active transporters involved in cobalt homeostasis (*e.g.,* cobaltochelatase encoded by *cobN*) (73, 74) (**Figure 5C**) or facilitated transport of Co(2+) bound to ligands (**Figure 5B**). We note that formal experiments assessing the contributions of different uptake pathways are required in future work. These processes allow atmospheric oxygen to continue oxidizing Co(0), driving the corrosion reaction until physiological limitations preclude keeping Co(2+) in solution.

**Figure 5:**
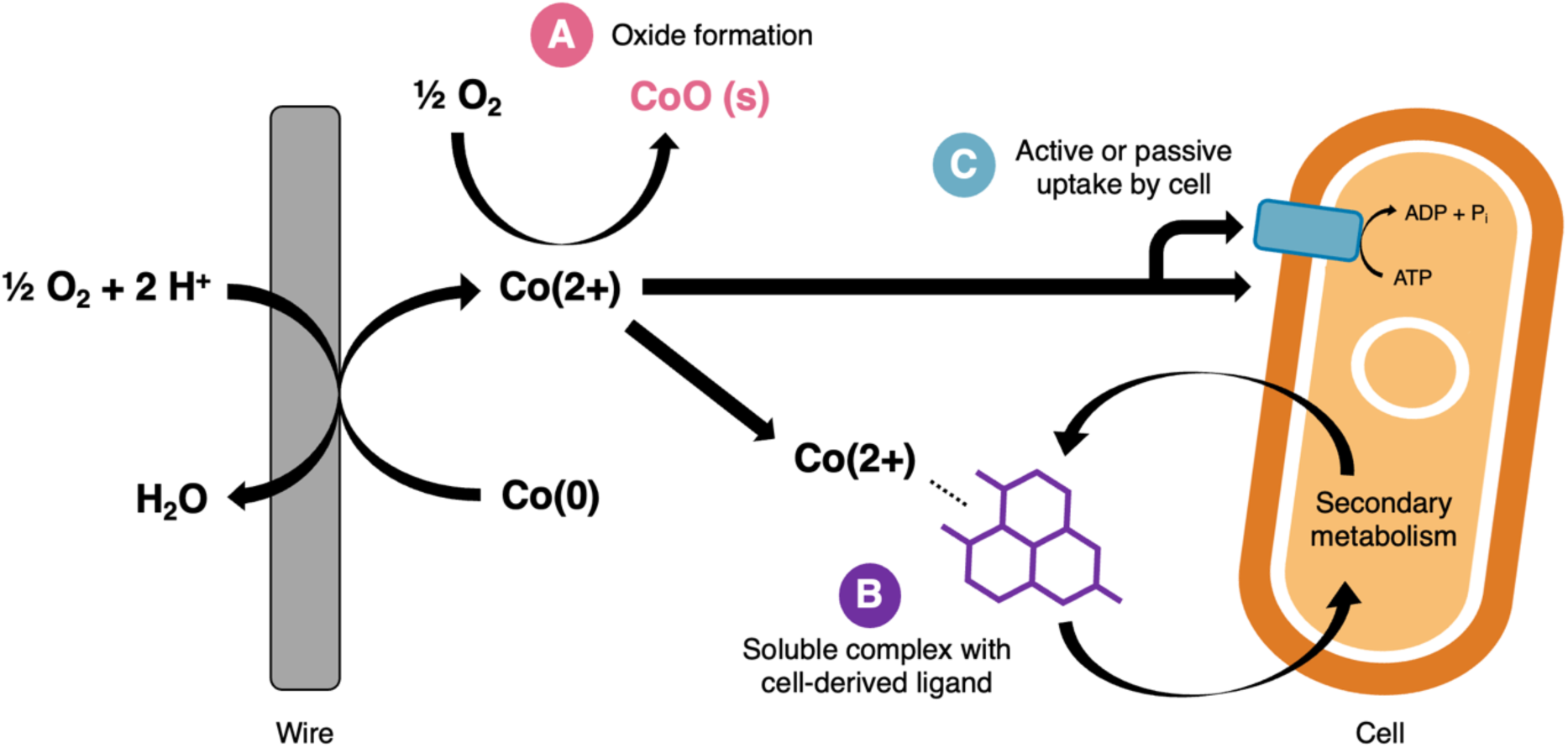
Conceptual diagram illustrating the alternative mechanistic hypothesis for abiotic and biotic processes contributing to cobalt corrosion in this study. Mechanistic details obtained from corrosion assays are summarized as follows: A) Abiotic formation of cobalt oxides from Co(2+). B) Complexation of Co(2+) with microbially-synthesized metabolites that can be taken back up into the cell. C) Direct Co(2+) uptake by cells through active or passive transport mechanisms.

We excluded DET pathways in our revised mechanistic explanation for the MIC of cobalt by *P. chlororaphis* subsp. *aureofaciens* based on corrosion occurring in spent media devoid of cells. We also excluded MET pathways that are hydrogen-dependent because this strain has no evidence of genes that could oxidize hydrogen gas to generate protons (75–78) (**Supporting Data 1**). Corrosion due to the deprotonation of small organic acids is unlikely because such pathways would have been maintained in spent medium incubated under anoxic conditions where the pH of the medium was the same as oxic incubations. Instead, our proposed mechanism aligns with previous work on fungi, where Cu(0) was abiotically oxidized to Cu(2+) before binding to oxalate, promoting copper corrosion (79).

Our revised mechanistic explanation counters previous studies that linked *phz* genes in *P. aeruginosa* to the corrosion of metal surfaces (26–30, 43) where deletions of *phz* genes (*e.g., phzM* and *phzS*) required to make PYO inhibited corrosion of stainless steel when cells were grown as a biofilm (28). This previous work showed that steel corrosion was restored when PYO was supplied exogenously (28), but it remains unclear whether this was due to phenazines contributing to redox cycling at the metal surface or supporting more robust biofilm growth. Based on our study, more robust growth may have favoured MIC by limiting metal toxicity (*i.e.,* less metal per cell). Siderophore production is linked to biofilm growth (80), offering a potential route for the solubilization of metals such as Fe(2+) that could precipitate if further oxidized. These explanations reflect why it is important to consider the interplay between abiotic and biotic redox cycling reactions, and metabolites that affect metal solubility in mechanistic MIC studies.

The contrast between previous work on iron and our work with cobalt also raises the question: Why did a model phenazine not oxidize Co(0) despite the reaction being thermodynamically favourable under our experimental conditions (*i.e.,* ΔG = −36.74 kJ/mol) (**Table S6**)? Although we calculated Fe(0) oxidation (ΔG = −67.23 kJ/mol) to be slightly more favourable (**Table S6**), the difference in ΔG alone is not enough to explain why such a reaction could proceed for iron and not cobalt. In this case, we suspect kinetic limitations such as competition for Co(0) between various oxidants or slow re-oxidation of phenazines by atmospheric oxygen in the absence of enzymes could limit the capacity for phenazines to oxidize Co(0). Note however, that we effectively ruled out phenazines as oxidants for Co(0) through multiple independent tests. We highlight the notion of kinetic limitations here as another dimension that must be considered when assessing the contributions of abiotic vs biotic corrosion processes, particularly in complex habitats with fluctuating redox gradients.

The alternative mechanism we propose for MIC of cobalt underscores the connections that exist between biosynthetic pathways for redox active metabolites and metabolites that control metal bioavailability. In future experiments, we will combine transcriptomic and metabolomic approaches targeting metallophore biosynthetic genes found in *P. chlororaphis* subsp. *aureofaciens* to confirm the exact genes and metal-ligand complexes responsible for cobalt solubilization. Identifying the genetic and metabolic basis of cobalt corrosion is critical to expanding our understanding of MIC processes that affect the fate of critical metals in different environments and optimizing MIC-based strategies for metal recovery.

There is a pressing need to develop sustainable recovery strategies for elemental metals that occur at mg/kg concentrations in waste streams such as electronic waste (45–47). MIC-based methods that rely on naturally occurring metabolites produced under mesophilic and circumneutral conditions offer an ideal way to circumvent the large environmental footprints associated with current pyrometallurgical and hydrometallurgical extraction methods (81, 82). Despite concerns about MIC damaging infrastructure driving research in the field, our study makes the case for shifting the narrative to how MIC can support sustainable metal reclamation starting with cobalt.

## Methods

### Bacterial Strains and Culture Conditions

All corrosion assays were conducted using bacterium *Pseudomonas chlororaphis* subspecies *aureofaciens* (ATCC 13985), originally isolated from kerosene-contaminated clay of the Maas River. Cells were grown aerobically at 28 °C and shaken at 220 rpm (New Brunswick Scientific Innova® 44) in modified pH 7 Delft mineral salt liquid medium comprised of a basal salt solution, trace metal solution, and glucose solution (54). The basal salt solution, containing 22.27 mM K_2_HPO_4_, 13.59 mM NaH_2_PO_4_, 15.14 mM (NH_4_)_2_SO_4_ and 491.88 µM MgCl_2_·6H_2_O in ultra-filtered (18.2 Ω) MilliQ, was autoclaved at 121 °C for 30 min. This solution was then supplemented with 20.04 mM anhydrous D-glucose as the sole carbon source and trace elements from a concentrated stock solution to yield final concentrations of 34.22 µM EDTA, 6.96 µM ZnSO_4_·7H_2_O, 6.80 µM CaCl_2_·2H_2_O, 17.98 µM FeSO_4_·7H_2_O, 0.83 µM Na_2_MoO_4_·2H_2_O, 0.80 µM CuSO_4_·5H_2_O, 1.68 µM CoCl_2_, and 6.16 µM MnCl_2_·4H_2_O. For the glucose and trace element solutions, 100-fold concentrated stocks were made in ultra-filtered (18.2 Ω) MilliQ water and filter sterilized using a 0.2 µm PES filter prior to addition to the autoclaved basal solution. To make solid defined medium, agar was added at a concentration of 15 g/L to the basal salt solution prior to autoclaving. The method used to generate a dose-response curve for Co(2+) exposure with *P. chlororaphis* subsp. *aureofaciens* is detailed in **Section 3** of the **Supplementary Information**.

### Construction of Δ*phz* and Δ*hcn* deletion mutants

Deletion mutants were created to remove either the *phz* operon (*phzA*, *phzB*, *phzC*, *phzD*, *phzE*, *phzF*, and *phzG*), *hcn* operon (*hcnA*, *hcnB*, and *hcnC*), or both from wild-type *P. chlororaphis* subsp. *aureofaciens* using a previously established 2-step allelic exchange method involving antibiotic resistance and sucrose sensitivity markers encoded by the pK18msB gene replacement plasmid (83, 84). pK18msB was obtained from the Addgene plasmid repository (Addgene plasmid # 177839; http://n2t.net/addgene:177839; RRID: Addgene 177839; deposited by Christopher Johnson, National Renewable Energy Laboratory). Strains, plasmids, and primers used in the strain constructions are listed in **Table 1**, and additional details on protocols are provided in **Section 8** of the **Supporting Information**.

**Table 1:**
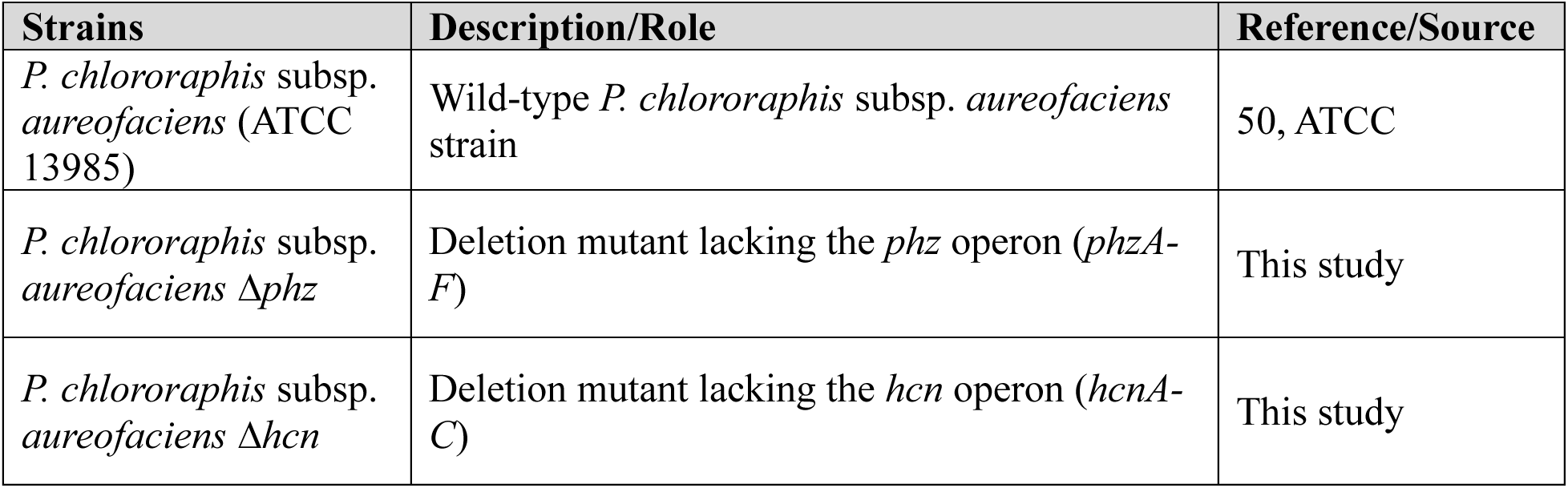
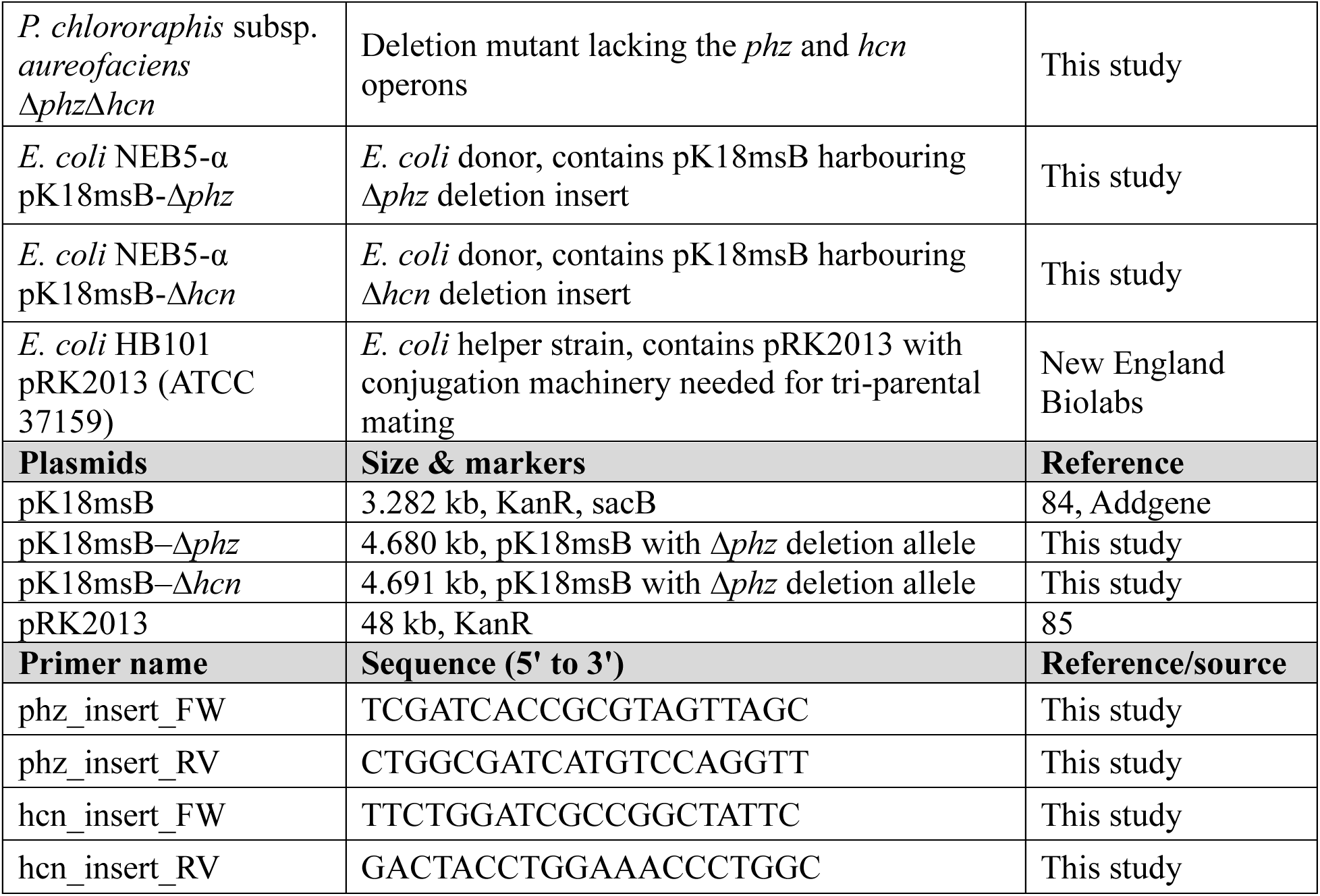
Summary of strains, plasmids, and primers used in genetic manipulations.

The *phz* operon deletion spans from the 31st nucleotide of *phz*A to the 639th nucleotide of *phz*G (*i.e.,* nucleotides 6142474 to 6148700 in wild-type genome). The *hcn* operon deletion spans from the 31st nucleotide of *hcn*A to the 1227th nucleotide of *hcn*C (*i.e.,* nucleotides 2956513 to 2959455 in wild-type genome). To generate each deletion allele, ∼700 bp of homologous sequence upstream and downstream of the deletion junctions were designed as 1.401 kb (*phz*) and 1.414 kb (*hcn*) deletion allele inserts. These inserts were generated as synthetic gene fragments (Twist Bioscience), with SacI and XmaI restriction sites added as extensions to facilitate cloning (**Figures S23 & S24**). SacI and XmaI digested gene fragments were ligated to the digested allelic exchange vector pK18msB using established cloning protocols (83). Ligation reactions were transformed into *E.coli* NEB5-α chemically competent cells (C2988J, New England Biolabs) with selection on Luria-Bertani (LB) agar amended with 30 µg/mL Kanamycin. KanR colonies were screened by colony PCR with respective insert-amplifying primers (**Table 1**) to identify transformants carrying the vector with successfully ligated insert. Candidate clones were verified for KanR by streaking on LB + 30 µg/mL Kan agar and verified for sucrose susceptibility (conferred by *sacB* present on pK18msB) on LB + 30 µg/mL Kan agar + 5% sucrose agar. Plasmid sequences were verified by Nanopore long-read sequencing of plasmid minipreps via Flow Genomics (Ontario, Canada).

Tri-parental mating was conducted to introduce the deletion vectors (pK18msB–Δ*phz* and pK18msB–Δ*hcn*) from *E. coli* NEB5-α donors into the recipient wild-type *P. chlororaphis* subsp. *aureofaciens* with the help of *E. coli* HB101 harbouring pRK2013, which provided necessary conjugation functions. Recipient cultures were incubated at 42 °C for 1 hour prior to setting up the conjugation reactions to inactivate host-nucleases and a 2:2:1 ratio (by volume) of donor:helper:recipient cultures was adopted to increase efficiency. Selection protocols detailed are based on those published by Liu et al (86). Mating mixtures were plated on LB + 50 µg/mL Kan + 100 µg/mL Ampicillin agar to select for homologous recombination of the deletion vector into the recipient genome via the sequences targeting the deletion junction. *P. chlororaphis* subsp. *aureofaciens* exconjugants gain KanR from genomic integration of pK18msB. The ampicillin counter-selects the *E. coli* donor and helper, while sparing the intrinsically AmpR *P. chlororaphis* subsp. *aureofaciens* cells. KanR exconjugants were plated on LB + 10% sucrose agar to select for the second recombination step, in which the pK18msB backbone was excised from the recipient genome via *sacB*-mediated sucrose counterselection.

Sucrose-resistant candidates were screened by colony PCR to distinguish successful deletion mutants possessing the Δ*phz* or Δ*hcn* allele from wild-type revertants (**Figures S25 & S26**). Phenotypic validation of loss of KanR, sucrose-sensitivity, and orange colour indicative of phenazine production was performed by streaking on LB alone, LB + 50 ug/mL Kan, and LB + 10% sucrose agar (**Figure S27**). Following validation of the single deletion mutants, a double deletion mutant was generated by deleting the *phz* operon from the Δ*hcn* strain using the same construction and validation methods (**Figures S27 & S28**). Comparisons of growth in defined medium showed no notable difference across any of the mutants with respect to the wild type (**Figure 3**).

The Δ*phz* and Δ*hcn* deletion mutants were further validated by whole-genome long-read Nanopore sequencing. Genomic DNA was isolated from LB cultures of the wild-type, Δ*phz*, and Δ*hcn* strains (One-4-All Genomic DNA MiniprepKit, BioBasic BS88505) and submitted to Flow Genomics (Ontario, Canada) for sequencing. Library preparation followed the manufacturer’s recommendations using Oxford Nanopore Rapid Chemistry (SQK-RBK114.96) and R10.4.1 flow cells. Sequencing was performed on an Oxford Nanopore GridION, and basecalling used the Sup v5.0.0 model. Genome assemblies provided by Flow Genomics were independently validated using Nanophase v 0.2.3 (87), which uses metaFlye v2.9-b176 (88) for genome assembly prior to genome binning with MetaBAT2 (89) and MaxBin2 (90). Coverage was calculated using minimap2 v.21-r1071 (91) prior to final polished bins being generated with Racon v1.4.22 (92) and medaka 1.4.3 (https://github.com/nanoporetech/medaka). Genomes of the wild type and mutants were aligned and annotated to confirm the absence of respective core operon genes using online software Proksee (93) which uses Prokka (94) for annotation (**Figure S29**). Additional genome annotation methods to assess the presence of biosynthetic gene clusters using the web version of antiSMASH v8.dev-220ad8f9 (95) and DRAM v1.5.0 (96) are detailed in **Section 2** of the **Supporting Information**.

### Co(0) Corrosion Assays

Corrosion assays were conducted by incubating 1 cm segments of pure Co(0) wires (0.5 mm diameter, 99%, 010947.BW, Puratonic) in various abiotic or biotic treatments to observe levels of corrosion over 168 hours. Co(0) wires were weighed and sterilized with 30 sec washes in acetone, ultra-filtered (18.2 Ω) MilliQ water, and ethanol prior to beginning the assays. ‘No wire’ controls were run in tandem with the wire-containing flasks to distinguish between background Co(2+) in the defined medium and Co(2+) resulting from wire corrosion (**Figure S1**). Live cell treatments were prepared by growing wild-type, Δ*phz*, Δ*hcn*, and Δ*phzΔhcn* strains in 5 mL of defined medium. After 16-18 hours of growth, inocula were combined with 45 mL of defined medium in 250 mL flasks to make 10% v/v inocula.

Abiotic treatments containing phenazine standards and spent medium, obtained by vacuum-filtering (0.22 µm pore size PSE filters, 83.3941.001, Sarstedt) 60 mL of wild type and Δ*phz* cultures grown for 24 hours to reach early stationary phase, were conducted alongside experiments with live cells. For abiotic experiments testing for Co(0) corrosion by phenazines alone, a commercially available phenazine-1-carboxylic acid standard (PCA, 98%, CAS: 2538-68-3, CS-W009046, ChemScene LLC), the main phenazine produced by *P. chlororaphis* subsp. *aureofaciens*, was dissolved in HR-GC grade methanol and spiked in sterile medium to a final concentration of 50 µg/mL. 6.5% v/v methanol only controls were also included to control for the addition of solvent. Preliminary experiments were run with DMSO as a solvent to solubilize PCA, however DMSO impeded Co(0) corrosion as observed by ICP-MS data at 168 hours (**Figure S21**) due to crystal formation on the wire (**Figure S22**) such that we omitted the use of DMSO in other experiments. Treatments were run alongside 50 mL of sterile medium to observe baseline corrosion caused by the medium devoid of cells, metabolites, and/or other solvents (**Figure S1**).

Subsampling of assay flasks shaken at 28 °C and 220 rpm occurred at 0, 24, 72, and 168 hours. Flasks were weighed before and after sub-sampling for water and cobalt mass balance purposes. For each subsample, aliquots of 5 mL for LC-HRMS and 2.5-3.5 mL for ICP-MS analyses were transferred to glass vials and polypropylene tubes respectively. 200 µL was taken to measure cell growth by OD_600_ using a BioTek Epoch spectrophotometer. Additionally, viable plate counts were conducted on the same defined medium to assess viability of live cell treatments.

Anoxic conditions for additional corrosion assays were created by bubbling 50 mL of sterile medium and spent medium from filtered Δ*phz* cultures in 150 mL bottles that were crimped shut under a sterile flow of N_2_ gas passed through a 0.22 µm pore size PTFE filter. After purging the bottles of oxygen, sterilized Co(0) wires were added to the bottles, which were decrimped in a Coy anaerobic glovebox containing an atmosphere of 97% N_2_/3 % H_2_. Bottles were subsequently recrimped under the same headspace for incubation. All subsampling was carried out using sterile needles that were scrubbed with sterile N_2_ to remove trace oxygen prior to being filled with N_2_ to ensure volume replacement of the liquid being removed and preclusion of oxygen infiltration during experimental manipulations.

### Co(2+) determination by ICP-MS

Subsampled volumes for ICP-MS were acidified to 2% v/v with trace element grade HNO_3_ (Baker Intra-Analyzed Plus, 9368-01, Avantor) prior to filtering through 0.22 µm pore size hydrophilic PTFE filters. Additional aliquots were taken from select wild type and sterile medium time-series to determine intracellular Co(2+) concentrations. These samples were first passed through 0.22 µm pore size hydrophilic PTFE filters to remove cells and then acidified to 2% v/v HNO_3_, allowing only the extracellular Co(2+) concentration to be detected. All processed samples were then diluted 20-fold prior to ICP-MS analysis. Co(2+) concentrations were determined using two Agilent systems: 7700x ICP-MS with ASX-500 autosampler and 7900x ICP-MS with SPS4 autosampler. Multi-internal standards monitored the recovery of selected elements (7700x ICP-MS = Ge, In; 7900x ICP-MS = Sc, Ge, Y) for quality assurance. Co(2+) standards ranging 0-50 ppm were prepared using VeriSpec® Cobalt Standard for ICP 1000 ppm in 2% HNO_3_ (RV010272-100N, Ricca). To ensure consistency between systems, a comparison was conducted with 4 randomly selected time-series across wild type, spent medium, and sterile medium treatments. With an average of 2.57% analytical variance between instruments, there were no noted analytical issues associated with comparing data from each instrument. Recovered Co(2+) concentrations were used to conduct a mass balance for cobalt in the system, by comparing aqueous Co(2+) corrected for sub-sampling removals to mass losses on the Co(0) wires. Mean Co(2+) recovery across all treatments was 99.13 ± 2.80 % and ranged from 93.17 ± 4.66 % to 125.45 ± 3.57 % (**Table S1**).

### Phenazine quantification by LC-HRMS

5 mL aliquots from duplicate samples of wild-type, Δ*phz*, and spent medium time-series were processed using a modified protocol from Liu et al (86). Briefly, samples underwent three freeze-thaw cycles and were then acidified to pH 2 with 110 µL of 3 M HCl before passing through 0.22 µm pore size hydrophilic PTFE filters. Phenazines were extracted with two volumes of ACS grade ethyl acetate and the combined organic layer was evaporated under N_2_ and reconstituted in 5 mL of HR-GC grade methanol.

Phenazine (PHE, 99+%, CAS: 92-82-0, A15770.06, Thermo Scientific), 2-hydroxyphenazine (2-OHPHZ, 95%, CAS: 4190-95-8, V32899, Ontario Chemicals Inc.), 1-hydroxyphenazine (1-OHPHZ, >95.0%, CAS: 528-71-2, H0289, TCI), phenazine-1-carboxylic acid (PCA, >95.0%, CAS: 2538-68-3, CS-W009046, ChemScene LLC) and phenazine-1-carboxamide (PCN [Oxychlororaphine], 97.46%, CAS: 550-89-0, CS-W022931, ChemScene LLC) standards were sonicated in HR-GC grade methanol until homogenous. These were combined into standards using a 1:1:1:1:1 ratio, where each compound had a final concentration ranging from 0.1 to 7 µg/mL. Extraction efficiency was evaluated by spiking 5 µg/mL of individual phenazine standards into sterile medium. These laboratory fortified solutions underwent extraction and were directly compared to unmanipulated 5-in-1 standards at 5 µg/mL to give mean extraction efficiency values of 81.60 ± 3.85 % for 2-OHPHZ, 84.72 ± 1.58 % for 1-OHPHZ, 84.59 ± 1.69 % for PCA, and 63.99 ± 1.39 % for PCN.

LC-HRMS spectra revealing phenazine species present in cell culture extracts were obtained with an Agilent 6230B Time of Flight (ToF) mass spectrometer coupled to an Agilent 1260 Infinity II HPLC system. An Eclipse Plus C18 RRHD column (2.1 × 100mm, 1.8 μm, Agilent Technologies) maintained at 35 °C was used to separate compounds present in 2 µL injections of 5-in-1 standards and experimental samples. The mobile phases consisted of (A) ddH2O with 0.1% formic acid and (B) acetonitrile (ACN) with 0.1 % formic acid, with a linear gradient ramping from 5 % (B) to 100 % (B) over 11 min at a flow rate of 0.4 mL/min. The Agilent 6230B ToF mass spectrometer was operated in ESI positive mode scanning from 100-1700 m/z. Examples of extracted ion chromatograms for phenazines of interest (**Figure S30**) and a summary of retention times, m/z values, and standard curve linearity (**Table S7 & Table S8**) are provided in **Section 8** of the **Supplementary Information**. Quantification of 2-OHPCA was conducted using PCA standards as a proxy.

### Co(0) wire surface analysis using SEM-EDX

After the 168-hour corrosion assays, Co(0) wires were gently rinsed with deionized water and dried under N_2_ before being weighed. For the anoxic wires, rinsing occurred under anoxic conditions with deoxygenated water in an anaerobic chamber where they were stored in 1.5 mL plastic tubes until weighing to minimize exposure to oxygen. Wires were imaged and measured by scanning-electron microscope (SEM, TESCAN VegaII XMU) for evidence of corrosion at 20 kV using backscattered electron (BSE) and low-vacuum secondary electron (LVSTD) detectors. Elemental surface composition was analyzed using an energy-dispersive X-ray spectrometer (EDX, Oxford Inca Energy X-Act). Selection of sections for scanning was driven by shading in images, wherein lighter areas indicated of regions high in pure cobalt compared to darker regions where potential oxides and passivation layers formed. All raw images and EDX data are provided through a repository: https://zenodo.org/records/17408329. Rough estimates of Co:O ratios were calculated by dividing the percent weight of cobalt over that of oxygen from selected EDX scans, as presented in **Table S5** and rounded to whole integers in the **Results & Discussion** section.

### Statistical analyses

All statistical analyses and data visualizations for ICP-MS, LC-HRMS, and plate counts were carried out using R v 4.3.2 in R studio 2024.09.0+375. Data manipulations and visualizations were generated using the tidyverse v 2.0.0 and patchwork v 1.3.0 packages. The main statistical tests conducted to compare Co(2+) concentrations were ANOVAs run with type III sum of squares to account for the unbalanced design of some experimental treatments. Where needed, data was log10 transformed to ensure assumptions for linear models required for parametric ANOVA tests were met. Multiple comparisons were carried out using the Tukey post-hoc test in R using 95% confidence, wherein a p-value < 0.05 was deemed statistically significant. Additional details on other statistical analyses and results are provided where relevant in figure legends within the **Supplementary Information.**

## Data availability statement

All data underlying main and supporting figures and the associated R notebooks containing code required to reproduce statistical analyses and visualizations has been uploaded to the GitHub page: https://github.com/carleton-envbiotech/Cobalt_corrosion/tree/main. All bash code required for genome annotation workflows has also been provided on the same GitHub page. Finally, all SEM images and summaries of EDX analysis from this work have been uploaded to Zenodo, and can be accessed using the following link: https://zenodo.org/records/17408329. All sequencing data for the wild type and deletion mutants has also been uploaded to NCBI under Bioproject PRJNA1336231. Samples were uploaded under biosample numbers SAMN52040257, SAMN52040258, and SAMN52040259, respectively. These samples were also uploaded to NCBI’s SRA database under SRR35667903, SRR35667902, and SRR35667901. Due to the suspension of the NCBI submission portal at the time of submission, we have also provided the genomes in **Supporting Data 2, Supporting Data 3,** and **Supporting Data 4**.

## Acknowledgements

AD contributed to the conceptualization, methodology, investigation, formal analysis, validation, writing the original draft. AK contributed to the initial conceptualization, methodology, investigation, formal analysis, and validation. AH contributed to the method development and provided resources used for the genetic manipulations. DRM contributed to the LC-HRMS methodology and instrumentation. DSG contributed to the conceptualization, data curation, formal analysis, project administration, resources, software, supervision, reviewing, and editing. All authors contributed to reviewing and editing the manuscript for submission. This work was supported by an NSERC CGS-M scholarship to AD and an NSERC Discovery Grant awarded to DSG (#RGPIN-2022-04891 and DGECR-2022-00512).

## Notes

https://github.com/carleton-envbiotech/Cobalt_corrosion/tree/main.

https://zenodo.org/records/17408329

### Competing Interest Statement

The authors have declared no competing interest.

### Summary of Updates

New data has been included using double deletion mutants and anaerobic experiments; A new mechanistic explanation has been summarized in Fgure 5; Major streamlining to the narrative was carried out to account for the new data and mechanistic insights

## References

1. Xu D, Gu T, Lovley DR. 2023. Microbially mediated metal corrosion. Nat Rev Microbiol 21:705–718.

2. Cámara M, Green W, MacPhee CE, Rakowska PD, Raval R, Richardson MC, Slater-Jefferies J, Steventon K, Webb JS. 2022. Economic significance of biofilms: a multidisciplinary and cross-sectoral challenge. npj Biofilms and Microbiomes 8:1–8.

3. Amendola R, Acharjee A. 2022. Microbiologically Influenced Corrosion of Copper and Its Alloys in Anaerobic Aqueous Environments: A Review. Frontiers in Microbiology 13.

4. Wang H, Ju L-K, Castaneda H, Cheng G, Zhang Newby B. 2014. Corrosion of carbon steel C1010 in the presence of iron oxidizing bacteria *Acidithiobacillus ferrooxidans*. Corrosion Science 89:250–257.

5. Booth GH. 1964. Sulphur Bacteria in Relation to Corrosion. Journal of Applied Bacteriology 27:174–181.

6. Gu T, Wang D, Lekbach Y, Xu D. 2021. Extracellular electron transfer in microbial biocorrosion. Current Opinion in Electrochemistry 29:100763.

7. Kracke F, Vassilev I, Krömer JO. 2015. Microbial electron transport and energy conservation – the foundation for optimizing bioelectrochemical systems. Front Microbiol 6:575.

8. Tang H-Y, Holmes DE, Ueki T, Palacios PA, Lovley DR. 2019. Iron Corrosion via Direct Metal-Microbe Electron Transfer. mBio 10:e00303–19.

9. Tang H-Y, Yang C, Ueki T, Pittman CC, Xu D, Woodard TL, Holmes DE, Gu T, Wang F, Lovley DR. 2021. Stainless steel corrosion via direct iron-to-microbe electron transfer by Geobacter species. The ISME Journal 15:3084–3093.

10. Hernández-Santana A, Suflita JM, Nanny MA. 2022. *Shewanella oneidensis* MR-1 accelerates the corrosion of carbon steel using multiple electron transfer mechanisms. International Biodeterioration & Biodegradation 173:105439.

11. Holmes DE, Dang Y, Walker DJF, Lovley DR. 2016. The electrically conductive pili of Geobacter species are a recently evolved feature for extracellular electron transfer. Microbial Genomics 2:e000072.

12. Liang D, Liu X, Woodard TL, Holmes DE, Smith JA, Nevin KP, Feng Y, Lovley DR. 2021. Extracellular Electron Exchange Capabilities of *Desulfovibrio ferrophilus* and Desulfopila corrodens. Environ Sci Technol 55:16195–16203.

13. Zhou E, Li F, Zhang D, Xu D, Li Z, Jia R, Jin Y, Song H, Li H, Wang Q, Wang J, Li X, Gu T, Homborg AM, Mol JMC, Smith JA, Wang F, Lovley DR. 2022. Direct microbial electron uptake as a mechanism for stainless steel corrosion in aerobic environments. Water Research 219:118553.

14. Holmes DE, Tang H, Woodard T, Liang D, Zhou J, Liu X, Lovley DR. 2022. Cytochrome-mediated direct electron uptake from metallic iron by *Methanosarcina acetivorans*. mLife 1:443–447.

15. Woodard TL, Ueki T, Lovley DR. H_2_ Is a Major Intermediate in Desulfovibrio vulgaris Corrosion of Iron. mBio 14:e00076–23.

16. Tsurumaru H, Ito N, Mori K, Wakai S, Uchiyama T, Iino T, Hosoyama A, Ataku H, Nishijima K, Mise M, Shimizu A, Harada T, Horikawa H, Ichikawa N, Sekigawa T, Jinno K, Tanikawa S, Yamazaki J, Sasaki K, Yamazaki S, Fujita N, Harayama S. 2018. An extracellular [NiFe] hydrogenase mediating iron corrosion is encoded in a genetically unstable genomic island in *Methanococcus maripaludis*. Sci Rep 8:15149.

17. Lahme S, Mand J, Longwell J, Smith R, Enning D. 2021. Severe Corrosion of Carbon Steel in Oil Field Produced Water Can Be Linked to Methanogenic Archaea Containing a Special Type of [NiFe] Hydrogenase. Applied and Environmental Microbiology 87:e01819–20.

18. Uchiyama T, Ito K, Mori K, Tsurumaru H, Harayama S. 2010. Iron-Corroding Methanogen Isolated from a Crude-Oil Storage Tank. Applied and Environmental Microbiology 76:1783–1788.

19. Jia R, Tan JL, Jin P, Blackwood DJ, Xu D, Gu T. 2018. Effects of biogenic H_2_S on the microbiologically influenced corrosion of C1018 carbon steel by sulfate reducing *Desulfovibrio vulgaris* biofilm. Corrosion Science 130:1–11.

20. Kato S, Yumotot I, Kamagata Y. 2015. Isolation of Acetogenic Bacteria That Induce Biocorrosion by Utilizing Metallic Iron as the Sole Electron Donor. Applied and Environmental Microbiology 81:67.

21. Davidova IA, Duncan KE, Wiley G, Najar FZ. 2022. Desulfoferrobacter suflitae gen. nov., sp. nov., a novel sulphate-reducing bacterium in the Deltaproteobacteria capable of autotrophic growth with hydrogen or elemental iron. International Journal of Systematic and Evolutionary Microbiology 72:005483.

22. Jin Y, Li J, Zhang M, Zheng B, Xu D, Gu T, Wang F. 2024. Effect of exogenous flavins on the microbial corrosion by *Geobacter sulfurreducens* via iron-to-microbe electron transfer. Journal of Materials Science & Technology 171:129–138.

23. Light SH, Su L, Rivera-Lugo R, Cornejo JA, Louie A, Iavarone AT, Ajo-Franklin CM, Portnoy DA. 2018. A flavin-based extracellular electron transfer mechanism in diverse Gram-positive bacteria. Nature 562:140–144.

24. Wang D, Kijkla P, Mohamed ME, Saleh MA, Kumseranee S, Punpruk S, Gu T. 2021. Aggressive corrosion of carbon steel by *Desulfovibrio ferrophilus* IS5 biofilm was further accelerated by riboflavin. Bioelectrochemistry 142:107920.

25. Yi Y, Zhao T, Zang Y, Xie B, Liu H. 2021. Different mechanisms for riboflavin to improve the outward and inward extracellular electron transfer of *Shewanella loihica*. Electrochemistry Communications 124:106966.

26. Huang L, Chang W, Zhang D, Huang Y, Li Z, Lou Y, Qian H, Jiang C, Li X, Mol A. 2022. Acceleration of corrosion of 304 stainless steel by outward extracellular electron transfer of *Pseudomonas aeruginosa* biofilm. Corrosion Science 199:110159.

27. Yang Y, Zhou E, Li L, Peng X, Huang Y, Jiang C, Gu T, Wang F, Xu D. 2025. The role of phenazines in marine *Pseudomonas aeruginosa* microbiologically influenced corrosion against 316L stainless steel. Corrosion Science 242:112587.

28. Huang L, Huang Y, Lou Y, Qian H, Xu D, Ma L, Jiang C, Zhang D. 2020. Pyocyanin-modifying genes *phz*M and *phz*S regulated the extracellular electron transfer in microbiologically-influenced corrosion of X80 carbon steel by *Pseudomonas aeruginosa*. Corrosion Science 164:108355.

29. Zhou E, Zhang M, Huang Y, Li H, Wang J, Jiang G, Jiang C, Xu D, Wang Q, Wang F. 2022. Accelerated biocorrosion of stainless steel in marine water via extracellular electron transfer encoding gene *phz*H of *Pseudomonas aeruginosa*. Water Research 220:118634.

30. Huang Y, Zhou E, Jiang C, Jia R, Liu S, Xu D, Gu T, Wang F. 2018. Endogenous phenazine-1-carboxamide encoding gene *phz*H regulated the extracellular electron transfer in biocorrosion of stainless steel by marine *Pseudomonas aeruginosa*. Electrochemistry Communications 94:9–13.

31. Price-Whelan A, Dietrich LEP, Newman DK. 2006. Rethinking “secondary” metabolism: physiological roles for phenazine antibiotics. Nat Chem Biol 2:71–78.

32. Price-Whelan A, Dietrich LEP, Newman DK. 2007. Pyocyanin Alters Redox Homeostasis and Carbon Flux through Central Metabolic Pathways in *Pseudomonas aeruginosa* PA14. Journal of Bacteriology 189:6372–6381.

33. Wang J, Xiong F, Liu H, Zhang T, Li Y, Li C, Xia W, Wang H, Liu H. 2019. Study of the corrosion behavior of *Aspergillus niger* on 7075-T6 aluminum alloy in a high salinity environment. Bioelectrochemistry 129:10–17.

34. Jirón-Lazos U, Corvo F, De la Rosa SC, García-Ochoa EM, Bastidas DM, Bastidas JM. 2018. Localized corrosion of aluminum alloy 6061 in the presence of *Aspergillus niger*. International Biodeterioration & Biodegradation 133:17–25.

35. Dai X, Wang H, Ju L-K, Cheng G, Cong H, Newby BZ. 2016. Corrosion of aluminum alloy 2024 caused by *Aspergillus niger*. International Biodeterioration & Biodegradation 115:1–10.

36. Onan M, Ilhan-Sungur E, Güngör ND, Cansever N. 2018. Biocides Effect on the Microbiologically Influenced Corrosion of Pure Copper by *Desulfovibrio* sp. J Electrochem Sci Technol 9:44–50.

37. Pu Y, Dou W, Gu T, Tang S, Han X, Chen S. 2020. Microbiologically influenced corrosion of Cu by nitrate reducing marine bacterium *Pseudomonas aeruginosa*. Journal of Materials Science & Technology 47:10–19.

38. Chen Z, Dou W, Chen S, Pu Y, Xu Z. 2022. Influence of nutrition on Cu corrosion by *Desulfovibrio vulgaris* in anaerobic environment. Bioelectrochemistry 144:108040.

39. Pu Y, Tian Y, Hou S, Dou W, Chen S. 2023. Carbon starvation considerably accelerated nickel corrosion by *Desulfovibrio vulgaris*. Bioelectrochemistry 153:108453.

40. Pu Y, Tian Y, Hou S, Dou W, Chen S. 2023. Enhancement of exogenous riboflavin on microbiologically influenced corrosion of nickel by electroactive Desulfovibrio vulgaris biofilm. npj Mater Degrad 7:1–16.

41. San NO, Nazır H, Dönmez G. 2014. Microbially influenced corrosion and inhibition of nickel–zinc and nickel–copper coatings by *Pseudomonas aeruginosa*. Corrosion Science 79:177–183.

42. Michalska J, Sowa M, Socha RP, Simka W, Cwalina B. 2017. The influence of *Desulfovibrio desulfuricans* bacteria on a Ni-Ti alloy: Electrochemical behavior and surface analysis. Electrochimica Acta 249:135–144.

43. Liu D, Yang H, Li J, Li J, Dong Y, Yang C, Jin Y, Yassir L, Li Z, Hernandez D, Xu D, Wang F, Smith JA. 2021. Electron transfer mediator PCN secreted by aerobic marine *Pseudomonas aeruginosa* accelerates microbiologically influenced corrosion of TC4 titanium alloy. Journal of Materials Science & Technology 79:101–108.

44. Saleem Khan M, Li Z, Yang K, Xu D, Yang C, Liu D, Lekbach Y, Zhou E, Kalnaowakul P. 2019. Microbiologically influenced corrosion of titanium caused by aerobic marine bacterium *Pseudomonas aeruginosa*. Journal of Materials Science & Technology 35:216–222.

45. Purchase D, Abbasi G, Bisschop L, Chatterjee D, Ekberg C, Ermolin M, Fedotov P, Garelick H, Isimekhai K, Kandile NG, Lundström M, Matharu A, Miller BW, Pineda A, Popoola OE, Retegan T, Ruedel H, Serpe A, Sheva Y, Surati KR, Walsh F, Wilson BP, Wong MH. 2020. Global occurrence, chemical properties, and ecological impacts of e-wastes (IUPAC Technical Report). Pure and Applied Chemistry 92:1733–1767.

46. Jain M, Kumar D, Chaudhary J, Kumar S, Sharma S, Singh Verma A. 2023. Review on E-waste management and its impact on the environment and society. Waste Management Bulletin 1:34–44.

47. Han W, Gao G, Geng J, Li Y, Wang Y. 2018. Ecological and health risks assessment and spatial distribution of residual heavy metals in the soil of an e-waste circular economy park in Tianjin, China. Chemosphere 197:325–335.

48. Bilal M, Guo S, Iqbal HMN, Hu H, Wang W, Zhang X. 2017. Engineering *Pseudomonas* for phenazine biosynthesis, regulation, and biotechnological applications: a review. World J Microbiol Biotechnol 33:1–11.

49. Maddula VSRK, Pierson EA, Pierson LS. 2008. Altering the Ratio of Phenazines in *Pseudomonas chlororaphis (aureofaciens)* Strain 30-84: Effects on Biofilm Formation and Pathogen Inhibition. Journal of Bacteriology 190:2759–2766.

50. Kluyver AJ. 1956. *Pseudomonas aureofaciens* nov. spec. and its pigments. J Bacteriol 72:406–411.

51. Harris DC. 2010. Quantitative Chemical Analysis, 8th ed. W. H. Freeman and Company.

52. Wang Y, Newman DK. 2008. Redox Reactions of Phenazine Antibiotics with Ferric (Hydr)oxides and Molecular Oxygen. Environ Sci Technol 42:2380–2386.

53. Faramarzi MA, Mogharabi-Manzari M, Brandl H. 2020. Bioleaching of metals from wastes and low-grade sources by HCN-forming microorganisms. Hydrometallurgy 191:105228.

54. Franco A, Elbahnasy M, Rosenbaum MA. 2023. Screening of natural phenazine producers for electroactivity in bioelectrochemical systems. Microbial Biotechnology 16:579–594.

55. Biessy A, Filion M. 2018. Phenazines in plant-beneficial *Pseudomonas* spp.: biosynthesis, regulation, function and genomics. Environmental Microbiology 20:3905–3917.

56. Shin D, Park J, Jeong J, Kim B. 2015. A biological cyanide production and accumulation system and the recovery of platinum-group metals from spent automotive catalysts by biogenic cyanide. Hydrometallurgy 158:10–18.

57. Pourhossein F, Mousavi SM, Beolchini F, Lo Martire M. 2021. Novel green hybrid acidic-cyanide bioleaching applied for high recovery of precious and critical metals from spent light emitting diode lamps. Journal of Cleaner Production 298:126714.

58. Wargo MJ, Szwergold BS, Hogan DA. 2008. Identification of Two Gene Clusters and a Transcriptional Regulator Required for *Pseudomonas aeruginosa* Glycine Betaine Catabolism. Journal of Bacteriology 190:2690–2699.

59. Anand A, Falquet L, Abou-Mansour E, L’Haridon F, Keel C, Weisskopf L. 2023. Biological hydrogen cyanide emission globally impacts the physiology of both HCN-emitting and HCN-perceiving *Pseudomonas*. mBio 14:e00857–23.

60. Wang S, Pearson LA, Mazmouz R, Liu T, Neilan BA. Heterologous Expression and Biochemical Analysis Reveal a Schizokinen-Based Siderophore Pathway in *Leptolyngbya* (Cyanobacteria). Appl Environ Microbiol 88:e02373–21.

61. Chuljerm H, Deeudom M, Fucharoen S, Mazzacuva F, Hider RC, Srichairatanakool S, Cilibrizzi A. 2020. Characterization of two siderophores produced by *Bacillus megaterium*: A preliminary investigation into their potential as therapeutic agents. Biochimica et Biophysica Acta (BBA) - General Subjects 1864:129670.

62. Wang Q, Liu Q, Ma Y, Zhou L, Zhang Y. 2007. Isolation, sequencing and characterization of cluster genes involved in the biosynthesis and utilization of the siderophore of marine fish pathogen *Vibrio alginolyticus*. Arch Microbiol 188:433–439.

63. Reydams H, Toledo-Silva B, Mertens K, Piepers S, Vereecke N, Souza FN, Haesebrouck F, De Vliegher S. 2024. Phenotypic and genotypic assessment of iron acquisition in diverse bovine-associated non-aureus staphylococcal strains. Vet Res 55:6.

64. Demange P, Bateman A, Dell A, Abdallah MA. 1988. Structure of azotobactin D, a siderophore of *Azotobacter vinelandii* strain D (CCM 289). Biochemistry 27:2745–2752.

65. Ghssein G, Ezzeddine Z. 2022. A Review of *Pseudomonas aeruginosa* Metallophores: Pyoverdine, Pyochelin and Pseudopaline. Biology (Basel) 11:1711.

66. Braud A, Geoffroy V, Hoegy F, Mislin GLA, Schalk IJ. 2010. Presence of the siderophores pyoverdine and pyochelin in the extracellular medium reduces toxic metal accumulation in *Pseudomonas aeruginosa* and increases bacterial metal tolerance. Environmental Microbiology Reports 2:419–425.

67. Siunova TV, Kochetkov VV, Validov SZ, Suzina NE, Boronin AM. 2002. The Production of Phenazine Antibiotics by the *Pseudomonas aureofaciens* Strain with Plasmid-Controlled Resistance to Cobalt and Nickel. Microbiology 71:670–676.

68. Barras F, Fontecave M. 2011. Cobalt stress in Escherichia coli and Salmonella enterica: molecular bases for toxicity and resistance†. Metallomics 3:1130–1134.

69. Dulay H, Tabares M, Kashefi K, Reguera G. 2020. Cobalt Resistance via Detoxification and Mineralization in the Iron-Reducing Bacterium *Geobacter sulfurreducens*. Front Microbiol 11.

70. Chen B-Y, Wu C-H, Chang J-S. 2006. An assessment of the toxicity of metals to *Pseudomonas aeruginosa* PU21 (Rip64). Bioresource Technology 97:1880–1886.

71. Greenwood NN, Earnshaw A. 2012. Chemistry of the Elements, 2nd ed. Elsevier Butterworth-Heinemann.

72. Yang J, Liu H, Martens WN, Frost RL. 2010. Synthesis and Characterization of Cobalt Hydroxide, Cobalt Oxyhydroxide, and Cobalt Oxide Nanodiscs. J Phys Chem C 114:111–119.

73. Crespo A, Blanco-Cabra N, Torrents E. 2018. Aerobic Vitamin B12 Biosynthesis Is Essential for *Pseudomonas aeruginosa* Class II Ribonucleotide Reductase Activity During Planktonic and Biofilm Growth. Frontiers in Microbiology 9.

74. Debussche L, Couder M, Thibaut D, Cameron B, Crouzet J, Blanche F. 1992. Assay, purification, and characterization of cobaltochelatase, a unique complex enzyme catalyzing cobalt insertion in hydrogenobyrinic acid a,c-diamide during coenzyme B12 biosynthesis in *Pseudomonas denitrificans*. Journal of Bacteriology 174:7445–7451.

75. Bürstel I, Siebert E, Frielingsdorf S, Zebger I, Friedrich B, Lenz O. 2016. CO synthesized from the central one-carbon pool as source for the iron carbonyl in O_2_-tolerant [NiFe]-hydrogenase. Proceedings of the National Academy of Sciences 113:14722–14726.

76. De Rosa E, Checchetto V, Franchin C, Bergantino E, Berto P, Szabò I, Giacometti GM, Arrigoni G, Costantini P. 2015. [NiFe]-hydrogenase is essential for cyanobacterium Synechocystis sp. PCC 6803 aerobic growth in the dark. Sci Rep 5:12424.

77. Vignais PM, Billoud B, Meyer J. 2001. Classification and phylogeny of hydrogenases. FEMS Microbiology Reviews 25:455–501.

78. Coppi MV, O’Neil RA, Lovley DR. 2004. Identification of an Uptake Hydrogenase Required for Hydrogen-Dependent Reduction of Fe(III) and Other Electron Acceptors by *Geobacter sulfurreducens*. Journal of Bacteriology 186:3022–3028.

79. Zhao J, Csetenyi L, Gadd GM. 2020. Biocorrosion of copper metal by *Aspergillus niger*. International Biodeterioration & Biodegradation 154:105081.

80. Harrison F, Buckling A. 2009. Siderophore production and biofilm formation as linked social traits. ISME J 3:632–634.

81. Pollmann K, Kutschke S, Matys S, Raff J, Hlawacek G, Lederer FL. 2018. Bio-recycling of metals: Recycling of technical products using biological applications. Biotechnology Advances 36:1048–1062.

82. Jerez CA. 2017. Biomining of metals: how to access and exploit natural resource sustainably. Microbial Biotechnology 10:1191–1194.

83. Hmelo LR, Borlee BR, Almblad H, Love ME, Randall TE, Tseng BS, Lin C, Irie Y, Storek KM, Yang JJ, Siehnel RJ, Howell PL, Singh PK, Tolker-Nielsen T, Parsek MR, Schweizer HP, Harrison JJ. 2015. Precision-engineering the *Pseudomonas aeruginosa* genome with two-step allelic exchange. Nature Protocols 10:1820.

84. Ling C, Peabody GL, Salvachúa D, Kim Y-M, Kneucker CM, Calvey CH, Monninger MA, Munoz NM, Poirier BC, Ramirez KJ, St. John PC, Woodworth SP, Magnuson JK, Burnum-Johnson KE, Guss AM, Johnson CW, Beckham GT. 2022. Muconic acid production from glucose and xylose in *Pseudomonas putida* via evolution and metabolic engineering. Nature Communications 13.

85. Figurski DH, Helinski DR. 1979. Replication of an origin-containing derivative of plasmid RK2 dependent on a plasmid function provided in trans. Proc Natl Acad Sci U S A 76:1648–1652.

86. Liu K, Hu H, Wang W, Zhang X. 2016. Genetic engineering of *Pseudomonas chlororaphis* GP72 for the enhanced production of 2-Hydroxyphenazine. Microbial Cell Factories 15:131.

87. Liu L, Yang Y, Deng Y, Zhang T. 2022. Nanopore long-read-only metagenomics enables complete and high-quality genome reconstruction from mock and complex metagenomes. Microbiome 10:209.

88. Kolmogorov M, Bickhart DM, Behsaz B, Gurevich A, Rayko M, Shin SB, Kuhn K, Yuan J, Polevikov E, Smith TPL, Pevzner PA. 2020. metaFlye: scalable long-read metagenome assembly using repeat graphs. Nature Methods 17:1103–1110.

89. Kang DD, Li F, Kirton E, Thomas A, Egan R, An H, Wang Z. 2019. MetaBAT 2: an adaptive binning algorithm for robust and efficient genome reconstruction from metagenome assemblies. PeerJ 7:e7359.

90. Wu Y-W, Simmons BA, Singer SW. 2016. MaxBin 2.0: an automated binning algorithm to recover genomes from multiple metagenomic datasets. Bioinformatics 32:605–607.

91. Li H. 2018. Minimap2: pairwise alignment for nucleotide sequences. Bioinformatics 34:3094–3100.

92. Vaser R, Sović I, Nagarajan N, Šikić M. 2017. Fast and accurate de novo genome assembly from long uncorrected reads. Genome Res 27:737–746.

93. Grant JR, Enns E, Marinier E, Mandal A, Herman EK, Chen C-Y, Graham M, Van Domselaar G, Stothard P. 2023. Proksee: in-depth characterization and visualization of bacterial genomes. Nucleic Acids Res 51:W484–W492.

94. Seemann T. 2014. Prokka: rapid prokaryotic genome annotation. Bioinformatics 30:2068–2069.

95. Blin K, Shaw S, Vader L, Szenei J, Reitz ZL, Augustijn HE, Cediel-Becerra JDD, de Crécy-Lagard V, Koetsier RA, Williams SE, Cruz-Morales P, Wongwas S, Segurado Luchsinger AE, Biermann F, Korenskaia A, Zdouc MM, Meijer D, Terlouw BR, van der Hooft JJJ, Ziemert N, Helfrich EJN, Masschelein J, Corre C, Chevrette MG, van Wezel GP, Medema MH, Weber T. 2025. antiSMASH 8.0: extended gene cluster detection capabilities and analyses of chemistry, enzymology, and regulation. Nucleic Acids Research gkaf334.

96. Shaffer M, Borton MA, McGivern BB, Zayed AA, La Rosa SL, Solden LM, Liu P, Narrowe AB, Rodríguez-Ramos J, Bolduc B, Gazitúa MC, Daly RA, Smith GJ, Vik DR, Pope PB, Sullivan MB, Roux S, Wrighton KC. 2020. DRAM for distilling microbial metabolism to automate the curation of microbiome function. Nucleic Acids Res 48:8883–8900.

