## Supplementary Information for "Metal-Binding Ligands Rather Than Redox Active Metabolites Are Essential to Microbially-Induced Corrosion of Cobalt"


<u>1) Additional ICP-MS data, mass balances, and statistical tests conducted</u> (p.2)

- **Table S1:** Average wire mass losses and average mass balance recoveries.
- **Figure S1:** Contribution of background Co(2+) in medium from "no wire" controls.
- **Figure S2:** Boxplot of log10 [Co(2+)] at 168 hours across abiotic and biotic treatments.
- **Figure S3:** Boxplot of [Co(2+)] at 168 hours for abiotic metabolite treatments.

<u>2) Biosynthetic gene cluster analyses</u> (p.5)

- **Bioinformatic analyses**
- **Figure S4:** Summary of antiSMASH output for biosynthetic gene clusters predicted to produce metabolites relevant to cobalt solubilization and transport.
- **Table S2:** DRAM annotations for core biosynthetic genes and select transport genes.
- **Table S3:** Predicted compounds made by gene clusters found by antiSMASH.

<u>3) Cell-associated Co(2+) and toxicity measurements</u> (p.9)

- **Figure S5:** Comparison of [Co(2+)] between acidify-filter and filter-acidify preservation methods with wild-type live cells.
- **Protocol for generating dose-response curve for Co(2+) exposure**
- **Figure S6:** Cobalt dose-response curve reporting growth rate of *P. chlororaphis* subsp. *aureofaciens* in medium amended with $CoCl_2$.
- **Figure S7:** Comparison of [Co(2+)] between acidify-filter and filter-acidify preservation methods with sterile medium.

<u>4) Additional SEM images, wire diameters, and EDX data</u> (p.12)

- **Table S4:** Diameters and pit depths of representative wires.
- **Figure S8-S18:** EDX scans of representative wires for each treatment.
- **Table S5:** Percent surface composition for EDX spectra of representative wires.

<u>5) Statistical analyses of ICP-MS data and SEM of anoxic and DMSO treatments</u> (p.19)

- **Figure S19:** Boxplot of [Co(2+)] at 168 hours comparing oxic and anoxic treatments
- **Figure S20:** Comparison of SEM images of wires across oxic and anoxic conditions.
- **Figure S21:** Boxplot of [Co(2+)] at 168 hours across DMSO biotic treatments.
- **Figure S22:** SEM image of Co(0) wire from sterile DMSO treatment after 168 hours.

<u>6) Redox reaction comparison between Fe & Co</u> (p.22)

- **Table S6:** Redox reaction and thermodynamic calculations using PCA concentration from sterile medium experiments.

<u>7) Genetic manipulations and validation</u> (p.23)

- **Detailed protocols and background used in deletion mutant generation**
- **Figure S23:** Annotated FASTA sequence of Δ*phz* deletion allele.
- **Figure S24:** Annotated FASTA sequence of Δ*hcn* deletion allele.
- **Figures S25-S27:** Colony PCR results identifying Δ*phz*, Δ*hcn*, and Δ*phz*Δ*hcn* mutants.
- **Figure S28:** Phenotypic validation for all deletion mutant candidates.
- **Figure S29:** Sequencing alignments of Δ*phz* and Δ*hcn* mutants to WT genome.

<u>8) LC-HRMS QA/QC</u> (p.31)

- **Figure S30:** Example EICs for phenazines of interest from wild-type sample and standards.
- **Table S7:** Retention times and m/z ratios of phenazines of interest.
- **Table S8:** Calibration curve equations for PCA and 2-OHPHZ on LC-HRMS.

<u>10) Supporting References</u> (p.33)

1) Additional ICP-MS data, mass balances, and statistical tests conducted

**Table S1:** Mean wire mass losses determined by weight, as well as mean cobalt and water mass balance recoveries for all abiotic and biotic corrosion assay treatments. Note that F-A indicates treatments where samples were filtered then acidified for ICP-MS preservation, while all other treatments were acidified then filtered. Weighted averages for the cobalt and water percent recoveries were calculated taking into account unequal sample sizes for each treatment.

| **Treatment** | **Mean wire mass relative loss (%)** | **Mean cobalt recovery (%)** | **Mean water recovery (%)** |
|---|---|---|---|
| Sterile medium | 3.65 ± 1.47 | 99.01 ± 1.28 | 96.35 ± 3.47 |
| Sterile medium (F-A) | 3.65 ± 1.47 | 97.86 ± 2.28 | 97.54 ± 1.22 |
| Sterile medium anoxic | 1.94 ± 0.66 | 98.87± 0.50 | 99.94 ± 0.22 |
| DMSO | 1.79 ± 2.26 | 98.85 ± 1.92 | 97.04 ± 2.38 |
| DMSO (F-A) | 1.79 ± 2.26 | 97.80 ± 2.47 | 97.04 ± 2.38 |
| MeOH | 2.31 ± 0.83 | 99.92 ± 0.74 | 91.62 ± 3.11 |
| PCA | 2.40 ± 0.56 | 99.51 ± 0.49 | 95.09 ± 0.19 |
| Spent medium | 10.02 ± 2.83 | 95.99 ± 2.59 | 96.43 ± 2.10 |
| Spent medium Δ*phz* | 14.86 ± 1.97 | 99.21 ± 0.50 | 97.67 ± 1.28 |
| Spent medium Δ*phz* anoxic | 0.69 ± 1.33 | 100.13 ± 1.60 | 99.93 ± 0.12 |
| Wild type | 26.22 ± 12.02 | 94.05 ± 6.34 | 96.86 ± 1.52 |
| Wild type (F-A) | 26.22 ± 12.02 | 93.17 ± 4.66 | 97.01 ± 1.92 |
| Wildtype with DMSO | 5.69 ± 2.31 | 98.95 ± 1.09 | 97.89 ± 0.72 |
| Wildtype with DMSO (F-A) | 5.69 ± 2.31 | 97.55 ± 1.38 | 97.89 ± 0.72 |
| Δ*phz* | 34.11 ± 2.65 | 125.45 ± 3.57 | 92.44 ± 3.02 |
| Δ*hcn* | 20.10 ± 2.47 | 96.25 ± 0.47 | 97.51 ± 0.48 |
| Δ*phz*Δ*hcn* | 51.44 ± 5.55 | 95.79 ± 2.11 | 95.82 ± 1.96 |
| **Weighted average across all treatments** | | 99.13 ± 2.80 | 96.67 ± 2.16 |

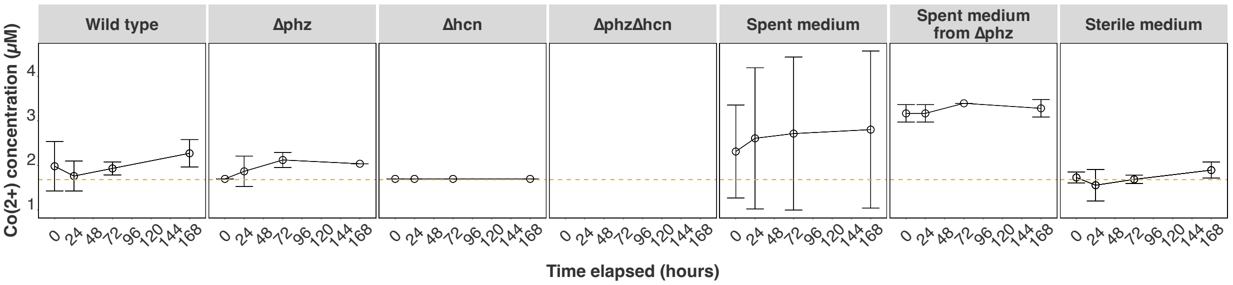


**Figure S1:** Mean Co(2+) concentrations ± SD obtained by ICP-MS for "no wire" controls demonstrating the background contribution of Co(2+) from the defined medium across treatments over 168 hours. The orange dashed line indicates the theoretical amount of Co(2+) expected in the defined medium (*i.e.,* 1.68μM). Mean concentrations were calculated for the following number of replicates: Wild type: n = 6, Δ*phz*: n = 4, Δ*hcn*: n = 4, Δ*phz*Δ*hcn*: n = 4, Spent medium: n = 4, Spent medium from Δ*phz*: n = 3, and Sterile medium: n = 9. A blank panel is present for the Δ*phz*Δ*hcn* treatment because no Co(2+) measurements were carried out for the no wire controls based on the consistently low background Co(2+) detected in all other no wire control treatments.

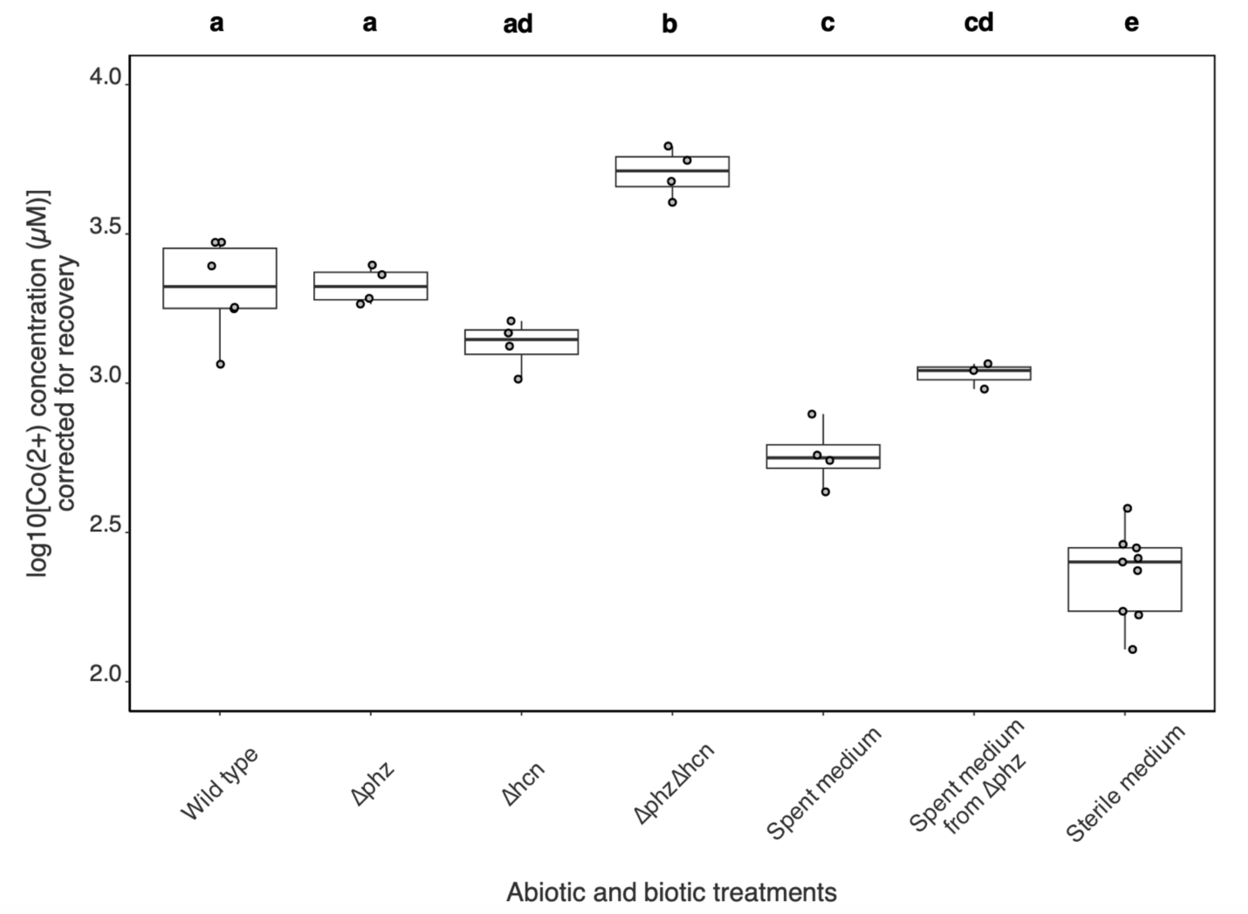


**Figure S2:** Boxplot summarizing one-way ANOVA of log10 transformed Co(2+) concentrations corrected for mass balance recoveries at 168 hours across biotic and abiotic treatments. The bottom and top of the boxes show the first and third quartiles, respectively, the bar in the middle shows the median and the whiskers represent 1.5 times the interquartile range. Type III sum of squares were used to account for the different sample numbers between treatments (Wild type: n = 6, Δ*phz*: n = 4, Δ*hcn*: n = 4, Δ*phz*Δ*hcn*: n = 4, spent medium: n = 4, spent medium from Δ*phz*: n = 3, sterile medium: n = 9). Abiotic and biotic treatment had a significant effect ($p < 0.05$) for the uncorrected ($p = 1.03 \text{ x } 10^{-11}$) and corrected data ($p = 1.93 \text{ x } 10^{-11}$). Letters not shared between boxes indicate significant differences between treatments according to the Tukey HSD test ($p < 0.05$).

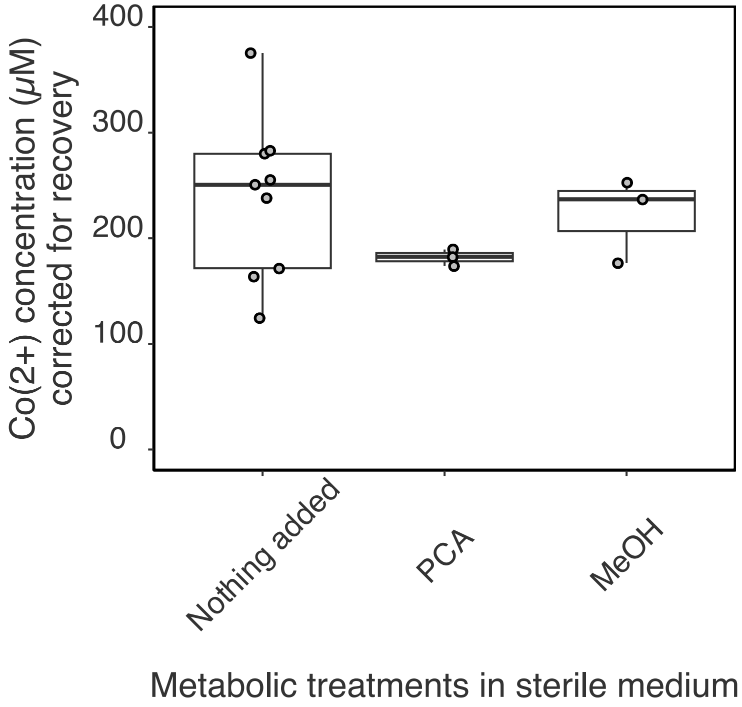


**Figure S3:** Boxplot summarizing one-way ANOVA of Co(2+) concentration at 168 hours corrected for mass balance recoveries across sterile medium, 50 µg/mL PCA, and 6.5% v/v methanol (MeOH) treatments. The bottom and top of the boxes show the first and third quartiles, respectively, the bar in the middle shows the median and the whiskers represent 1.5 times the interquartile range. Type III sum of squares were used to account for the different sample numbers between treatments (Nothing added: $n = 9$, PCA: $n = 3$, and MeOH: $n = 3$). Metabolite treatment had a p-value $> 0.05$ and did not have a significant effect on cobalt concentration.

2) Biosynthetic gene cluster analyses

**Bioinformatic analyses**

The full genome for *P. chlororaphis* subsp. *aureofaciens* generated using Illumina and Oxford Nanopore long-read sequencing was downloaded from ATCC via this link: https://genomes.atcc.org/genomes/60f7ef6772424c16. Genomes that were independently sequenced including the wild-type and deletion mutants (**Methods**) were also deposited to NCBI Bioproject PRJNA1336231. Samples were uploaded under biosample numbers SAMN52040257, SAMN52040258, and SAMN52040259, respectively. These samples were also uploaded to NCBI's SRA database under SRR35667903, SRR35667902, and SRR35667901. Due to interruption in the portal's service, we are also providing the genomes as **Supporting Data 2, Supporting Data 3,** and **Supporting Data 4,** and on the GitHub page associated with this work: https://github.com/carleton-envbiotech/Cobalt_corrosion/tree/main.

Biosynthetic gene clusters and their predicted metabolites were identified by loading the genome from ATCC directly into the web version of antiSMASH v8.dev-220ad8f9 adjusted to the "strict" setting (1). Predicted metabolites were verified by clicking through the top two best hits for each cluster's MIBiG database link, which is available through the graphic user interface in the web portal. Specific biosynthetic gene clusters of interest were manually exported as .svg files and edited in graphic software to highlight genetic loci of interest. The genome was also annotated using the Distilled Refined Annotation of Metabolism (DRAM) v.1.5.0 (2) using the KEGG and UniRef90 databases. The annotation file output by DRAM (**Supporting Data 1**) was used to manually search for any gene names containing the strings of text "cobalt", "cob", "cyanide", "cytochrome", "glutathione", "hcn", "hydrogenase", "peroxidase", "siderophore", and "super oxide dismutase" to identify pathways that could be contributing to oxidative stress response and cobalt cycling. Annotations from DRAM were cross-referenced with antiSMASH to confirm putative metabolites and transport pathways potentially involved in the MIC of cobalt. Visualizations were exported and manually edited in image editing software to add antiSMASH loci indicators for core biosynthetic genes and select transport-related genes that were independently annotated using KEGG and Uniref90 databases via DRAM (shown in **Table S2**).

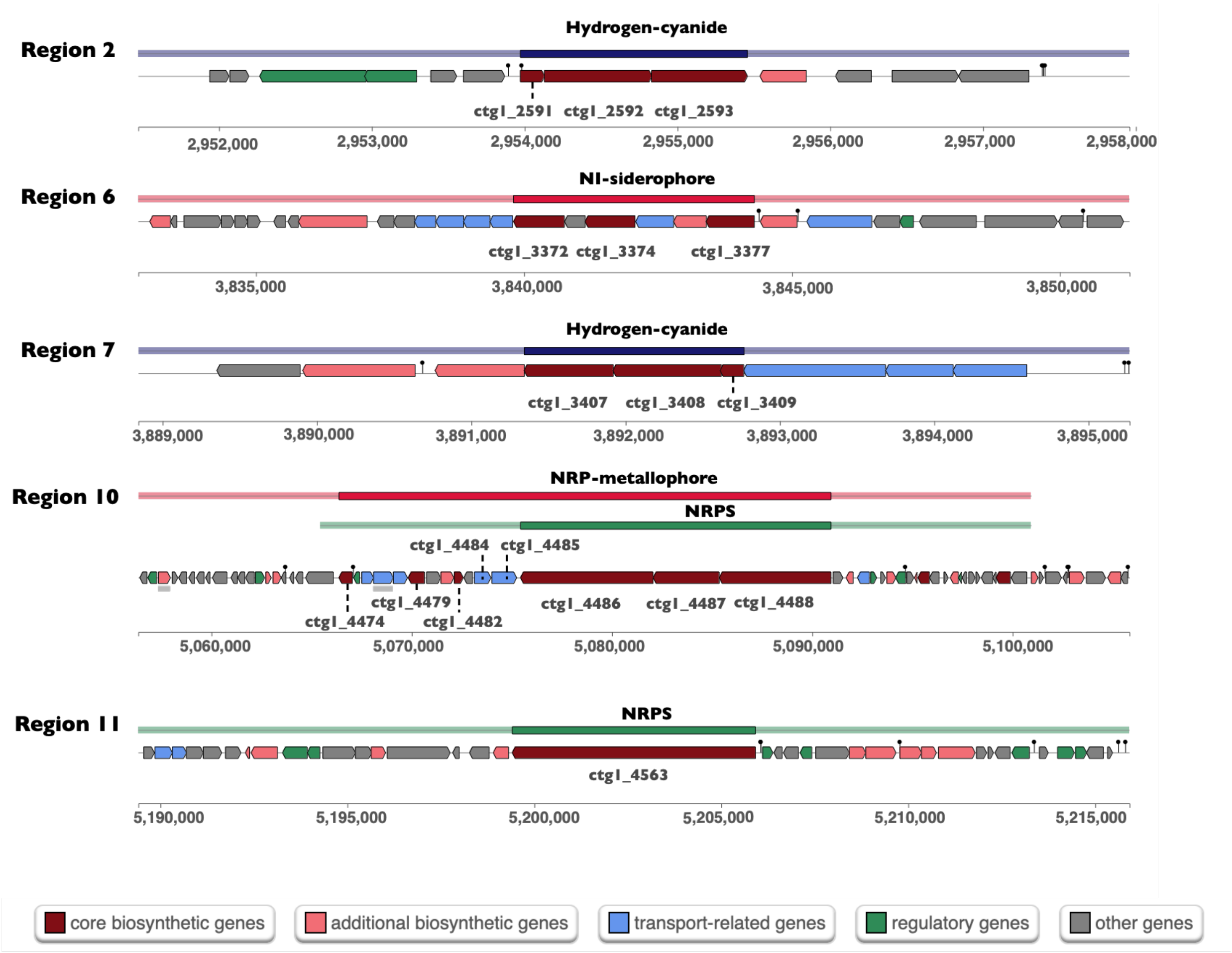


**Figure S4:** Summary of antiSMASH output for biosynthetic gene clusters predicted to produce metabolites relevant to cobalt solubilization and transport. For each region detected, the coloured line at the top indicates the span of genes associated with each biosynthetic cluster, where the shaded region highlights the core biosynthetic genes involved in the production of the metabolite titled above that region. The locations of these regions within wild-type *Pseudomonas chlororaphis* subspecies *aureofaciens* genome are indicated by numbered coordinates underneath the coloured gene clusters. Genes within these regions are further categorized by their roles in metabolite biosynthesis, as denoted by the legend at the bottom of the figure. Hashed lines were manually added to link identifiers to smaller genes (expanded in **Table S2**) in the gene clusters for clarity.

**Table S2:** DRAM annotations for core biosynthetic and select transport genetic loci within biosynthetic gene clusters predicted by antiSMASH. The start positions of each loci detected by antiSMASH were used to determine alignment with the start codons of the genes annotated in DRAM, providing unique identifier matches and insight into their specific functions.

| Region | antiSMASH identifier | Start position | End position | KEGG or UniRef90 ID | KEGG or Uniref90 hit |
|---|---|---|---|---|---|
| 2 | ctg1_2591 | 2956483 | 2956800 | K10814 | Hydrogen cyanide synthase HcnA |
| 2 | ctg1_2592 | 2956797 | 2958206 | K10815 | Hydrogen cyanide synthase HcnB |
| 2 | ctg1_2593 | 2958199 | 2959455 | K10816 | Hydrogen cyanide synthase HcnC |
| 6 | ctg1_3372 | 3846837 | 3848735 | K23375 | Staphyloferrin B synthase |
| 6 | ctg1_3374 | 3848740 | 3849513 | K23376 | Staphyloferrin B biosynthesis citrate synthase |
| 6 | ctg1_3377 | 3854022 | 3855821 | UniRef90_A0A0C5EL69 | AcsD protein |
| 7 | ctg1_3407 | 3893854 | 3895008 | UniRef90_A0AA45XIN0 | Sarcosine oxidase subunit beta |
| 7 | ctg1_3408 | 3895005 | 3896411 | UniRef90_A0A3G7K0E2 | Opine oxidase subunit A |
| 7 | ctg1_3409 | 3896393 | 3896701 | UniRef90_A0A5M7CTD1 | (2Fe-2S)-binding protein |
| 10 | ctg1_4474 | 5076434 | 5077768 | K10531 | L-ornithine N5-monooxygenase |
| 10 | ctg1_4479 | 5083355 | 5084977 | UniRef90_A0A554PAC0 | Twin-arginine translocation signal domain-containing protein |
| 10 | ctg1_4482 | 5087959 | 5088855 | UniRef90_Q4K997 | Chromophore maturation protein PvdO |
| 10 | ctg1_4484 | 5089961 | 5091613 | K06160 | Putative pyoverdin transport system ATP-binding/permease protein |
| 10 | ctg1_4485 | 5091714 | 5094182 | K16088 | Outer-membrane receptor for ferric coprogen and ferric-rhodotorulic acid |
| 10 | ctg1_4486 | 5094570 | 5107805 | UniRef90_A0A554PAB6 | Amino acid adenylation domain-containing protein |
| 10 | ctg1_4487 | 5107818 | 5114414 | UniRef90_A0A554PAC2 | Non-ribosomal peptide synthetase |
| 10 | ctg1_4488 | 5114414 | 5125576 | UniRef90_UPI002407C9DD | Amino acid adenylation domain-containing protein |
| 11 | ctg1_4563 | 5209456 | 5222472 | UniRef90_A0A554PEL7 | Non-ribosomal peptide synthetase |

**Table S3:** Predicted compounds produced by biosynthetic gene clusters identified by antiSMASH. Note that the top two best hits in the MIBiG database for each region are summarized for clarity.

| Region | Compound name and MIBiG link | Structure |
|---|---|---|
| 6 | Schizokinen (https://pubchem.ncbi.nlm.nih.gov/compound/3082425) | 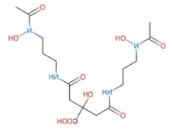 |
| | Vibrioferrin (https://mibig.secondarymetabolites.org/repository/BGC0000947.5/index.html#r1c1) | 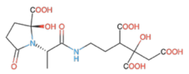 |
| 10 | Pf-5 pyoverdine (https://mibig.secondarymetabolites.org/repository/BGC0000413.5/index.html#r1c1) | 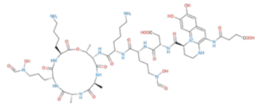 |
| | Pyoverdine SXM-1 (https://mibig.secondarymetabolites.org/repository/BGC0002693.2/index.html#r1c1) | 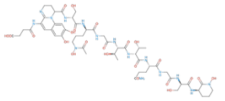 |
| 11 | Pf-5 pyoverdine (https://mibig.secondarymetabolites.org/repository/BGC0000413.5/index.html#r1c1) | 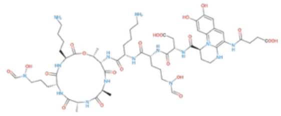 |
| | Azotobactin-D (https://mibig.secondarymetabolites.org/repository/BGC0002433.3/index.html#r1c1) | 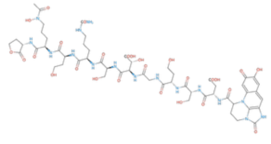 |

3) Cell-associated Co(2+) and toxicity measurements

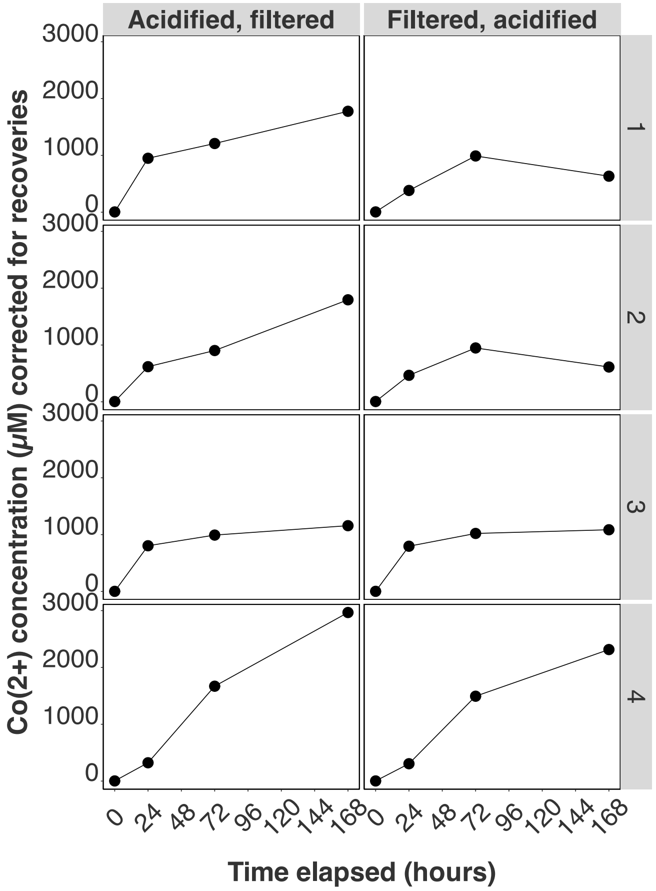


**Figure S5:** Co(2+) concentrations corrected for mass balance recoveries over 168 hours for paired samples that were acidified then filtered (n = 4) or filtered then acidified (n = 4) for wild type.

**Protocol for generating dose-response curve for Co(2+) exposure**

Dose-response curves were carried out to assess cobalt tolerance of *P. chlororaphis* subsp. *aureofaciens*. 10% v/v primary inoculums in defined medium were treated with Co(2+) standards made from $CoCl_2$ dissolved in ultra-filtered water to yield final concentrations of 0-500 µM. Triplicates of these solutions were transferred to a clear 96-well plate, alongside abiotic blanks of sterile defined medium matching the selected Co(2+) concentration levels for biotic samples. The microplate was incubated shaking at 26 ˚C, with $OD_{600}$ measurements recorded every 30 min for 96 hours. Dose-response curves were generated by calculating growth rates using the growthcurver v0.3.1 package (3) in R. The drc v3.0-1 (dose-response curve) package (4) was used to fit a five-parameter model to a dose-response curve using determined growth rates as a function of Co(2+) concentration. The 50 % inhibitory concentration (*i.e.,* IC50) value was obtained using the 'drc' package in R to provide a simplified metric assessing Co(2+) tolerance. Results are summarized in **Figure S6**.

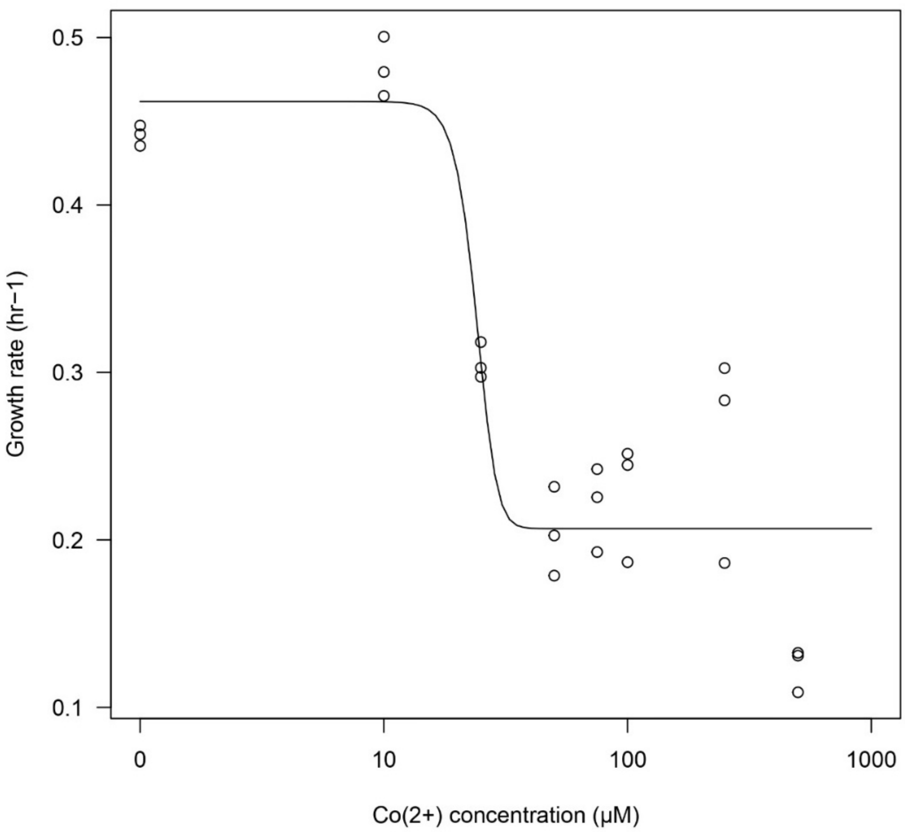


**Figure S6:** Dose response curve reporting growth rate of *P. chlororaphis* subsp. *aureofaciens* in response to treatment with Co(2+) concentrations ranging from 0-500 µM (presented on a log10 scale). Dose response curves were fitted by extracting growth rates using the growthcurver package in R and fitting a 5-parameter log-logistic equation using the drc package in R. Fitted IC50 values for growth rate ranged from 16.13 to 31.69 µM with a mean estimate of 23.9 µM.

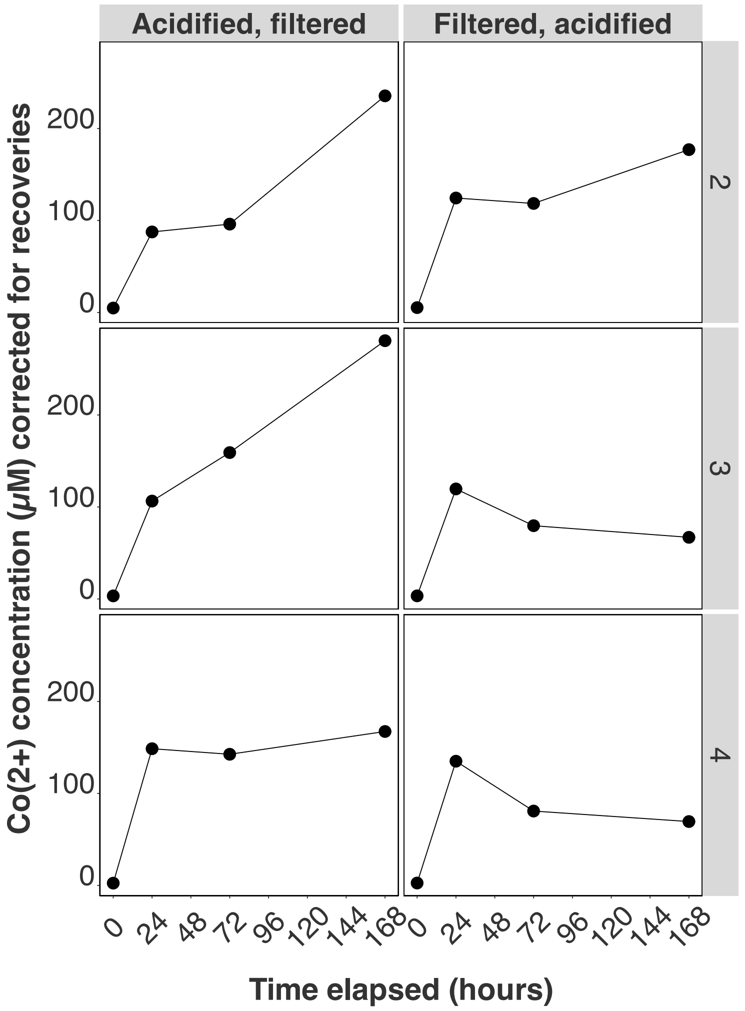


**Figure S7:** Co(2+) concentrations corrected for mass balance recoveries over 168 hours for sterile medium samples acidified then filtered (n = 3) or filtered then acidified (n = 3).

<u>4) Additional SEM images, wire diameters, and EDX data</u>

**Table S4:** Summary of diameters measured for representative wires presented in **Figure 4** using backscattered electron (BSE) detector. Note that the diameter for Δ*phz*Δ*hcn* was reported as a range due to corrosion resulting in inward curvature of the wire section imaged in **Figure 4I**.

| **Treatment** | **Diameter (µm)** |
|---|---|
| Sterile medium | 501.87 |
| Sterile medium anoxic | 498.63 |
| PCA | 498.05 |
| Spent medium | 474.89 |
| Spent medium from Δ*phz* | 472.51 |
| Spent medium from Δ*phz* anoxic | 501.62 |
| Wild type | 441.64 |
| Δ*phz* | 420.09 |
| Δ*hcn* | 461.46 |
| Δ*phz*Δ*hcn* | 333.71 to 359.79 |

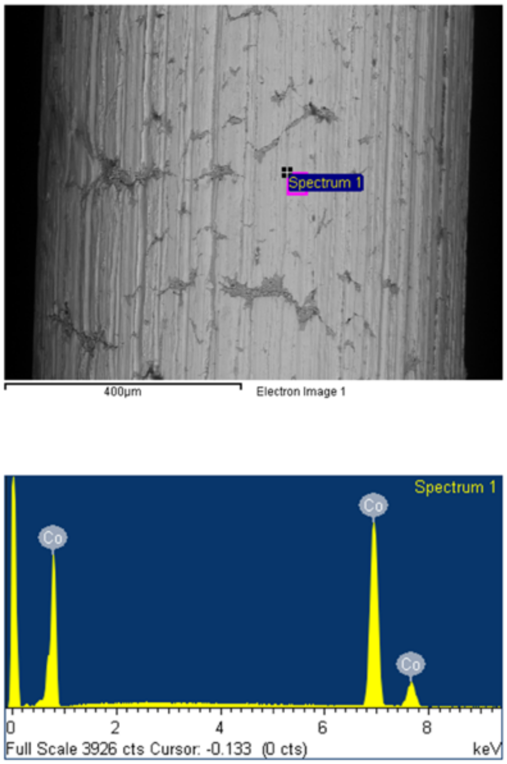


**Figure S8:** EDX scan of representative unmanipulated wire using BSE detector.

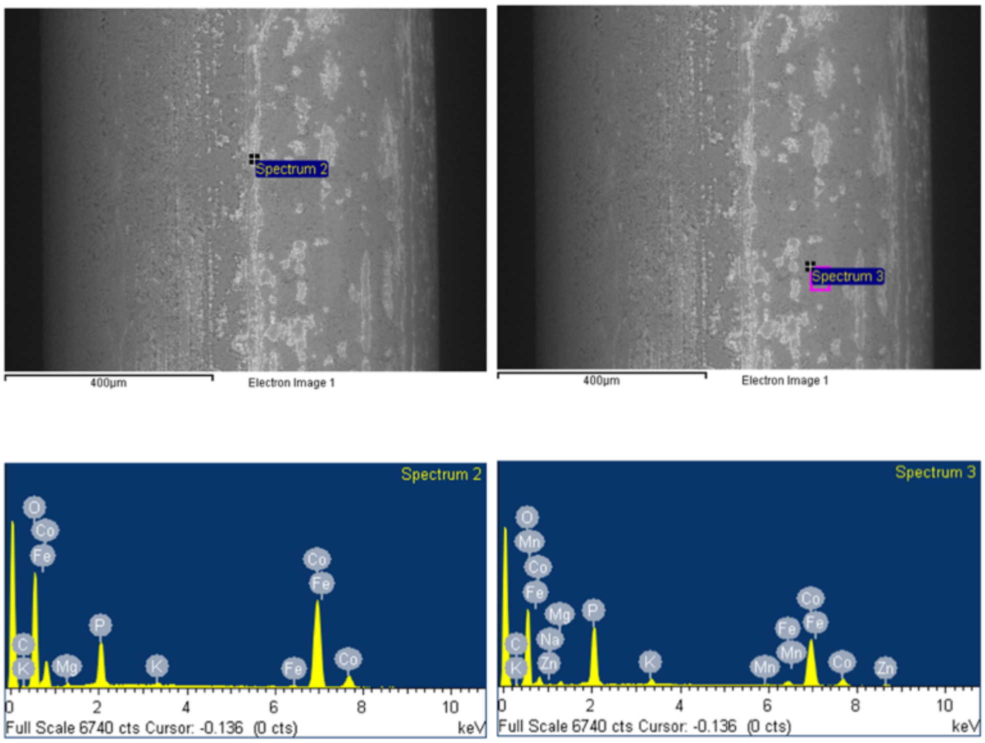


**Figure S9:** EDX scans of representative wire from sterile medium oxic treatment using BSE detector.

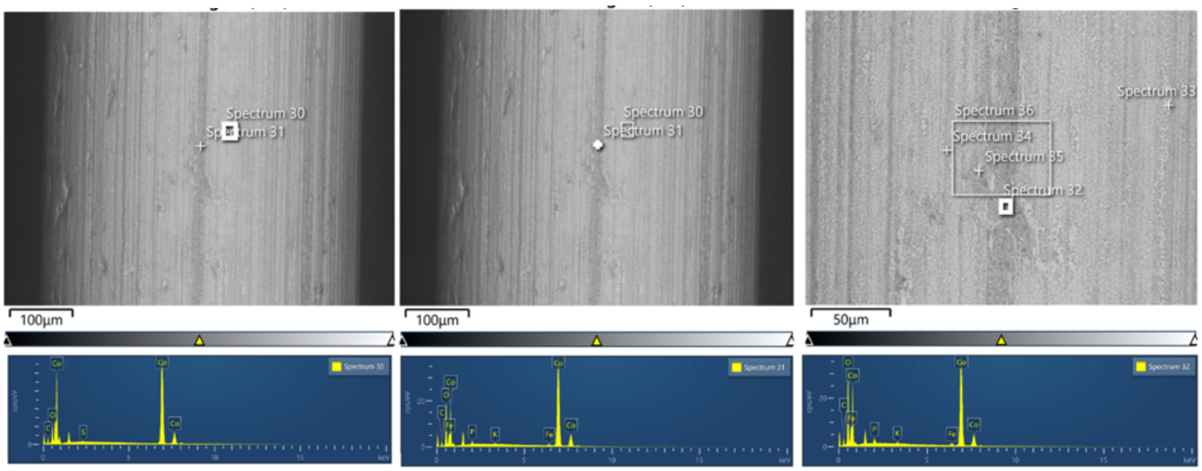


**Figure S10:** EDX scans of representative wire from sterile medium anoxic treatment using BSE detector.

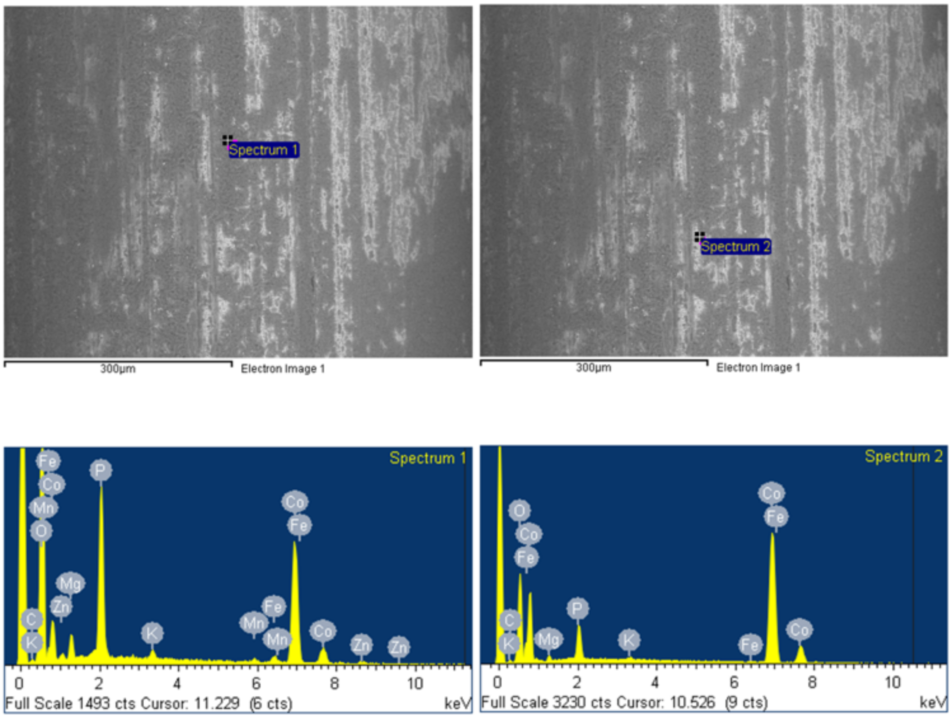


**Figure S11:** EDX scans of representative wire from PCA treatment using BSE detector.

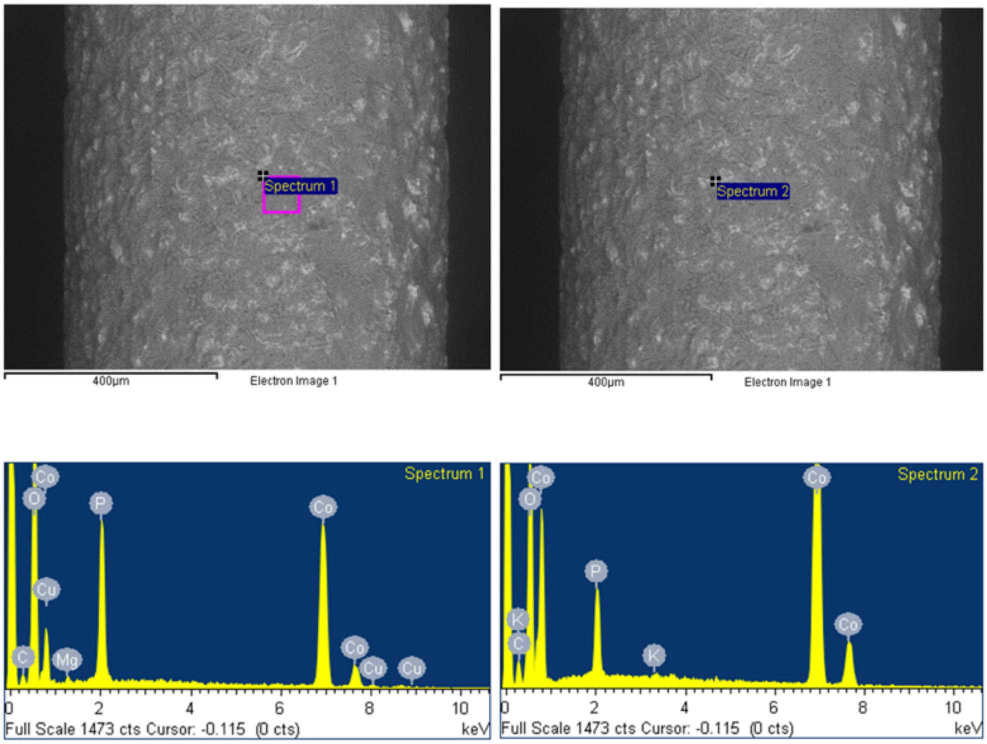


**Figure S12:** EDX scans of representative wire from spent medium treatment using BSE detector.

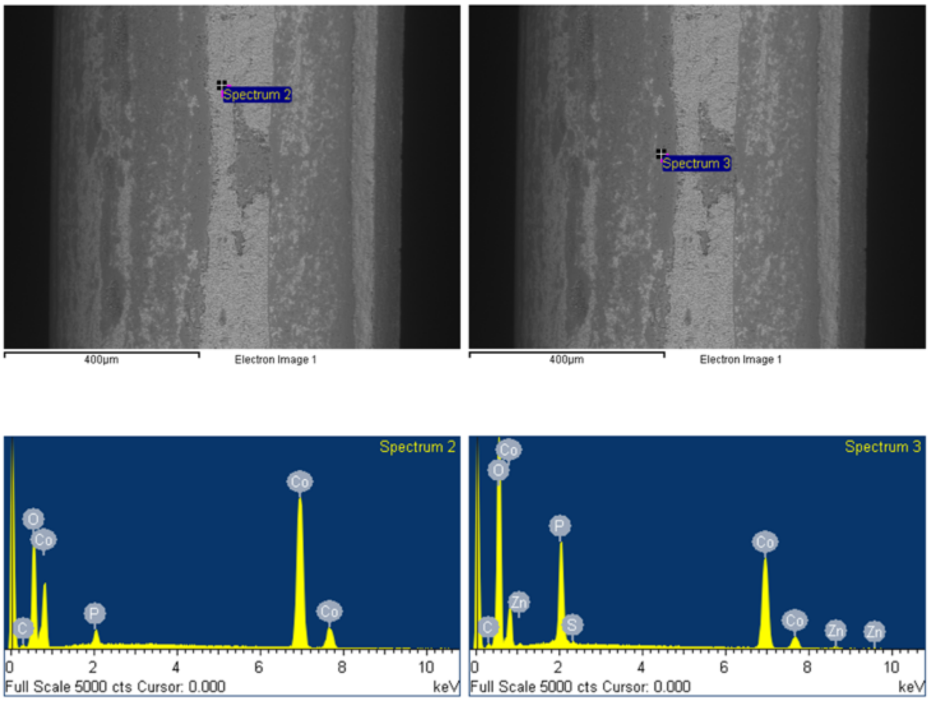


**Figure S13:** EDX scans of representative wire from spent medium from Δ*phz* oxic treatment using BSE detector.

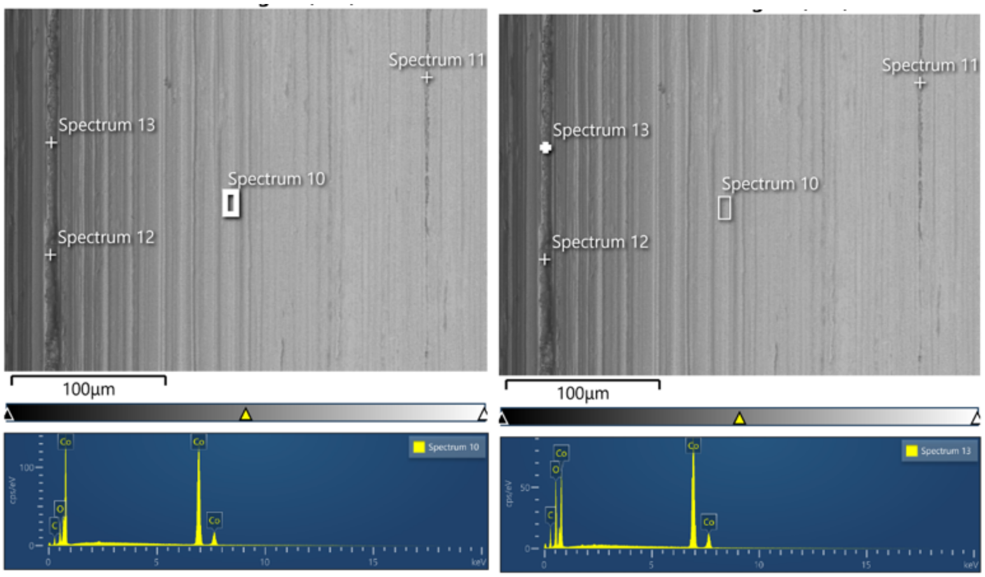


**Figure S14:** EDX scans of representative wire from spent medium from Δ*phz* anoxic treatment using BSE detector.

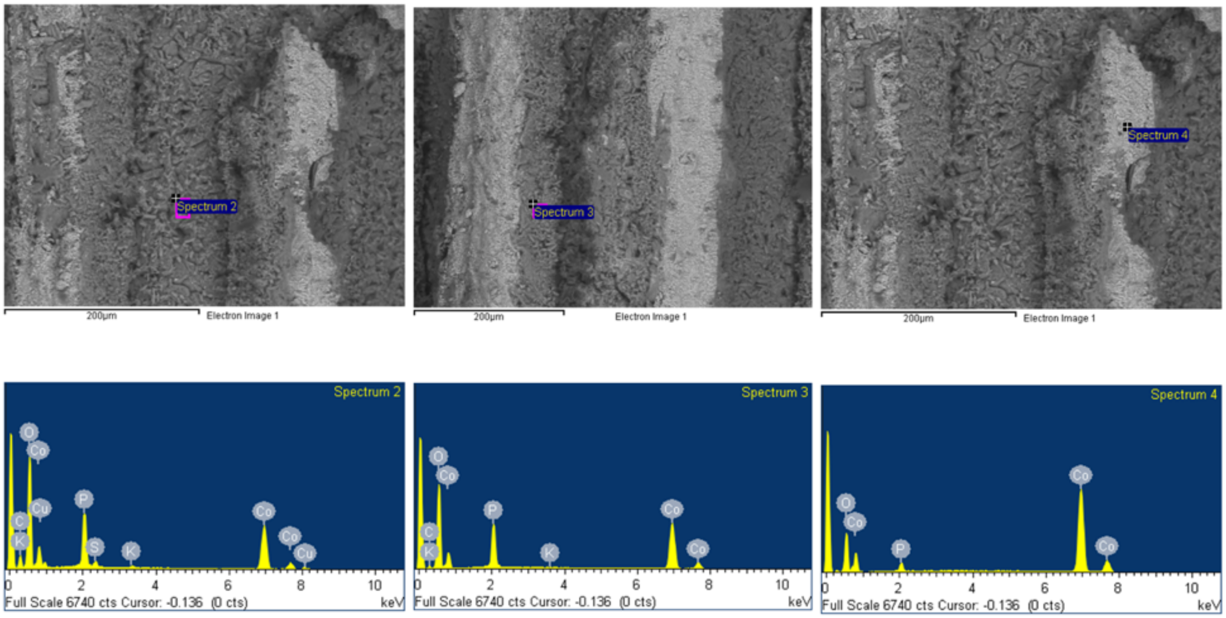


**Figure S15:** EDX scans of representative wire from wild type treatment using BSE detector.

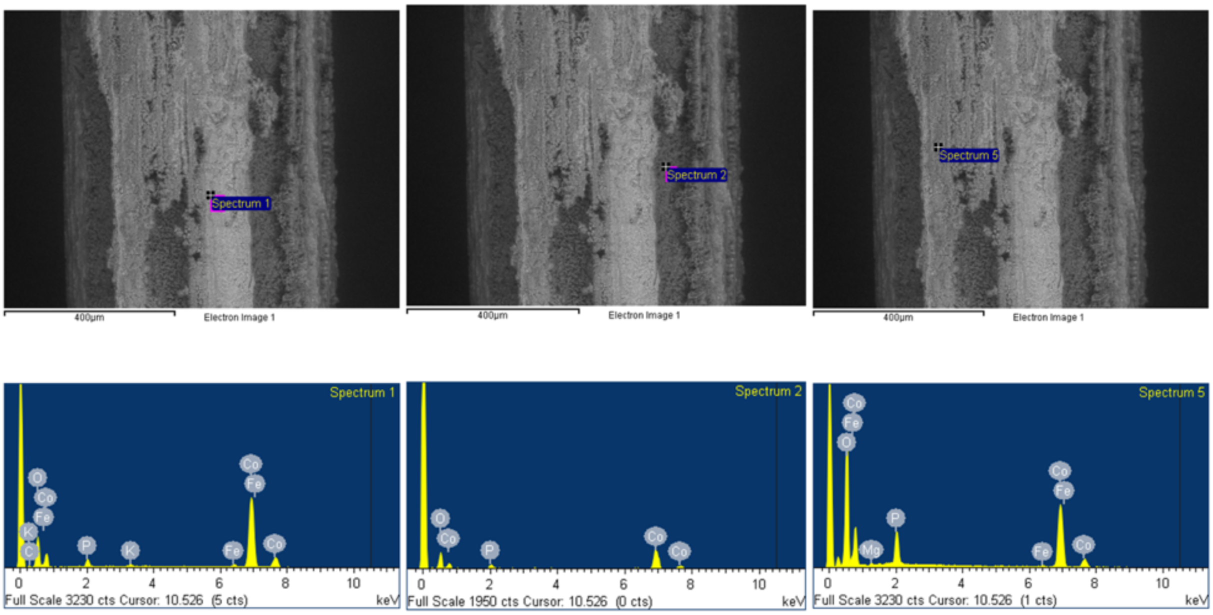


**Figure S16:** EDX scans of representative wire from Δ*phz* treatment using BSE detector.

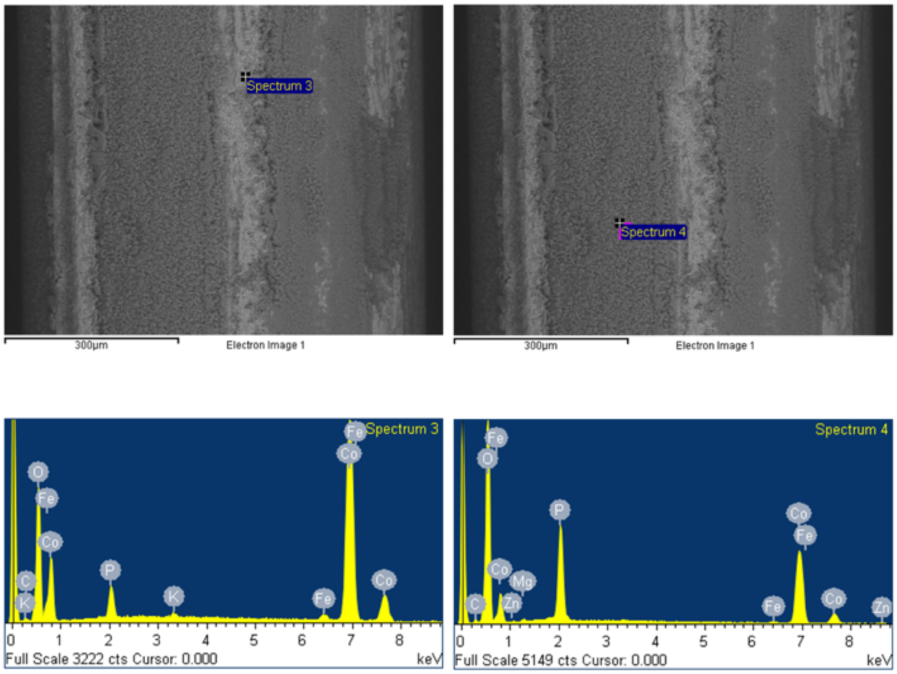


**Figure S17:** EDX scans of representative wire from Δ*hcn* treatment using BSE detector.

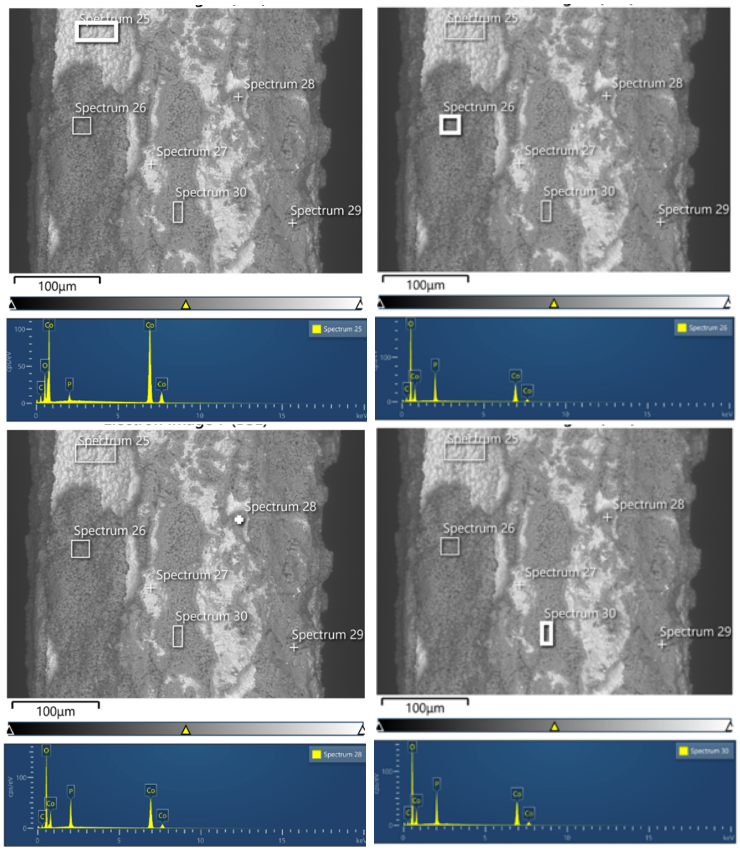


**Figure S18:** EDX scans of representative wire from Δ*phz*Δ*hcn* treatment using BSE detector.

**Table S5:** Percent weight surface composition of cobalt and oxygen from selected EDX spectra of representative wires presented in **Figures S8-S18**. Selection of scanning spectrum areas was driven by the shading of different areas of the wire shown by the BSE detector, comparing darker areas indicative of oxidized regions to lighter areas of greater Co(0) content. Rough estimates of Co:O ratios were calculated by dividing the percent weight of cobalt over that of oxygen from the EDX scans. Co:O ratios displayed as <1:1 or ~1:1 are highlighted in yellow and represent areas of potential cobalt oxide formation, with approximate makeup providing insight into possible cobalt oxides present.

| **Treatment** | **Spectrum** | **Co (%)** | **O (%)** | **Co:O ratio** |
|---|---|---|---|---|
| Unmanipulated | 1 | 100 | 0 | N/A |
| Sterile medium | 2 | 53.29 | 35.84 | 1.49:1 |
| | 3 | 39.01 | 37.27 | 1.05:1 |
| Sterile medium anoxic | 30 | 83.92 | 5.61 | 14.96:1 |
| | 31 | 79.81 | 13.01 | 6.13:1 |
| | 32 | 69.70 | 21.00 | 3.32:1 |
| PCA | 1 | 32.82 | 47.20 | 0.70:1 |
| | 2 | 66.76 | 24.21 | 2.76:1 |
| Spent medium | 1 | 38.88 | 43.06 | 0.90:1 |
| | 2 | 63.4 | 25.50 | 2.49:1 |
| Spent medium from Δ*phz* | 2 | 68.65 | 25.89 | 2.65:1 |
| | 3 | 36.05 | 47.78 | 0.75:1 |
| Spent medium from Δ*phz* anoxic | 10 | 85.99 | 4.98 | 17.27:1 |
| | 13 | 69.29 | 14.69 | 4.72:1 |
| Wild type | 2 | 29.26 | 46.45 | 0.63:1 |
| | 3 | 42.7 | 42.34 | 1.01:1 |
| | 4 | 79.1 | 18.62 | 4.25:1 |
| Δ*phz* | 1 | 74.87 | 17.79 | 4.21:1 |
| | 2 | 67.35 | 28.96 | 2.33:1 |
| | 5 | 47.26 | 44.65 | 1.06:1 |
| Δ*hcn* | 3 | 69.11 | 23.65 | 2.92:1 |
| | 4 | 34.85 | 48.68 | 0.72:1 |
| Δ*phz*Δ*hcn* | 25 | 78.02 | 10.21 | 7.64:1 |
| | 26 | 30.96 | 49.39 | 0.63:1 |
| | 28 | 44.24 | 40.99 | 1.08:1 |
| | 30 | 36.76 | 44.90 | 0.82:1 |

5) Statistical analyses of ICP-MS data and SEM of anoxic and DMSO treatments

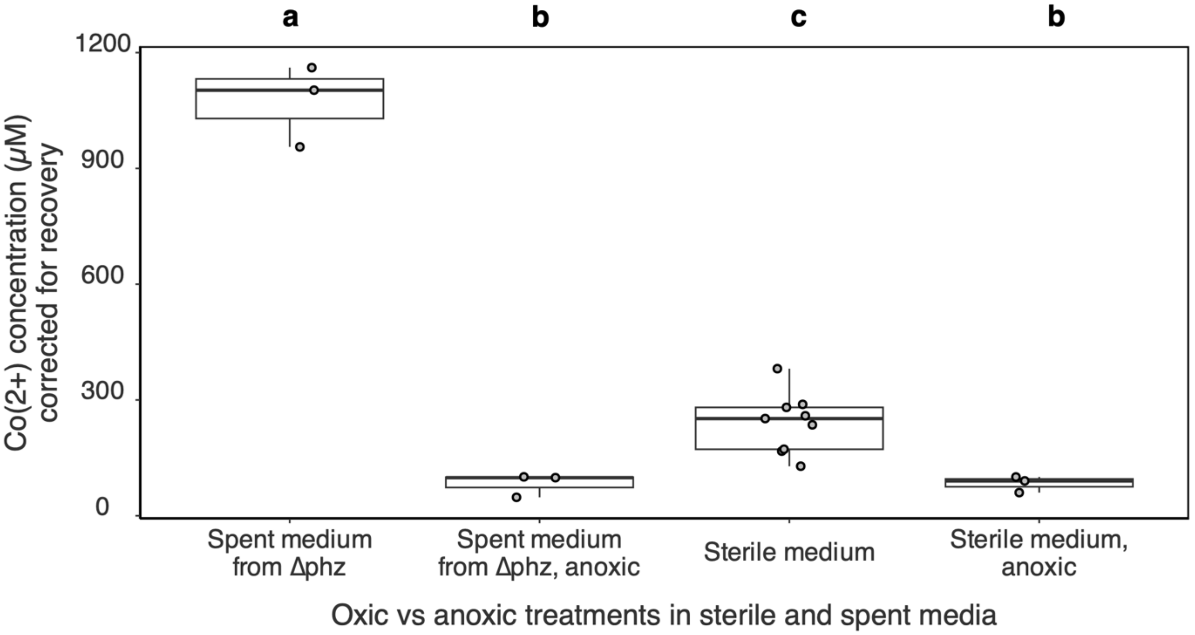


**Figure S19:** Boxplot summarizing one-way ANOVA of Co(2+) concentration corrected for mass balance recoveries at 168 hours across oxic and anoxic treatments. The bottom and top of the boxes show the first and third quartiles, respectively, the bar in the middle shows the median and the whiskers represent 1.5 times the interquartile range. Type III sum of squares was used to account for the different sample numbers between treatments (Spent medium from Δ*phz*: n = 3, Spent medium from Δ*phz*, anoxic: n = 3, Sterile medium: n = 9, Sterile medium, anoxic: n = 3). The different combinations of oxic and anoxic treatments and spent media and sterile media had a significant effect ($p < 0.05$) on mean cobalt concentrations ($p = 1.36 \times 10^{-10}$). Letters not shared between boxes indicate significant differences between treatments according to the Tukey HSD test ($p < 0.05$).

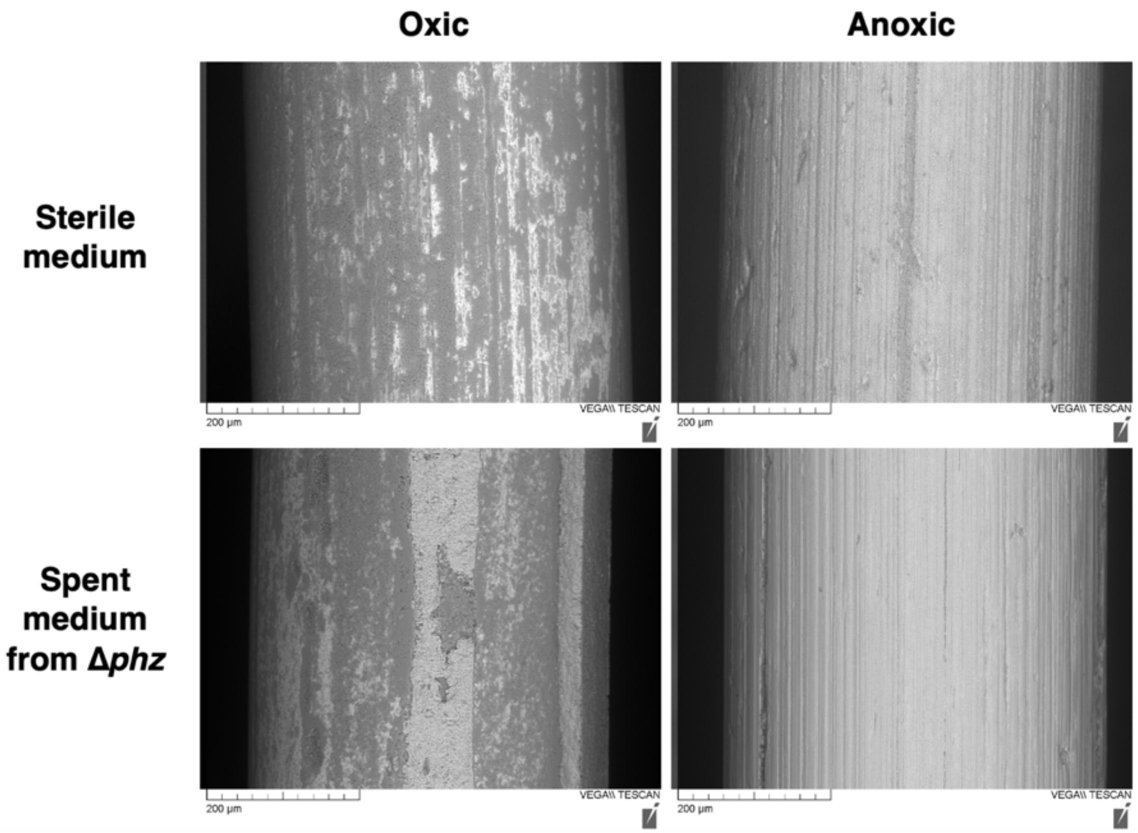


**Figure S20:** SEM images of Co(0) wires in oxic and anoxic conditions for sterile medium and spent medium from Δ*phz* treatments after 168 hours. Wires were magnified to 400X and images obtained using the BSE detector are displayed.

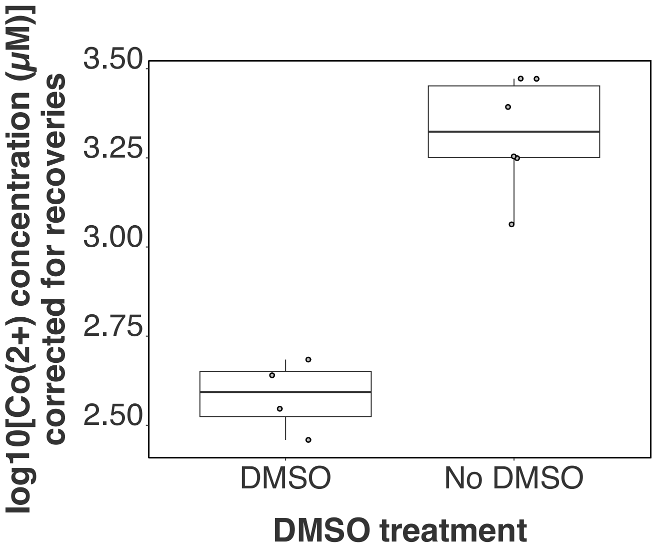


**Figure 21:** Boxplot summarizing one-way ANOVA of log10 transformed Co(2+) concentrations corrected for mass balance recoveries at 168 hours across wild-type treatments with and without DMSO. The bottom and top of the boxes show the first and third quartiles, respectively, the bar in the middle shows the median and the whiskers represent 1.5 times the interquartile range. ANOVA using Type III sum of squares was used to account for the different sample numbers between treatments (DMSO: $n = 4$, no DMSO: $n = 6$). The addition of DMSO significantly lowered ($p < 0.05$, $p = 0.00056$) cobalt concentration compared to treatments that did not receive DMSO. No multiple comparison tests were carried out because only two treatment levels were included.

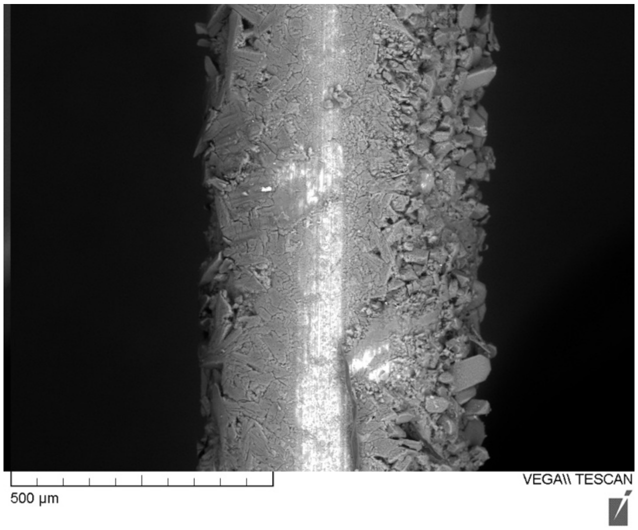


**Figure S22:** Representative SEM image of Co(0) wires from sterile DMSO treatment after 168 hours, taken at 200X magnification using the BSE detector. Formation of large crystals can be seen, leading to an increased diameter of 551.41µm (compared other sterile wires ranging closer to the manufacturer listed diameter of 500µm).

7) Redox reaction comparison between Fe & Co

**Table S6:** Free energy values at standard and non-standard conditions (*i.e.;* $\Delta G^0$ and $\Delta G$, respectively) for possible redox reactions contributing to the oxidation of cobalt. Mean concentrations for starting Co(0), recovered Co(2+), and concentration of spiked-in PCA ($PCA_{ox}$) were taken from the PCA experimental treatment. Conversion of Co(0) to Co(2+) coupled to PCA reduction was compared to abiotic oxidation driven by dissolved $O_2$. Dissolved [$O_2$] was determined using the partial pressure of $O_2$ in the atmosphere within the flask and Henry's Law constant for $O_2$ in water at 28˚C. These free energies were compared to the oxidation of iron by the same oxidants in standard and non-standard conditions.

| **Overall redox reaction & concentrations** | **$\Delta G^0 = -nFE^0_{cell}$ (kJ/mol)** | **$\Delta G = \Delta G^0 + RTlnQ$ (kJ/mol)** |
|---|---|---|
| Co(0) + $PCA_{ox}$ → Co(2+) + $PCA_{red}$<br><br>Recovery correct mean [Co(2+)] = 185.52 µM<br>[$PCA_{red}$] = [Co(2+)] = 185.52 µM<br>[Co(0)] = 6186.66 µM – [Co(2+)] = 6001.14 µM<br>[$PCA_{ox}$] = 223.005 µM – [$PCA_{red}$] = 37.49 µM | -32.04 | -36.74 |
| Fe(0) + $PCA_{ox}$ → Fe(2+) + $PCA_{red}$<br><br>[Fe(2+)] = 185.52 µM<br>[$PCA_{red}$] = [Fe(2+)] = 185.52 µM<br>[Fe(0)] = 6186.66µM – [Fe(2+)] = 6001.14 µM<br>[$PCA_{ox}$] = 223.005µM – [$PCA_{red}$] = 37.49 µM | -62.53 | -67.23 |
| Co(0) 1/2$O_2$ + 2$H^+$ → Co(2+) + $H_2O$<br><br>[Co(2+)] = 185.52 µM<br>[$H_2O$] = [Co(2+)] = 185.52 µM<br>[Co(0)] = 6186.66 µM – [Co(2+)] = 6001.14 µM<br>[$O_2$] = 249.04 µM – [$H_2O$] = 63.52 µM | -288.90 | -294.92 |
| Fe(0) 1/2$O_2$ + 2$H^+$ → Fe(2+) + $H_2O$<br><br>[Fe(2+)] = 185.52 µM<br>[$H_2O$] = 185.52 µM<br>[Fe(0)] = 6186.66 µM – [Fe(2+)] = 6001.14 µM<br>[$O_2$] = 249.04 µM – [$H_2O$] = 63.52 µM | -319.11 | -325.13 |

8) Genetic manipulations and validation

### Detailed protocols and background used in deletion mutant generation

Geneious was used as a visualization software to design the Δ*phz* and Δ*hcn* deletion alleles and validation primers. SacI and XmaI restriction sites were chosen since they are unique in the multiple cloning site (MCS) of the recipient plasmid pK18msB, absent from the insert sequence, and compatible in double digestion reactions. Additional random 6 bp buffer sequences were added exterior to the restriction sites to stabilize the restriction enzyme and DNA interactions. The synthetic gene fragment encoding the deletion alleles (**Figures S23 & S24**) were ordered from Twist Bioscience, resuspended in nuclease-free water to 20 μg/mL, and stored at -20 ˚C. For each operon, a pair of 20 bp primers flanking the deletion junction was designed to validate the presence of the deletion allele in mutants (sequences in **Table 1** of Main Text).

To purify pK18msB (177839 Addgene) from *E. coli* DH5 stab culture, cells were streaked onto Luria-Bertani (LB) agar plates amended with kanamycin (Kan). Solid LB medium was made by combining 5 g tryptone, 2.5 g yeast extract, 5 g NaCl, and 7.5 g of agar in 500 mL of MilliQ water. After autoclaving at 121 ˚C for 30 min, Kan was added to a final concentration of 30 μg/mL. Streak plates were incubated at 37 ˚C for 17hr, and single colonies were picked to generate primary inoculums in LB + 30 μg/mL Kan liquid medium shaking at 220 rpm and 37 ˚C overnight. pK18msB was purified using EZ-10 Spin Column Plasmid DNA Minipreps Kit (BS414 BioBasic). Volumes of 3 mL and 5 mL of primary inoculums were used, with 3 mL determined as optimal for this miniprep kit. DNA recovered was quantified using the Qubit™ dsDNA Broad Range Assay Kit (Q32853 Invitrogen) with a Qubit3 Fluorometer (Invitrogen).

Restriction digestions were conducted enable ligation of deletion inserts to the pK18msB vector. 50 μL digestion reactions with SacI-HF and XmaI (R3156S & R0180S New England Biolabs) were prepared with DNA inputs of 1 μg purified pK18msB or 333 ng insert DNA. Single digestions of pK18msB with each restriction enzyme and undigested plasmid were run as controls. All reactions were incubated at 37 ˚C for 1 hour, 65 ˚C for 20 min, and held at 12 ˚C in a Bio-Rad T100 Thermal Cycler. Products from the double digestions were purified using EZ-10 Spin Column PCR Products Purification Kit (BS364 BioBasic). Purified products were quantified and visualized alongside controls by gel electrophoresis to confirm fragment sizing and enzyme activity. Ligations between digested pK18msB and digested Δ*phz* or Δ*hcn* inserts were performed with T4 DNA Ligase (M0202S New England Biolabs). The standard protocol was halved for final reaction volumes of 10 μL and a molar ratio of 3:1 for insert to plasmid was chosen. Correcting for size, 73.7 ng of Δ*phz* and Δ*hcn* inserts and 100 ng pK18msB were added to the ligation reaction which was incubated at 16 ˚C for 16 hours, 65 ˚C for 10 min, and then held at 4 ˚C in a Bio-Rad T100 Thermal Cycler. A reaction containing water instead of the Δ*phz* insert was included as a 'no insert' ligation control.

Ligation products were transformed into *E.coli* NEB5-alpha competent cells (C2988J New England Biolabs) using a standard heat-shock transformation protocol. 2 μL of ligation reactions were added to 50 μL of thawed cells. Positive and negative transformation controls included 0.5 μL of undigested vector or no input, respectively. Transformation mixtures were incubated on ice for 30 min, subjected to 90 sec of heat shock at 42 ˚C, and immediately chilled on ice. Following addition of 1mL of LB, transformation mixtures were incubated for 1.5 hours at 37 ˚C with shaking, and 150 μL of the undiluted mixtures were spread onto LB + 30 μg/mL Kan plates.

Overnight growth at 37 ˚C yielded KanR transformants that had uptaken pK18msB containing the KanR gene.

Selected transformants carrying pK18msB with the Δ*phz* insert were identified by colony PCR with primers homologous to insert-specific sequences. 20 µL PCR reactions were prepared using Invitrogen Platinum II Hot Start PCR Master Mix (14000012 ThermoFisher Scientific). Sterile pipette tips were used to pick KanR colonies, streak onto LB + 30 µg/mL Kan plates for isolation, and inoculate the PCR reaction tubes with cells as the source of template DNA. 1 µL of the unmanipulated Δ*phz* insert DNA obtained from Twist Bioscience was used as a positive control, and no template was added for the negative control. Reactions were incubated in the thermal cycler at 94 ˚C for 10 min, followed by 30 PCR cycles programmed at 94 ˚C for 15 sec, 60 ˚C for 15 sec, and 68 ˚C for 30 sec. Reactions were held at 12 ˚C until storage at -20 ˚C.

PCR products were visualized by gel electrophoresis with the voltage was set to 90 V. Colonies yielding 1.4 kb PCR products, corresponding to the insert size, were recovered from streak plates and inoculated in LB + 30 µg/mL Kan liquid medium. Cultures were incubated at 37 ˚C overnight and stored as -80 ˚C cryostocks in 1:1 ratio with 50% filter-sterilized glycerol solution. Cryostocks were streaked on selective media to confirm activity of the KanR and SacB selectable markers present on pK18msB. Growth on LB + 30 µg/mL Kan + 5% sucrose agar confirmed sucrose-susceptibility conferred by the *sac*B genes, with kanamycin included in the sucrose test medium to prevent loss of the plasmid.

Tri-parental matings were conducted to introduce the pK18msB–Δ*phz* and pK18msB–Δ*hcn* vectors from *E. coli* NEB5-α transformants to the wild-type *P. chlororaphis* subsp. *aureofaciens*. Independent *E. coli* NEB5-α transformants (named Δ*phz*3 and Δ*phz*10) were the donor strains and the wild-type pseudomonad was the recipient. *E. coli* HB101 containing plasmid pRK2013 (37159 ATCC) was included to provide necessary conjugation functions *in trans* encoded by the pRK2013 helper plasmid (5). Primary inoculums of the donor and helper were prepared by inoculating from cryostocks into LB + 30 µg/mL Kan liquid medium and incubating at 37 ˚C overnight. A primary inoculum of the recipient was prepared by inoculating a cryostock into LB liquid medium and incubating at 28 ˚C overnight. Recipient cultures were incubated at 42 ˚C for 1 hour prior to setting up the conjugation reactions to inactivate host nucleases and increase efficiency (6).

Conjugations were prepared by washing cells via centrifugation and resuspension in LB, and combining washed cells in a 2:2:1 ratio of donor to helper to recipient cultures. This ratio was selected to ensure viability of the donor and helper strains after a failed conjugation with a 1:1:2 ratio, where the *E. coli* strains were likely killed by the pseudomonad. Negative controls included matings using unmanipulated pK18msB from Addgene as the donor (to confirm requirement of Δ*phz* or Δ*hcn* inserts), lacking the helper (to confirm requirement of helper plasmid), and lacking both the donor and helper (to confirm antibiotic susceptibility of the recipient). Mixtures were centrifuged at 10,000 xg for 3 min, and the supernatant was discarded to leave 30-50 µL. Pellets were resuspended in this volume and spotted into LB plates that were incubated at 30 ˚C overnight.

The following selection process is based on published protocols conducted by Liu et al (7). Conjugation spot colonies were scraped with sterile pipette tips into 0.5 mL LB liquid medium. After vortexing to resuspend cells, the mixtures were centrifuged at 10,000 xg for 3 min and pellets were resuspended in remaining 100 µL of the supernatant. These volumes were spread onto LB + 50 µg/mL Kan + 100 µg/mL Amp plates and incubated at 28 ˚C for 48 hours. These plates were designed to select for merodiploid conjugates that underwent homologous recombination with pK18msB-Δ*phz*, since the plasmid is incapable of replication in the recipient strain. The successful recombinants should be resistant to kanamycin due to the presence of the KanR gene encoded by pK18msB. The ampicillin would kill off the donor and helper strains, while permitting growth of the pseudomonad recipient due to its intrinsic AmpR. The presence of colonies on the selective medium, coupled with the lack of colonies associated with the negative controls indicated that the conjugations were successful. Selected colonies were streaked for isolation on LB + 50 µg/mL Kan + 100 µg/mL Amp plates and incubated at 28 ˚C for 48 hours. Sterile pipette tips were used to streak biomass from the isolation plates onto LB + 10% sucrose plates that were incubated at 28 ˚C for 30 hours. These plates selected for KanR conjugates that underwent a second homologous recombination event to excise the integrated pK18msB by *sac*B-mediated sucrose counterselection, which is expected to yield a mixture of successful deletion mutants and wild-type revertants.

Single colonies from the sucrose selection plates were streaked for isolation onto new LB + 10% sucrose plates and incubated at 28 ˚C for 72 hours, and a single SucR colony from each of these isolation streaks was screened by colony PCR to identify Δ*phz* or Δ*hcn* deletion mutants (**Figures S25 & S26**). Colonies of candidate deletions were streaked for isolation on LB + 10% sucrose, inoculated in LB broth, and stored as cryostocks. Phenotypic validation of the candidate deletion mutants was performed by streaking on LB alone, LB + 50 ug/mL Kan, and LB + 10% sucrose plates (**Figure S28**). Successful deletions should lack the KanR and *sac*B genes from pK18msB, and therefore were predicted to not grow on the kanamycin plates, but grow on the sucrose plates.

Both genotypic and phenotypic validation confirmed strains Δ*phz*3.1, Δ*phz*3.3, and Δ*phz*10.4 as successful *phz* operon deletion mutants, and strains Δ*hcn*10.3, Δ*hcn*14.1, Δ*hcn*14.2, and Δ*hcn*14.4 as successful *hcn* operon deletion mutants. Streaks of each of the validated Δ*phz* mutants lacked the distinctive orange colour of the wild type as shown in **Figure S28**. Finally, a Δ*phz*Δ*hcn* double mutant was constructed, in which the Δ*phz* deletion allele was sequentially introduced into *P. chlororaphis* subsp. *aureofaciens* Δ*hcn*14.1. Confirmed strains Δ*phz*Δ*hcn*3.3, Δ*phz*Δ*hcn*3.4, Δ*phz*Δ*hcn*10.3, and Δ*phz*Δ*hcn*10.4 showed presence of the Δ*phz* allele (**Figure S27**) and lacked the orange colouration indicative of phenazine production (**Figure S28**). The validated strains Δ*phz*10.4, Δ*hcn*14.1, and Δ*phz*Δ*hcn*3.3 were selected for cobalt corrosion tests. Strains Δ*phz*10.4 and Δ*hcn*14.1 were selected for whole genome sequencing and confirmed to lack their respective operons (**Methods** and **Figure S29**). No notable differences in growth were observed between cultures of the wild-type and deletion strains grown in defined medium, when comparing $OD_{600}$ values after 16 hours shaking at 28 ˚C or via viable plate counts (**Figure 3**)

>phz_operon_insert_buffer_sequence Pseudomonas chlororaphis subsp. aureofaciens ATCC® 13985™, contig 1
ACTGAGGAGCTCGATCGATCACCGCGTAGTTAGCCGCCTGATATCGCTGCACCCAGTC
CTCGGGGTAATTGCCGTACATATAAGTCCTGGGCCGCATGAACGGGGTCACGCTGCAC
ATGCCGTAGGCAAAAAAATCAAACCGTAGTTCGCGCAAGGCCCTCAACGCGACAGC
CGTAAACTCCTGCATATCCATGGTTCGCGCAAAGATACTATAAAAGTACGCATCCCATC
CCAACTGCTGCCCTAATTCCATTTTGAGCACCACTAAAGTTGAAAACAGGCCGTTAGA
CTAGACCAACCTGAATCTCTGTCAACAAGCAAAATCACATCGTCATAGAAGGCTTTGC
GCTAGCACTAGATTCATTGCGCCTAATCCAACCAACTACTTAGAACATTGCATCCCGG
CGATGTTAGTAAGCCATTACCGCCTCTCAGCCGACGAATGAACCTGCCTCTTCACAAT
AATGCAACAGTTCATTCGGTGGGCTGCAGCCAACCCGTTCATACGCTAATCACTACAA
GATCTGGTAGTTCCACCCCCAAGAAACGCAGGTGTATAAACAACACCCGCTCAGCAC
CGCCACGCAGGAACATAGTTAAACACTGTAAATATTCACCCACGTCAATTAACTAAAT
ATTTACTTTCAACATTATCACCCCCCAACTAAGGAGGATGCTGCCCATGCCTGCTTCGC
TTTCCCCTAGCGGCTTTGGCTGGAAACATCGCCGATTACAGCCGTAGGGTACCGAGAT
AAATATGCTTTGAAGTGCTGGCTGCTCCAACTTCGAACTCATTGCGCGAACTTCAACA
CTTATGACACCCGGTCAACATGAGAAGAGTCCAGATGCGAAAGAACGCGTATTCGAA
ATACCAAACAGAGAGTCCGGATCACCAAAGTGTGTAACGACATTAATTCCTATCTGAA
TCTTATAGTTGCTCTAGAACGTTGTCCTTGACCCAGCGATAGACATCGGGCCAGAGAC
TACACAAACAAAGTTAGACATTACTGAGGCTGCTACCATGCTAGATCTTCAAAACAAG
CGTAAATATCTGAAAAGTGCAGAATCCTTCAAAGCTTCACTGCGTGATGACCGCACTG
TTATTTATCAAGGCCAAGTTGTTGAGGATGTGACTACACACTTCTCTACGGCTGGAGG
CATATCGCAAGTTGCAGAAATCTACGAAGAACAATTCAGCGGTGAACACGACGACAT
TCTGACTTACGTACGCCCCGACGGTTACCTGGCCTCTTCTGCCTATATGCCCCCTAGAA
ACAAAGAAGACTTGGCGTCGCGACGCCGCGCAATCACGTACGTCTCGCAAAAAACC
TGGGGCACCCACTGCCGCAACCTGGACATGATCGCCAGCTTCACCGTCGGCATGATG
GGATATCGGCCGACATTCAGGAAAAACCCGGGACTGAG

**Figure S23:** FASTA formatted nucleotide sequence used as the gene synthesis fragment encoding the Δ*phz* deletion allele. Sequences spanning 700 bp upstream and 701 bp downstream of the fusion junction (where the pink region meets the yellow region) are included to target homologous recombination to the corresponding sequences in the *Pseudomonas chlororaphis* subsp. *aureofaciens* genome. The deletion replaces the *phzABCDEFG* operon with the deletion allele, in which the initial 30 bp of *phzA* (pink) are fused in-frame with the terminal 30 bp of *phzG* (yellow). Restriction sites for SacI (blue) and XmaI (green) were added exterior to the homologous region of the allele for downstream ligation. Additional 6 bp extension sequences (grey) were included to improve digestion efficiency.

```
>hcn_operon_with_flanking_regions_deleted_-_hcn_insert_-_buffer_sequence_added
Pseudomonas chlororaphis subsp. aureofaciens ATCC® 13985™, contig 1
ACTGAGGAGCTCGCCGGGCGCTCCAGGGGCGAAACCTTTGCCGCGTTTGCCGGCAG
CGGTATCAAGGAGCGGCTGCTGGCGTTCTGGATCGCCGGCTATTCCGACCCCAAGCT
CTATGCCCGGCACCGGCGCAAGATCCAGATCCTGCTGGCGGTGATGCTGTTGCAGGC
GGTGGTCGGCGCGGTGGCCGGGTTCTTCCTCGGCTCGGTCATAGGTCCGAAGGCTGC
ATGGGTATTCGGACTGTTGGGCGGGGTGGTGCCGCTGCTGTTCGCCTGGGGCTTCCAT
AAACCTTCGGTGAACGCCTACAGCCTCTATGCCCTTCTGACCCTGTCACAATTCCCCC
GATTGTTCGAAGGTTATGCCGAGGACCCGTTGGGTACCCTGATCGGCGCGGCCATGA
CCTTGGGGGTGGTGTTTTTCGCGATGTACGTGCGCAGCCTGATCTTTCCGGATTTCGC
TTTTATCGGGCCGCGCAAGGTCAAGGGTCGCTACGTGTTTTCCAGTTAGGGCAGGGC
GGCGCGAGGGCTGTTCCAGAGCCTTCCGCTTTGCTTGGCCCAGATCAATGTTCTTGTT
CGACAAACCTACTAGATTGGCCTGGTGTTCAGCTCCGAGGAGCTGGCGTCGCGCAAC
CCTGAACCGGGTCGAAGTCCAGGTTCAGGGTGATGCACAGCAGGACGCTGGCATTCC
TCTTCACTCAAGGATGAATTTACTGATGCGCCATACCGAACGAACTTTCGATATCGTG
GCGCCGTTGCAGGTGGCGCTGGCCTGAGCCTGGCGCGGTGCTTCAGAAGCACCGCG
TCGCCCTTGAGCCTTTTCGCGGGCAAGCCTCGCTCCTACAGATGCACGCATATCTTCT
GTAGGAGCGAGGCTTGTCCGCGATGGAGGTAAACGATAACGCGGTGTTACTGAAACA
CCGCGCCGTTTCGTGGCTTCAAACCATCGGTGTACTGACAAACAACTCCAACCCTTG
CGCATACTCCGCAAACGCCTTGACCCTTGGAAACCGCGTGGCATCCAGGTGCTCGGC
CACCACCAGCTGGGTAAAGCTCCAGGCCACGGCCAGGCTGATGCCGGCCTGATCGAT
GTTGCCGTCGGTGGCCAGCGGGCGTTTTTCCAGTTCGTTTTCCAGCTCGCCATAAGCC
GCCGCCAACTGCCCGCGAACCCGCTCCAGCCACGGCTCGTGCTGTTTTTCCGCAGGG
CGCAACTGGTACTCGTAGTAGATCTGCACCGACTTCTCGCATGCCGCCAGCGCCAGG
CCGATCAGCCGCAGGTCGTGGCCCGCGCCTGGGGCGCCGCCGGCAACAGGCTGCG
GCCGGCCAGGGTTTCCAGGTAGTCGATGATCAGGCACGAATCCATCAGCACCGTACC
GTCGTCGCAGATCAGGGTCGGCGCCTTGACCACCGGGTTGATGCGCTGGAACTGCCC
GGGACTGAG
```

**Figure S24:** FASTA formatted nucleotide sequence used as the gene synthesis fragment encoding the Δ*hcn* deletion allele. Sequences spanning 700 bp upstream and 654 bp downstream of the fusion junction (where the pink region meets the yellow region) are included to target homologous recombination to the corresponding sequences in the *Pseudomonas chlororaphis* subsp. *aureofaciens* genome. The deletion replaces the *hcnABC* operon with the deletion allele, in which the initial 30 bp of *hcnA* (pink) are fused in-frame with the terminal 30 bp of *hcnC* (yellow). Restriction sites for SacI (blue) and XmaI (green) were added exterior to the homologous region of the allele for downstream ligation. Additional 6 bp extension sequences (grey) were included to improve digestion efficiency.

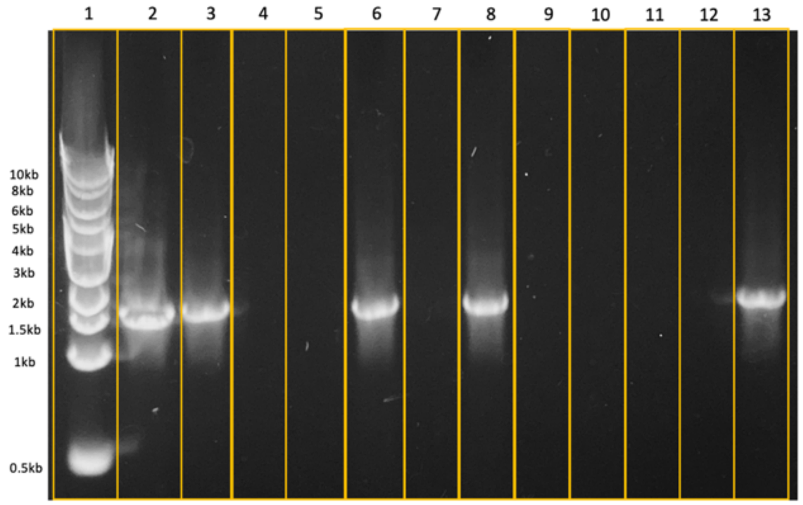


| Lane | Sample |
|---|---|
| 1 | Ladder |
| 2 | PC phz 1 |
| 3 | PC phz 2 |
| 4 | NC |
| 5 | NTC ($H_2O$) |
| 6 | phz 3.1 |
| 7 | phz 3.2 |
| 8 | phz 3.3 |
| 9 | phz 3.4 |
| 10 | phz 10.1 |
| 11 | phz 10.2 |
| 12 | phz 10.3 |
| 13 | phz 10.4 |

**Figure S25:** Colony PCR gel for identifying isolated SucR conjugates containing the Δ*phz* deletion allele. 1% agarose gel in 0.5X TAE Buffer supplemented with GelGreen dye was run at 90 V for 47 min. Ladder = 1 kb DNA Ladder. Positive control (PC) phz 1 = Δ*phz* insert DNA from Twist Bioscience. PC phz 2 = KanR *P. chlororaphis* subsp. *aureofaciens* Δ*phz*. Negative control (NC) = KanR *P. chlororaphis* subsp. *aureofaciens* without the Δ*phz* insert. No template control (NTC) = water added in place of template DNA. Successful deletion mutants can be seen in lanes 6, 8, and 13, based on amplification of Δ*phz* deletion allele with a size of ~1.4 kb.

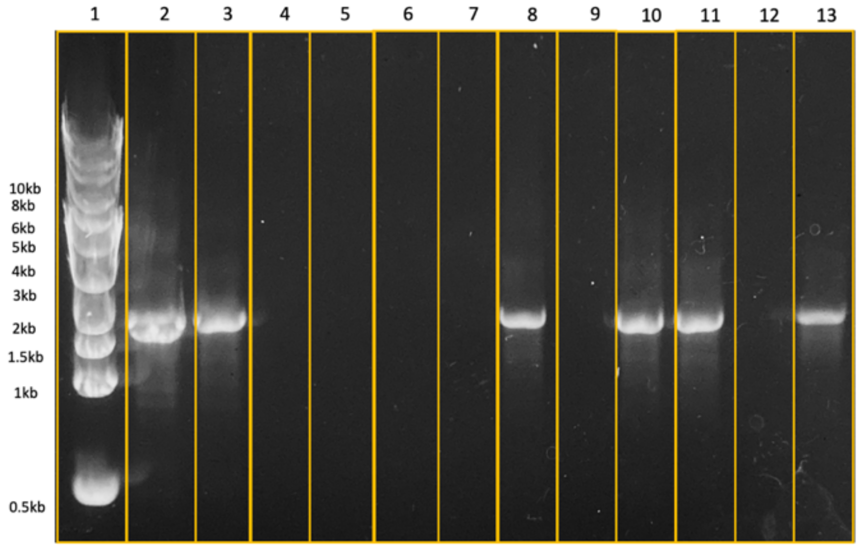


| Lane | Sample |
|---|---|
| 1 | Ladder |
| 2 | PC hcn 1 |
| 3 | PC hcn 2 |
| 4 | NC |
| 5 | NTC ($H_2O$) |
| 6 | hcn 10.1 |
| 7 | hcn 10.2 |
| 8 | hcn 10.3 |
| 9 | hcn 10.4 |
| 10 | hcn 14.1 |
| 11 | hcn 14.2 |
| 12 | hcn 14.3 |
| 13 | hcn 14.4 |

**Figure S26:** Colony PCR gel for identifying isolated SucR conjugates containing the Δ*hcn* deletion allele. 1% agarose gel in 0.5X TAE Buffer supplemented with GelGreen dye was run at 90V for 47min. Ladder = 1kb DNA Ladder. Positive control (PC) hcn 1 = Δ*hcn* insert DNA from Twist Bioscience. PC hcn 2 = KanR *P. chlororaphis* subsp. *aureofaciens* Δ*hcn*. Negative control (NC) = KanR *P. chlororaphis* subsp. *aureofaciens* without the Δ*hcn* insert. No template control (NTC) = water added in place of template DNA. Successful deletion mutants can be seen in lanes 8, 10, 11, and 13, based on amplification of Δ*hcn* deletion allele with a size of ~1.4kb.

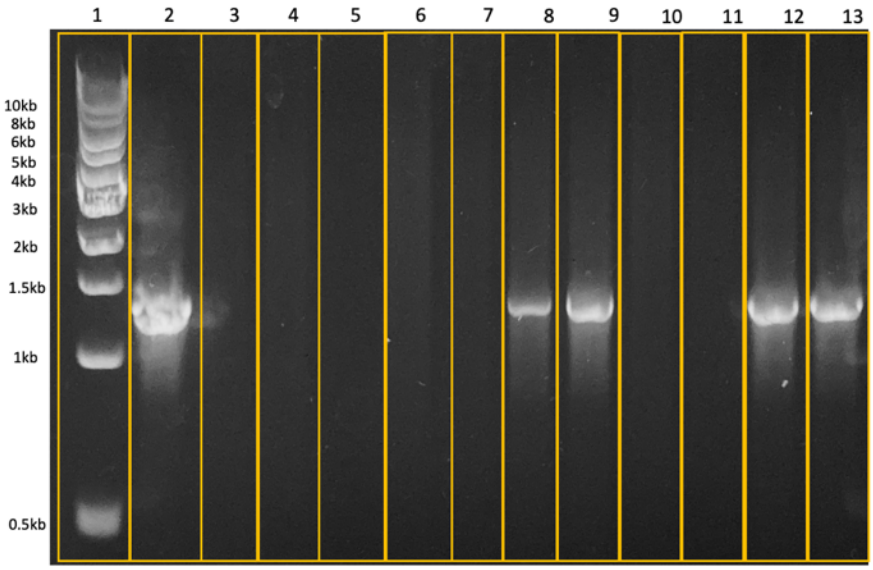


| Lane | Sample |
|---|---|
| 1 | Ladder |
| 2 | PC |
| 3 | NC 1 |
| 4 | NC 2 |
| 5 | NTC ($H_2O$) |
| 6 | phz 3.1 |
| 7 | phz 3.2 |
| 8 | phz 3.3 |
| 9 | phz 3.4 |
| 10 | phz 10.1 |
| 11 | phz 10.2 |
| 12 | phz 10.3 |
| 13 | phz 10.4 |

**Figure S27:** Colony PCR gel for identifying isolated Δ*hcn* SucR conjugates containing the Δ*phz* deletion allele. 1% agarose gel in 0.5X TAE Buffer supplemented with GelGreen dye was run at 90V for 45min. Ladder = 1kb DNA Ladder. Positive control (PC) = Δ*phz* insert DNA from Twist Bioscience. Negative control (NC) 1 = *P. chlororaphis* subsp. *aureofaciens* Δ*hcn* 14.1, without the Δ*phz* insert. NC 2 = WT *P. chlororaphis* subsp. *aureofaciens*. No template control (NTC) = water added in place of template DNA. Successful deletion mutants can be seen in lanes 8, 9, 12, and 13, based on amplification of Δ*phz* deletion allele with a size of ~1.4kb.

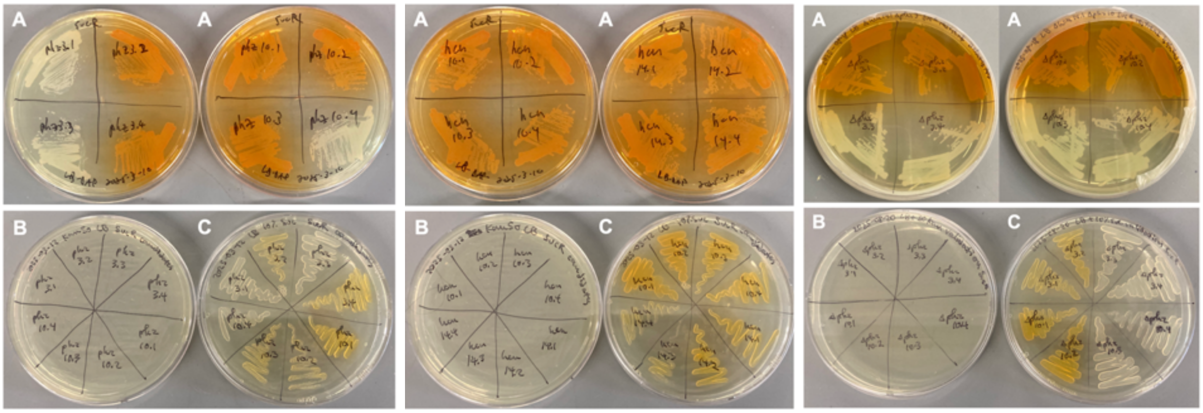


**Figure S28:** Phenotypic validation of SucR *P. chlororaphis* subsp. *aureofaciens* mutant candidates on LB (A), LB + 50 ug/mL Kan (B), and LB + 10% sucrose (C) plates. Deletion mutant candidates are shown for Δ*phz* (left), Δ*hcn* (middle), and Δ*phz*Δ*hcn* (right). Note that all mutants were capable of growth on plates containing sucrose, while Δ*phz* & Δ*phz*Δ*hcn* mutants did not produce the characteristic orange colour associated with phenazine-1-carboxylic acid (PCA).

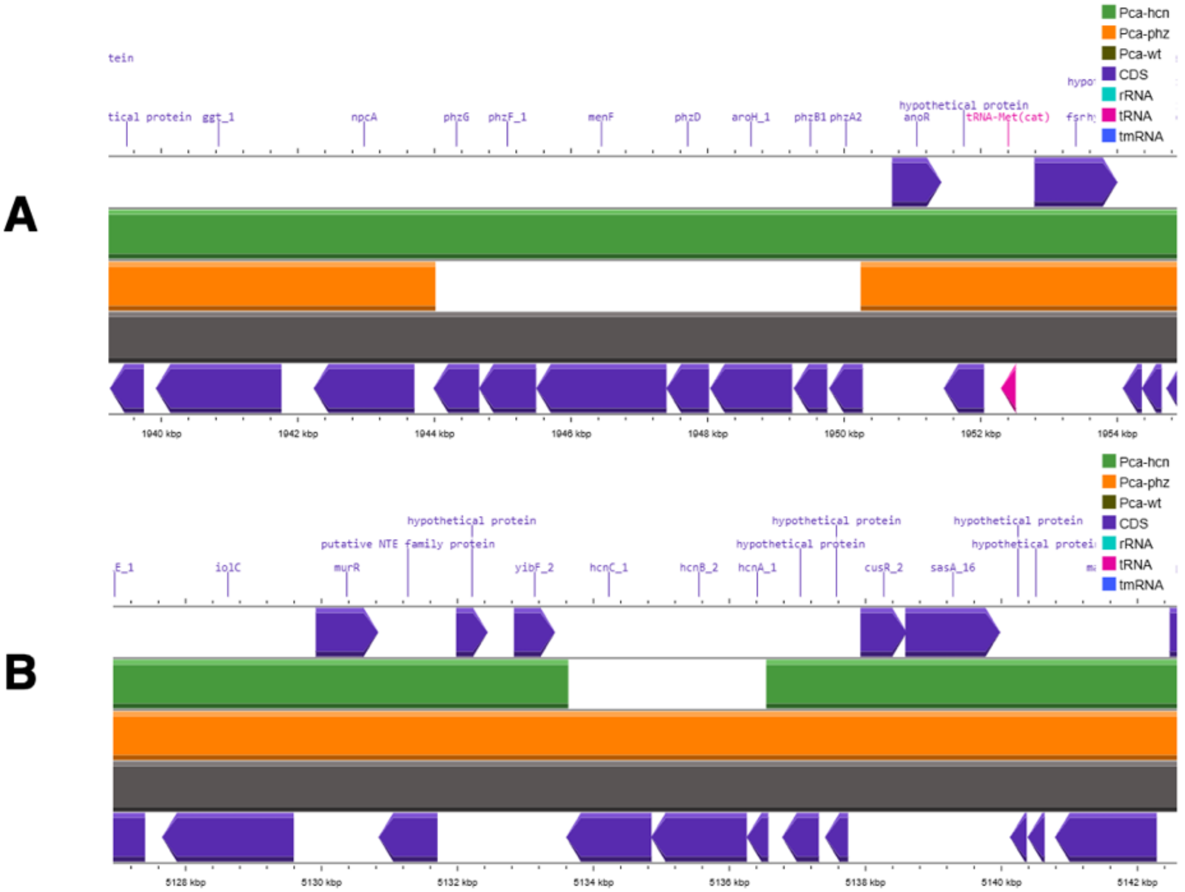


**Figure S29:** Genomic alignments of Δ*phz* and Δ*hcn* mutants in regions corresponding to the *phz* and *hcn* operons present in wild-type *P. chlororaphis* subsp. *aureofaciens*, based on whole-genome sequencing results from Flow Genomics. Visualization was done using Proksee with Prokka to provide genome annotations to the uploaded fasta sequences (**Methods**). A) Region of *phz* operon, showing the Δ*phz* deletion mutant (Pca-phz in orange), lacks the core genes for phenazine biosynthesis found in the wild type (Pca-wt in grey) and Δ*hcn* mutant (Pca-hcn in green). B) Region of *hcn* operon, showing the Δ*hcn* deletion mutant (green), lacks the genes encoding for critical proteins in cyanide production present in the wild type (grey) and Δ*phz* mutant (orange).

9) LC-HRMS QA/QC

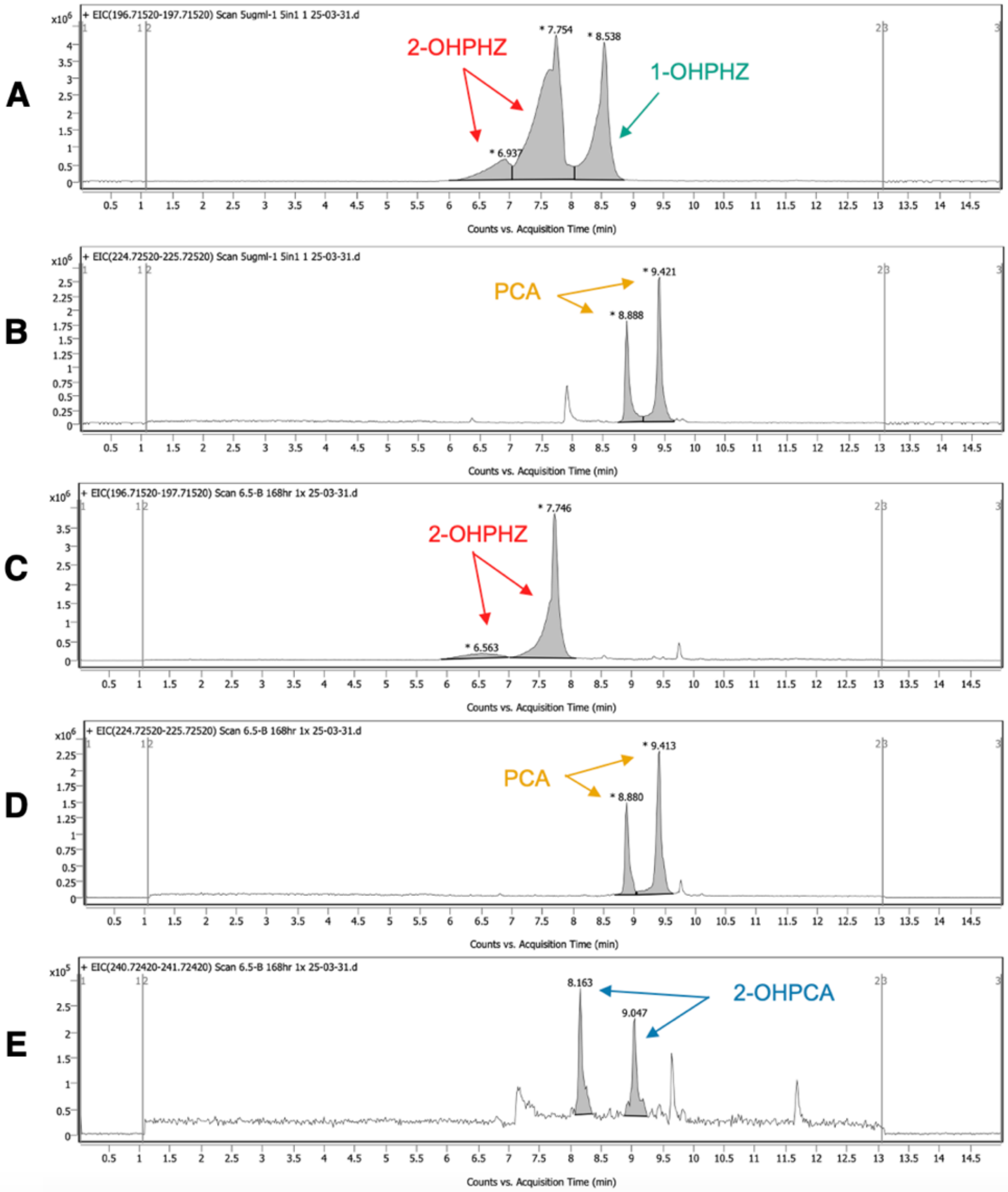


**Figure S30:** EICs of 5 µg/mL PCA and OHPHZ isomers in 5-in-1 standard (A,B), compared to peaks corresponding to 2-OHPHZ, PCA, and suspected 2-OHPCA in a randomly selected wild type sample at 168 hours (C-E).

**Table S7:** Summary of mass-to-charge ratios and retention times of all compounds in 5-in-1 standard solutions and 2-hydroxyphenazine-1-carboxylic acid (2-OHPCA) hypothesized to be present in wild type cell cultures.

| Metabolite | Mass-to-charge ratio (m/z) | Retention time (min) |
|---|---|---|
| PHE | 181.2162 | 9.049 |
| 1-OHPHZ | 197.2152 | 8.557 |
| 2-OHPHZ | 197.2152 | 6.942, 7.774 |
| PCN | 224.241 | 7.924, 8.559 |
| PCA | 225.2252 | 8.881, 9.414 |
| 2-OHPCA | 241.2242 | 8.158, 9.033 |

**Table S8:** Calibration curves of PCA and 2-OHPHZ of all three corrosion runs on LC-HRMS. Each curve spanned concentrations 0.1 to 7 µg/mL in triplicate, resulting in 27 total points at 9 calibration levels.

| Standard compound | Treatment of batch run | Slope (AU*mL/µg x $10^6$) | Intercept (AU x $10^6$) | Regression coefficient ($R^2$) |
|---|---|---|---|---|
| 2-OHPHZ | Wild type | 24.29 ± 0.31 | 3.97 ± 1.19 | 0.996 |
| | Spent medium | 32.86 ± 0.99 | 15.85 ± 3.78 | 0.978 |
| | Δ*phz* | 34.08 ± 1.21 | 17.78 ± 4.61 | 0.970 |
| PCA | Wild type | 4.13 ± 0.12 | 1.89 ± 0.47 | 0.979 |
| | Spent medium | 5.06 ± 0.31 | 6.27 ± 1.19 | 0.913 |
| | Δ*phz* | 4.43 ± 0.23 | 4.81 ± 0.87 | 0.938 |

10) Supporting References


1. Blin K, Shaw S, Vader L, Szenei J, Reitz ZL, Augustijn HE, Cediel-Becerra JDD, de Crécy-Lagard V, Koetsier RA, Williams SE, Cruz-Morales P, Wongwas S, Segurado Luchsinger AE, Biermann F, Korenskaia A, Zdouc MM, Meijer D, Terlouw BR, van der Hooft JJJ, Ziemert N, Helfrich EJN, Masschelein J, Corre C, Chevrette MG, van Wezel GP, Medema MH, Weber T. 2025. antiSMASH 8.0: extended gene cluster detection capabilities and analyses of chemistry, enzymology, and regulation. Nucleic Acids Research gkaf334.
2. Shaffer M, Borton MA, McGivern BB, Zayed AA, La Rosa SL, Solden LM, Liu P, Narrowe AB, Rodríguez-Ramos J, Bolduc B, Gazitúa MC, Daly RA, Smith GJ, Vik DR, Pope PB, Sullivan MB, Roux S, Wrighton KC. 2020. DRAM for distilling microbial metabolism to automate the curation of microbiome function. Nucleic Acids Research 48:8883–8900.
3. Sprouffske K, Wagner A. 2016. Growthcurver: an R package for obtaining interpretable metrics from microbial growth curves. BMC Bioinformatics 17:172.
4. Ritz C, Streibig JC. 2005. Bioassay Analysis Using R. Journal of Statistical Software 12:1–22.
5. Figurski DH, Helinski DR. 1979. Replication of an origin-containing derivative of plasmid RK2 dependent on a plasmid function provided in trans. Proc Natl Acad Sci U S A 76:1648–1652.
6. Hmelo LR, Borlee BR, Almblad H, Love ME, Randall TE, Tseng BS, Lin C, Irie Y, Storek KM, Yang JJ, Siehnel RJ, Howell PL, Singh PK, Tolker-Nielsen T, Parsek MR, Schweizer HP, Harrison JJ. 2015. Precision-engineering the *Pseudomonas aeruginosa* genome with two-step allelic exchange. Nature Protocols 10:1820.
7. Liu K, Hu H, Wang W, Zhang X. 2016. Genetic engineering of *Pseudomonas chlororaphis* GP72 for the enhanced production of 2-Hydroxyphenazine. Microbial Cell Factories 15:131.
